# A workflow for combined detection of protein interactions and cell types for translational studies

**DOI:** 10.64898/2026.05.19.725895

**Authors:** Lisa Okasmaa, Tony Ullman, Pranauti Panshikar, Rita Hutyra-Gram, David Krantz, Päivi Östling, Anders Ullén, Charlotte Stadler

**Affiliations:** Department of Protein Science, KTH Royal Institute of Technology, Stockholm, Sweden; Department of Pelvic Cancer, Genitourinary Oncology, Karolinska University Hospital, Stockholm, Sweden; Department of Clinical Pathology and Cytology, Karolinska University Hospital, Stockholm, Sweden; Department of Oncology and Pathology, Karolinska Institute, Stockholm, Sweden

**Keywords:** Spatial Proteomics, multiplexed immunofluorescence, in situ Proximity Ligation Assay

## Abstract

Multiplexed imaging approaches of various molecular modalities in tissues are becoming increasingly adopted in discovery and translational studies. For clinical implementation, novel instrumentation, complex analysis workflows and high costs per sample are bottlenecks that hinders broader introduction in the clinical setting. Here, we demonstrate a cost efficient integrated workflow that combines multiplexed immunofluorescence of a handful of protein markers, with *in situ* proximity ligation assay, to detect direct protein interactions between neighboring cells. As a proof of concept case of relevance for clinical adaptation, we target the major immunotherapy signalling axis of programmed death receptor 1 (PD-1) and its ligand PD-L1, to demonstrate the interaction between immune cells in germinal centers of tonsil tissue and in a tertiary lymphoid structure in bladder cancer tissue, respectively, from a patient treated with immunotherapy.

## 1. Introduction

Spatial proteomics, recognized as method of the year in 2024 [1], has advanced biomedical research by providing highly multiplexed, single-cell resolution images of protein expression in intact tissue sections that preserve spatial context and reveal cellular organization and functions [2]. Today, multiple commercial platforms are available, offering diverse strategies to perform low to high-plex protein imaging in cells and tissue sections [3]. Many of these approaches use fluorescence or sequencing as a readout, to allow for the high number of protein target signals to be separated and analysed. Further, large scale multiomics platforms that target hundreds or thousands of proteins and transcripts in the same tissue section are commercially available [3,4]. Together these combined modalities bring unprecedented information with potential to deepen our understanding of the tissue microenvironment in both health and disease conditions. The opportunities for analysing large numbers of individual proteins and transcripts, can further help to discover subtle differences across cell types and functional states.

In a sophisticated study by Aung et al. they demonstrated that immunotherapy responses are highly variable, highlighting the need for other biomarkers than the currently used PD-L1. In this study based on non-small cell lung cancer (NSCLC) patients treated with immunotherapy, only a subset of patients responded, and approximately 40% showed resistance despite high PD-L1 expression. Using spatial proteomics and transcriptomics on 234 patients (67 and 131, respectively), they identified signatures correlating to immunotherapy outcome involving multiple cell types, not just those expressing PD-L1. Proliferating tumor cells, granulocytes, vessels and macrophages present in the tumor region were associated with increased risk for progression, while a response signature included presence of M1/M2 macrophages and CD4 T cells. Their findings emphasize the importance of exploring the spatial tumor microenvironment and that a deeper analysis that better captures the complex tumor landscape provides better guidance for predicting response to immunotherapy compared to current single biomarkers [5].

Despite its technical advances, increasing use in research and recognition for spatial omics being the next era in histopathology and diagnostics [6,7], the translation of multiplexed imaging methods into the clinical setting is so far limited. Some of the main challenges for clinical translation and implementation are infrastructure challenges including new instrumentation and analysis workflows and lack of experience from analyzing complex image data not based on chromogenic stains. Further, the need for standardized and reproducible protocols with defined readouts for a given assay are crucial in a clinical setting, as they often guide treatment decisions. Last, the costs associated with implementation of new methods and the price tag for performing multiplexed analysis per sample is high compared to current set-up.

The cost for highly multiplexed imaging methods are often scaled with plexity, meaning the more markers stained for, the higher cost for the assay. Many of the available commercial platforms for obtaining higher plex numbers, rely on end to end solutions with combined sample preparation, staining and data acquisition on their instrument using proprietary reagents. While pushing the multiplexing capacity to hundreds or even thousands of protein markers is desirable for discovery research, this is not currently feasible in a clinical setting. Instead, multiplexing methods that allow for simultaneous detection of a handful of markers could be of high value, compared to the traditionally used histopathology methods like hematoxylin and eosin stain, or single plex immunohistochemistry (IHC). For example, primary diagnosis of lung cancer can require staining of several protein markers such as TTF-1, Napsin A, p40 and p63 to accurately define the tumor subtype [8]. Another example would be cytokeratin profiling that can help narrow down the site of origin in carcinomas of unknown site (CUPs) [9]. Using a multiplexed method rather than traditional single plex IHC, staining for these respective markers could be done in one experiment and a single tissue section, instead of using one section per marker. In the era of personalized medicine, panels of protein markers targeted by new therapies, such as for antibody-drug conjugates (ADC) targets could be combined to help in treatment decisions where several potential therapies are at the table. Overall, incorporating a limited number of clinically relevant biomarkers into diagnostic workflows could improve disease characterization and treatment selection. In addition, sample material can be minimized and secure more information even from cases with limited material.

While staining of certain protein markers is currently the main application for clinical histopathology, the methods for detecting relevant markers is not limited to protein expression such as Human Epidermal Growth Factor Receptor 2 (HER 2), cytokeratins and novel therapeutic targets. Instead, the opportunities with multiplexing allow for simultaneous detection of several markers across modalities. With appropriate assays and staining protocols, protein detection can be combined with RNA or DNA targets, or reveal more functional states by looking at protein interactions. For example, protein interaction has been shown to be more predictive to certain therapy responses as signalling events between cells can be detected *in situ*. As an example, the programmed cell death receptor 1 (PD-1) and its ligand programmed death ligand 1 (PD-L1) are major targets in cancer immunotherapy. This axis mediates immune tolerance by suppressing excessive T-cell activation and signalling normally occurring between immune cells [10]. Treatment decisions for immune checkpoint inhibitors targeting this signaling axis, often rely on PD-L1 expression assessed by IHC staining, however, its predictive power for response is limited [11–13]. In a recent study of non-small cell lung cancer (NSCLC), authors explored the predictive value of the presence of direct interaction between PD-1 and PD-L1 and found this assay to outperform standard PD-L1 staining as a response predictive marker [14]. However, this approach did not distinguish between different cell types in the samples, and thus not whether the PD-1/PD-L1 interaction between immune cells or between cancer and immune cells were detected.

Based on the promise for detection of protein interactions in response prediction, we developed a protocol for an automated low-plex immunofluorescence panel, integrated with *in situ* Proximity Ligation Assay (isPLA), for simultaneous detection of protein-protein interaction and cell phenotypes in the same tissue section. In contrast to the abovementioned study, such a combined workflow can distinguish between immune cell interactions and cancer-immune cell interactions and allow for more in depth exploration of protein interactions as a biomarker. Further, the workflow is cost efficient relative to highly multiplexed assays, and can be adapted to target relevant interactions and cell subtypes based on clinical needs.

## 2. Results

### 2.1 An automated immunofluorescence workflow with tyramide signal amplification for tissue

In this study, a 5-plex immunofluorescence workflow was automated for tissue staining, utilizing poly-horseradish peroxidase (HRP)-mediated tyramide signal amplification (TSA). The assay relies on antibody-based detection, by which primary antibodies recognize target proteins, and poly-HRP-conjugated secondary antibodies catalyze fluorescent deposition at tyrosine residues around protein epitopes, producing a covalently bound and amplified signal. Intervening rounds of heat-induced epitope retrieval (HIER) enables cycling of several markers. For this study, focusing on a low-plex immunofluorescence panel for a limited set of samples, the autostainer Spatial Station from Parhelia Biosciences was used for automation.

The workflow was separated into three main stages: Step 1. Manual tissue preparation, Step 2. Tissue staining and Step 3. Image acquisition, as shown in **Figure 1**. Given the availability of an already existing manual tissue preparation protocol, this step remained unchanged, with efforts focused on automating the staining process (Step 2). The automated workflow reduced hands-on time to approximately 1 hour for the staining step **(Figure 1B)**, in comparison to a manual staining workflow, included as a reference, which required up to 20.5 hours distributed over several days **(Figure 1A)**. Overall, the automated workflow represents an estimated reduction of 16.5 hours in active hands-on time per tissue and limits manual work to a single day.

**Figure 1.**
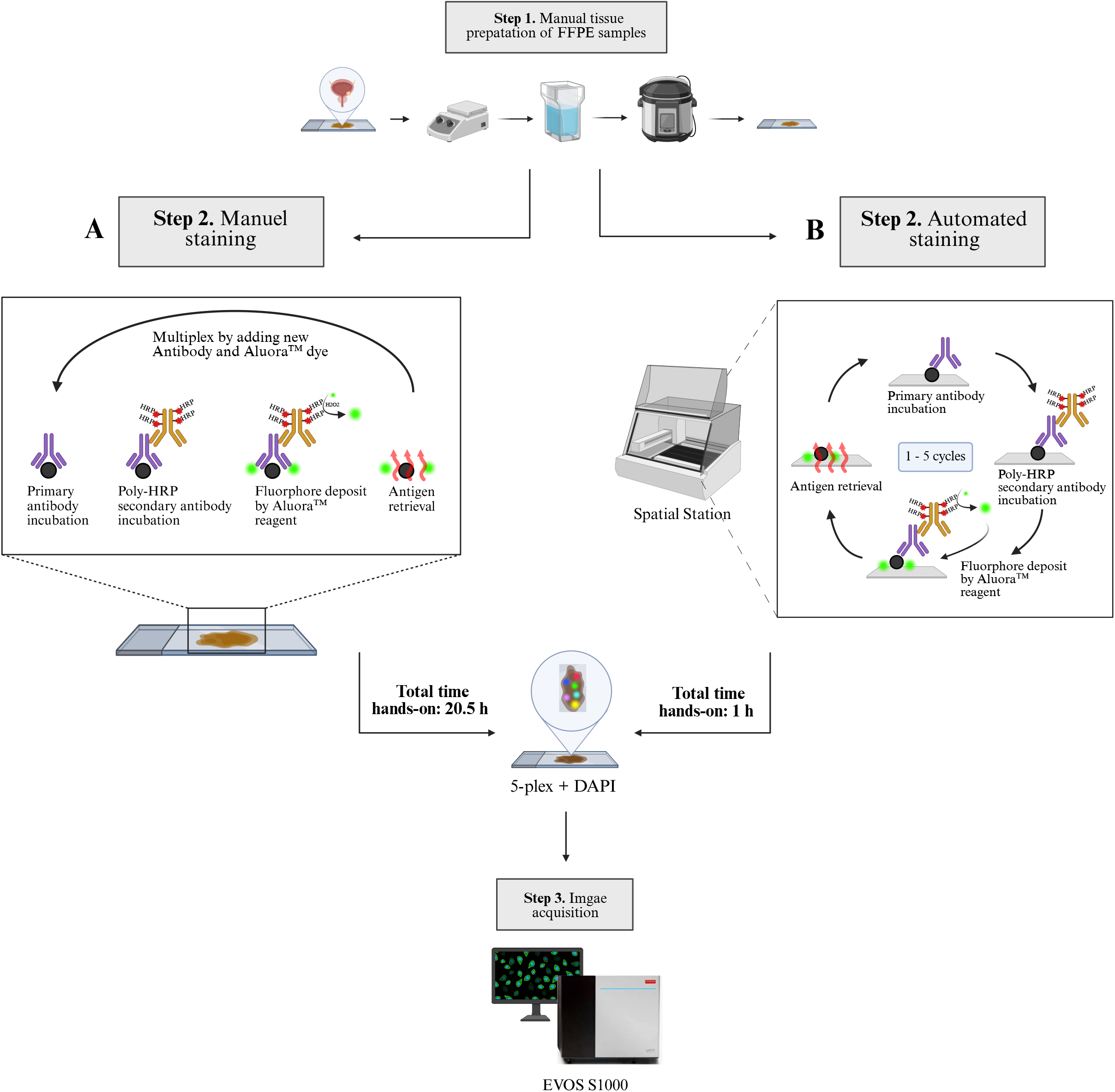

#### 2.1.1 Validation of the automated immunofluorescence workflow on human tissue

Once the workflow was established, we applied it to formalin-fixed paraffin-embedded (FFPE) tumor tissue samples derived from human bladder cancer. The marker panel included three membranous immune cell markers (CD45, CD3, CD8a) and two membranous epithelial markers (Pancytokeratin “PanCK”, Nectin-4), providing a representative overview of the tissue. The marker panel, alongside corresponding fluorescent dyes and cycle order, is summarized under the method section. All protein markers were successfully detected, as illustrated in **Figure 2A**, demonstrating that the protocol can be automated for amplified detection of a limited set of markers for cell phenotyping in tissue. Despite repeated exposure of samples to HIER, tissue integrity and protein epitopes remained intact across all cycles.

**Figure 2.**
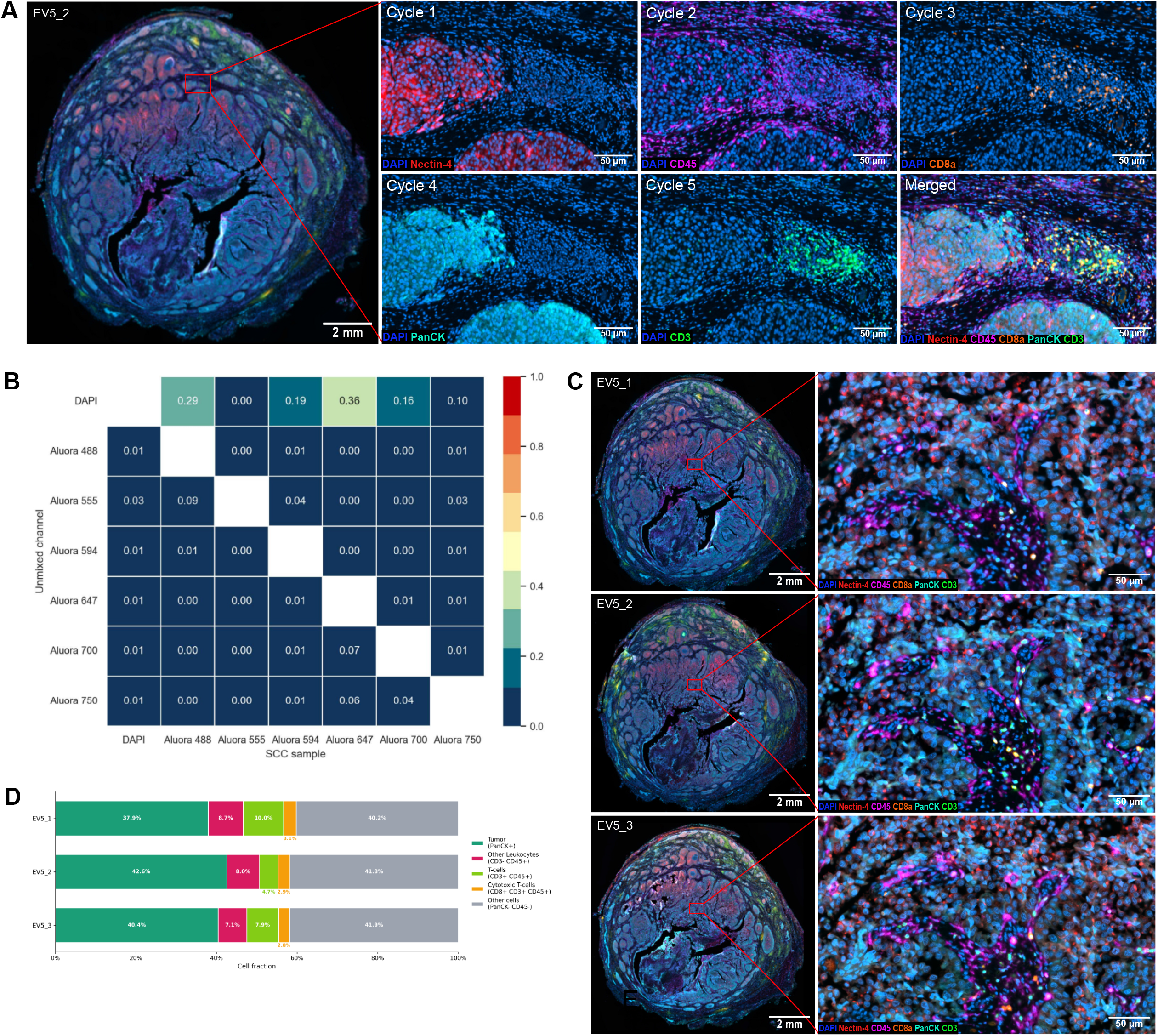

Visualization of five protein markers was enabled by spectral unmixing using the EVOS s1000 Spatial Imaging System, which separates overlapping fluorescent signals during image acquisition. Unmixing quality can be quantitatively assessed via Fraction Bleedthrough (FB), the ratio of residual signal in the off-target channels compared to signal intensity of fluorophores in their primary channels post-unmixing. The imaging software provides a FB heatmap **(Figure 2B)**, with values < 0.25 indicating high quality unmixing. All markers had acceptable FB values (< 0.25), except moderate DAPI spillover in the 488 and 647 channel, however, the DAPI bleedthrough did not comprise overall panel performance.

To assess reproducibility in the automated workflow, we stained three consecutive sections (EV5_1, EV5_2 and EV5_3) from the same bladder cancer tumor tissue block. The three sections were processed in parallel and showed consistent staining intensity and spatial distribution **(Figure 2C)**. To more quantitatively evaluate the reproducibility of biologically meaningful results of the automated workflow across the consecutive bladder cancer sections, we compared the major cell-phenotype fractions obtained from image analysis of the serial EV5 tissue sections **(Figure 2D)**. Here, Nectin-4 expression was excluded from the quantitative analysis, as it does not define a distinct cellular population and was observed to be co-localizised with PanCK+ tumor cells. The overall cellular composition was highly similar between sections. PanCK+ tumor cells accounted for 37.9–42.6% of all segmented cells, Other cells (PanCK-C45-) for 40.2–41.9%, and other leukocytes (CD3-CD45+) for 7.1–8.7%. Cytotoxic T cells (CD8+ CD3+ CD45+) remained consistently low across sections, varying only from 2.8% to 3.1%. The greatest relative variation was observed in the T-cell (CD3+ CD45+) compartment, which ranged from 4.7% to 10.0%. Overall, these findings indicate that the major tissue compartments were stably recovered across adjacent sections, with somewhat greater fluctuation confined to smaller immune subsets. While all samples analysed came from the same tissue specimen, subtle differences in cell type distribution are expected due to being from different sections.

### 2.2 Combining and automating an immunofluorescence and in situ Proximity Ligation Assay on tissue

While an immunofluorescence assay can visualize protein expressions and their localization in tissue, it provides limited insight of cellular interactions and protein functional states. We therefore incorporated isPLA into the workflow, to enable simultaneous detection of protein expression and protein-protein interactions in the same tissue section. As proof of concept, we examined the PD-1/PD-L1 axis, given its clinical relevance in cancer.

The isPLA was integrated as the first staining cycle of the automated workflow to detect interactions between target proteins (PD-1/PD-L1) when in close proximity (≤ 40 nm) [15]. Primary antibodies recognize the target proteins, followed by incubation with probe-conjugated secondary antibodies, probe ligation and rolling circular amplification (RCA), to allow hybridization of complementary HRP-labeled oligonucleotides. Instead of adding the chromogenic detection provided by the kit manufacturer (Navinci Diagnostics), we applied remaining Aluroa™ dye in the 647 channel for amplified detection, utilizing the HRP present in the Rolling circular product (RCP) to enable protocol integration. The automated immunofluorescence workflow could then proceed as before, as shown in **Figure 3**, staining protein markers in cycle 2-6.

**Figure 3.**
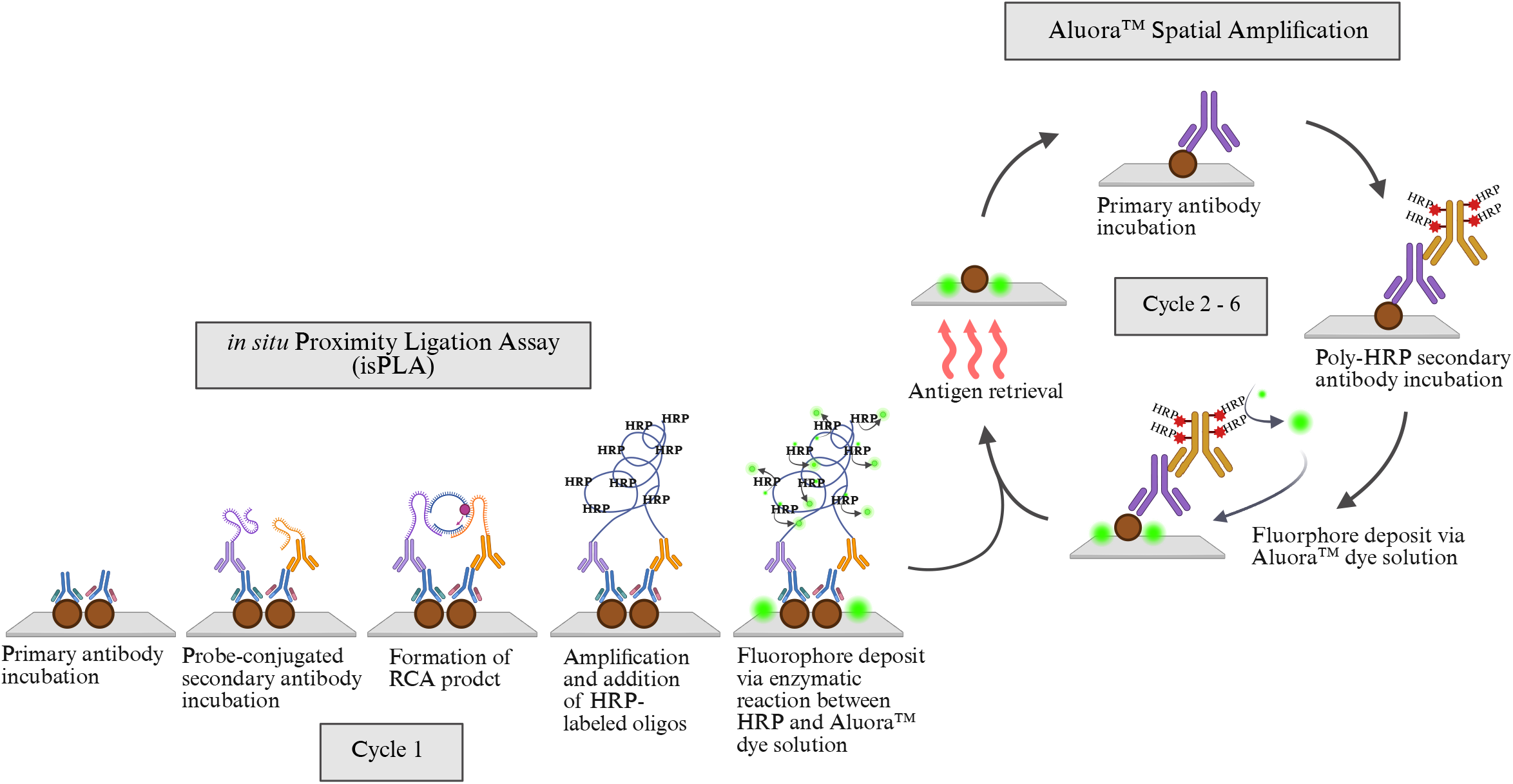

#### 2.2.1 Validation of the combined and automated immunofluorescence and in situ Proximity Ligation Assay workflow on human tissue

We began by evaluating the automated isPLA workflow in human tonsil FFPE tissue, a biological positive control with known PD-1/PD-L1 interactions, predominantly localized to germinal centers, and thus selected for initial validation of the assay. isPLA signal was detected in the Aluora 647 channel with clear fluorescent dots marking the interaction within several of the germinal centers across the tonsil tissue **(Figure 4A)**. The initial isPLA signal persisted throughout all staining cycles and immune cell markers (CD45, CD8a, CD3) had clear, detectable signals. In contrast, markers not typically expressed in tonsil tissue (PanCK, Nectin-4) showed no specific staining.

**Figure 4.**
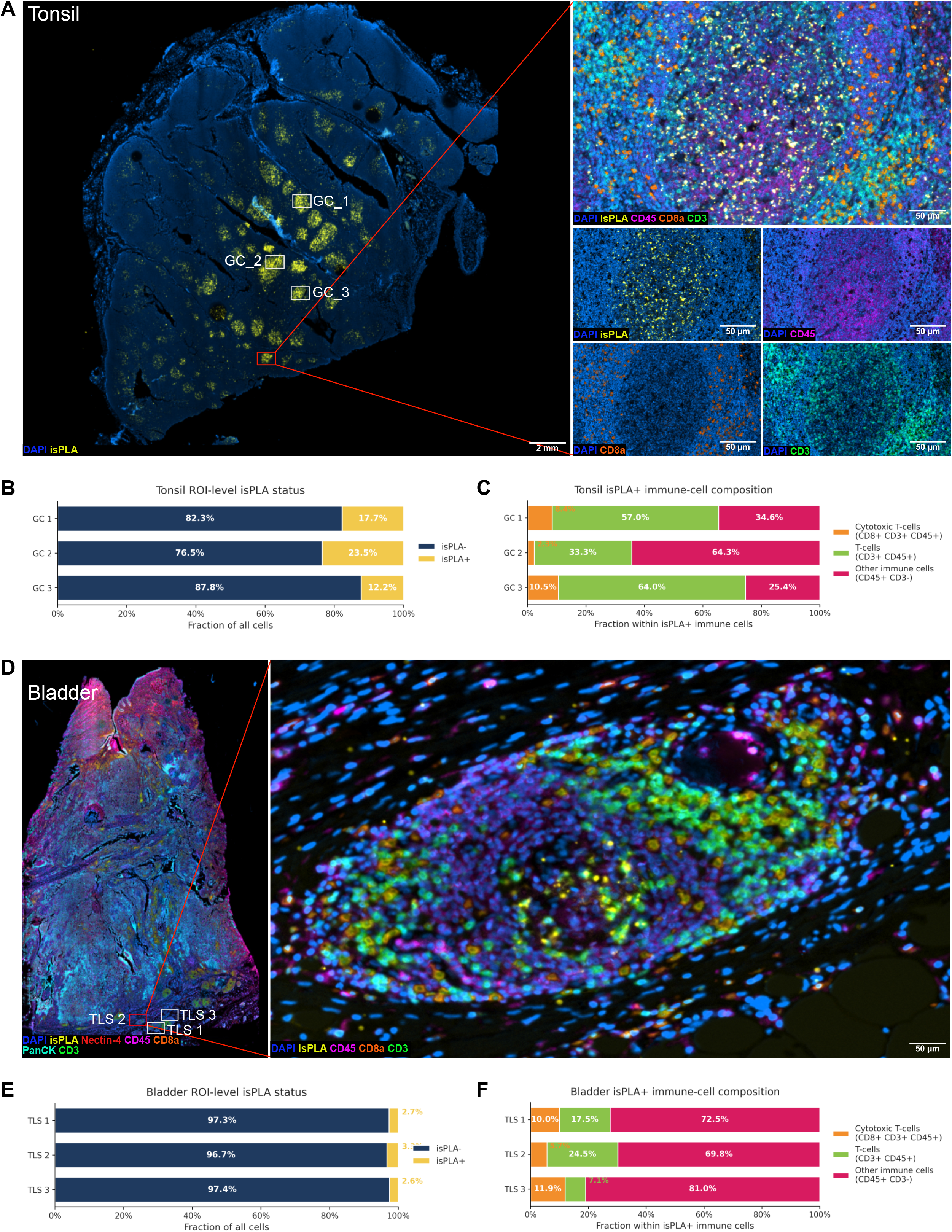

We next analyzed a few germinal centers to characterize the extent and distribution of PD-1/PD-L1 isPLA signal. In total, 29,589 annotated cells were identified across the three analyzed germinal centers (GC 1-3), including 9,895 cells in GC 1, 9,541 cells in GC 2, and 10,153 cells in GC 3. Of these, 5,242 cells (17.7%) were classified as isPLA+. The isPLA+ fraction of cells ranged from 12.2% to 23.5% across the analysed GC **(Figure. 4B)**. The relatively high abundance of isPLA+ cells observed in tonsil germinal centers is consistent with established germinal center biology, as these structures are known to contain abundant PD-1 expressing T-cell populations [16]. Correspondingly, PD-L1 is expressed by multiple antigen-presenting immune cell populations, such as B cells and myeloid cells, where PD-1/PD-L1 signaling plays an important role in regulating local immune responses [17].

As our workflow combines the isPLA assay with immunofluorescence of a few markers, the interactions between PD-1/PD-L1 could be annotated to defined cell types based on our markers in the panel. As the GC only contains immune cells, we explored the distribution of isPLA+ cells across the immune cell subtypes. As we only had markers to define T cells (CD3+) and the cytotoxic T cells (CD3+ CD8+), the remaining immune cells were annotated as “other” with the definition of CD45+CD3-CD8-. Among the isPLA+ immune cell types, T cells (CD3+) constituted a major component, although with variations across the analysed GC ranging from 33.3% to 64.0%, while other immune cells ranged from 25.4% to 64.3% and cytotoxic T cells from 2.3% to 10.5% (**Figure. 4C**). Combining all cells from the 3 GC, 5,215 isPLA+ immune cells were identified, including 2,530 T cells (48.5%), 2,356 other immune cells (45.2%), and 329 cytotoxic T cells (6.3%). These data indicates that PD-1/PD-L1 interactions occur across different immune cell types in GCs, with a substantial contribution from T-cell-rich regions (**Figure S1A**). Following successful validation in human FFPE tonsil tissue, we appl ied the integrated workflow to FFPE bladder cancer tumor tissue samples, for which the marker panel was originally designed. The complete workflow showed detection of isPLA signals alongside protein marker staining **(Figure 4 D)**. Notably, the isPLA signal was confined to regions identified as tertiary lymphoid structures (TLS) by immune cell markers, structures prone to interactions since TLS mediates anti-tumor response [18]. To verify specific isPLA signal, a technical negative control was included from the same FFPE bladder cancer block, processed through the full isPLA workflow but without primary antibodies. No detectable isPLA signal was observed in the technical negative control (data not shown).

We next performed the same analysis as in the tonsil section, here within a few selected TLS. In total, 10,559 annotated cells were identified across the three analyzed bladder ROIs, including 4,576 cells in ROI 1, 2,723 cells in ROI 2, and 3,260 cells in ROI 3. Of these, 300 cells (2.8%) were classified as isPLA+. In contrast to tonsil, the proportion of isPLA+ cells was uniformly low across the analysed TLS, corresponding to 124 isPLA+ cells in TLS 1, 91 in TLS 2, and 85 in TLS 3 **(Figure. 4E)**. Also for the TLS regions in the bladder section, the composition of different immune cells within the isPLA+ immune-cell fraction was analysed. isPLA+ immune cells were predominantly assigned to the “other” immune-cell category, accounting for 69.8–81.0% across ROIs, whereas T-cell and cytotoxic T-cell fractions remained comparatively small **(Figure. 4F)**. In total across the analysed TLS, 135 isPLA+ immune cells were identified in bladder TLSs, of which 100 (74.1%) were classified as “other” immune cells, 23 (17.0%) as T cells, and 12 (8.9%) as cytotoxic T cells (**Figure S1B**).

While the markers in this panel are limited and not selected to explore interactions between different immune subsets that are expected to be present in both GC and TLS, they demonstrate that the automated and integrated workflow enables detection of protein-protein interaction between defined cell types in a single tissue section. No modifications were required when applying the automated workflow to a different type of tissue, however, staining performance is marker dependent and may vary with other protein panels.

## 3. Discussion

In this work, we developed an automated protocol for low plex IF alone, and a protocol for the combined detection of low plex IF and cell interactions using isPLA in a single tissue section. While the combination of multiplexed IF and interaction assays such as isPLA has been demonstrated previously [19], this workflow is focused on common needs for translational studies, with detection of relatively few protein markers at reasonable cost, to allow for higher sample numbers to be analysed. Further, the protocols are automated with little hands-on time, allowing for completion within one day of up to 12 slides, compared to three days using a manual workflow. Conceptually, the number of slides prepared simultaneously depends on what autostainer is used, here being limited to 12 on the current set-up of the instrument. It should be noted that our integrated protocol includes steps performed at several different temperatures, 36 - 38°C for enzymatic reactions step for the isPLA part, up to 90°C for elution steps of antibodies at the end of every staining cycle and with antibody incubation steps ranging from room temperature to 38°C, meaning not only different temperatures are used but they are also altered back and forth during the protocol. These specific temperature ranges are needed for successful assay results, and could limit the options of suitable autostainers to perform the assay, depending on its heating and cooling capacities. An option could be to split the protocols in two, performing the isPLA part separately, followed by the multiplexed IF (mIF) part to avoid the large changes in temperature. While this would make the workflow not fully automated, the results with integrated isPLA and mIF readout in a single scanning experiment would remain. Also in our protocol set-up on Spatial Station, it should be noted that while all slides remain on the instrument from first to last step of our assay, the isPLA and mIF protocols are separated and thus require starting the second protocol after completion of the first. This is due to a current limitation in the number of steps that can be programmed for a single protocol on the autostainer and will likely be solved with future upgrades. In the end, the intervention with hands-on time to start the next protocol is limited to 10 minutes.

A strong argument for automating these protocols is the tedious process for generating mIF image data with amplified readouts such as TSA. Even though these protocols are cumbersome relative to workflows using directly labeled antibodies, the amplified methods have the advantage of providing better signal to noise ratios. In turn, this makes quantitative downstream analysis easier and more robust, even in samples with high endogenous fluorescence. Ideally, a panel of protein markers should be transferable from one tissue type to another, a transfer more easily introduced with high signal to noise ratios. In this work, we developed a 5-plex IF panel in bladder cancer tissue using the TSA based protocol, and then successfully transferred that panel to the tonsil section without need for any adaptation for the markers expected to be present in both tissue types (CD45, CD3 and CD8a). This is beneficial for translational studies and future clinical implementation, as many protein markers would overlap between mIF panels across cancer indications. For example, protein markers used to define presence of different immune cells types are relevant across tumor types, as well as structural proteins of tissue architecture or markers for certain cellular functional states such as ki67, a marker for proliferation. Even therapeutic targets can be tumor agnostic to some extent, like PD-1 or PD-L1 and ADC targets.

While amplified signals can simplify downstream analysis and cross tissue adaptation, the signal from TSA based readout has to be balanced to avoid oversaturation that can reduce dynamic range that challenge comparison of protein expression levels, or disguise more precise location of key protein markers of interest. An oversaturated TSA based readout can make protein staining appear as blobs, taking away the opportunities to reveal subcellular differences such as membrane versus cytosolic expression.

The concept of integrating a low plex IF and isPLA, could be adapted to use directly fluorescently labeled antibodies and a version of isPLA detection using a fluorescently labeled detection oligo that hybridizes directly to the RCP product. This workflow would be faster and allow for all antibodies of the mIF panel to be incubated in one single step, without the need for temperature ranges necessary for the amplified signal readouts. As for the amplified TSA based system demonstrated here, the number of antibodies and thus protein markers are limited by the unique fluorofores and the imaging instrument’s ability to separate the signals via spectral unmixing. However, we believe such set-up to be more context dependent, as signals would be weaker and perhaps not sufficiently strong for many protein targets in a tissue setting and with instrumentation often used to scan tissue sections, even if antibodies are well validated. On the other hand, a protocol without signal amplification and thus without repetitive steps at high temperature to elute antibodies as needed for the TSA based protocols, could improve tissue integrity and be needed for more fragile tissue specimens. While FFPE samples are standard in clinical routine and used for most translational studies, providing alternatives for fresh frozen or fixed frozen tissue sections could make the integrated protocol useful for a broader range of samples and research questions.

To conclude, we have developed an automated workflow for the combined detection of multiple protein markers and with an option to include a protein-protein interaction using isPLA. The workflow demonstrates a significant reduction in hands-on time and robust performance both between technical replicates and different tissue types. Last, the cost per tissue slide is limited to a few hundred dollars. Together, the workflow fulfills the main criteria to be suitable for large scale translational studies, where protein markers and the protein interaction can easily be adapted to fit the clinical question.

## Materials and Method

### Biological materials

Formalin-fixed paraffin-embedded (FFPE) bladder cancer tumor tissue included anonymized residual material retrieved from the pathology archive. Additional tumor tissue was obtained from a separate patient through routine clinical diagnostics and included under a parallel ethically approved study. All research performed on the material was conducted within the scope of the approved ethical permit (Swedish Ethical Review Authority, approval no. 2024-02569-01).

FFPE human tonsil tissue was provided by Navinci Diagnostics, who acquired the tissue from Acepix Biosciences, later sectioned at Rudbeck Laboratory (Uppsala). All tissue samples were anonymized and approved according to ethical guidelines and Swedish law.

### Tissue sample preparation

FFPE tissue samples were incubated on a hot plate at 57 °C for 30 min, followed by dewaxing and rehydration through sequential washes in the following solvents: 100% Histoclear (Merck, Cat# H2779-1L) for 2 × 10 min, 100% Ethanol (Sigma-Aldrich, Cat# 8187601000) for 2 × 5 min, 90% Ethanol (Sigma-Aldrich, Cat# 8187601000) for 5 min, 70% Ethanol (VWR, Cat# 8301.360) for 5 min, 50% Ethanol (Sigma-Aldrich, Cat# 8187601000) for 5 min, 30% Ethanol (Sigma-Aldrich, Cat# 8187601000) for 5 min and ddH_2_O for 2 × 5 min. Antigen retrieval was performed through high-pressure heat exposure in a pressure cooker at 114-121°C for 9 min in 1X Tris-EDTA buffer (pH 9) (Abcam, Cat# ab93684). Afterwards, samples were cooled to room temperature and washed twice in MilliQ water.

### Protocol automation on the autostainer Spatial Station

The autostainer Parhelia Spatial Station (PSS) from Parhelia Bioscience was used to automate the workflow. The system operates with two independent pipetting arms for liquid handling and can process up to twelve slides per run. It has controlled cooling and heating, which is required for the temperature-dependent enzymatic reactions and staining cycle separations for this workflow. On the platform, protocols are made manually by programming reagent steps together with their associated modules and plates. All liquid handling was performed with 300 µL CO-RE pipette tips (Hamilton, Cat# 235902), staining samples through a capillary laminar flow.

Prior to initiating the automated protocol, samples were mounted onto Chamfered Covering Pads (Parhelia Biosciences, Cat#40221) with 1XPBS and loaded into the ST-12 module (Parhelia Bioscience) together with superheating fluid (Parhelia Biosciences, Cat# 50005), which worked as an incubation chamber and maintained controlled temperatures.

### Implementation of the immunofluorescence protocol on Spatial Station

The staining workflow was initiated immediately after tissue preparation by placing samples in the PSS platform. Reagents were kept in a deep 96-well plate (Parhelia Biosciences, Cat#40207), while buffers, Wash buffer (1X PBS) and HIER buffer (1X Tris-EDTA buffer, pH 9, Abcam, Cat# ab93684), were kept in two deep 24-well plates (Parhelia Biosciences, Cat#40604). The automated protocol was performed according to the manufacturer’s instructions for the Aluora™ Spatial Amplification kit (Thermo Fisher Scientific, Cat# AS100HRP) unless otherwise specified. To adapt the protocol for automated processing, minor modifications were implemented as specified below.

### Modifications to the immunofluorescence protocol for automated implementation

Prior to the first staining cycle, samples underwent a 30 min incubation with 3% hydrogen peroxide (Thermo Fisher Scientific, Cat# AS100HRP). Primary antibodies were incubated for 60 min at a specified dilution **(Table 1)**, while HRP-conjugated secondary antibodies (Thermo Fisher Scientific, Cat# AS100HRP) were applied for 45 min. Between staining cycles, HIER with 1X Tris-EDTA buffer (pH 9) (Abcam, Cat# ab93684) was performed for 45 min at 90°C. After the automated protocol, samples were incubated at room temperature for 4 min with DAPI (1:5000) (Thermo Fisher Scientific, Cat# H3570), followed by three 1X PBS washes. A more detailed description of protocol parameters and deck setup can be found in the attached PSS run report **(Figure S2)**.

**Table 1.** Cycle order, protein markers and antibodies with corresponding Aluora™ dyes used in the automated mIF workflow.

| Cycle order | Marker | Supplier | Lot no. | Clone | Catalog no. | Aluora<sup>T</sup><br>M dye | Ex/Em | Emission filters | Dilution | Exposure time (ms) |
| --- | --- | --- | --- | --- | --- | --- | --- | --- | --- | --- |
| 1 | Nectin-4 | Abcam | 109814<br>8-11 | EPR156<br>13-68 | AB251<br>110 | Aluora<br>594 | 589/61<br>5 | Texas<br>Red | 1:300 | 50 |
| 2 | CD45 | Cell Signaling Technology | 10 | D9M8I | 13917 | Aluora<br>700 | 687/70<br>6 | Cy5.5 | 1:300 | 15 |
| 3 | CD8a | Thermo<br>Fisher<br>Scientific | 317527<br>2 | C8/144<br>B | MA5-13473 | Aluora<br>555 | 552/56<br>7 | RPF | 1:1000 | 20 |
| 4 | PanCK | Abcam | GR3418<br>528-2 | AE1/A<br>E3 | ab803<br>26 | Aluora<br>750 | 757/78<br>3 | Cy7 | 1:300 | 150 |
| 5 | CD3 | Thermo<br>Fisher<br>Scientific | 795703<br>43 | 301 | MA5-20109 | Aluora<br>488 | 493/51<br>8 | GFP | 1:1000 | 5 |

### Cycle configuration for the automated immunofluorescence protocol

Cycle configuration was designed according to epitope stability, antigen abundance and spectral compatibility, summarized in **Table 1**. Highly expressed and stable epitopes were assigned lower intensity fluorophores and positioned in later cycles, while co-expressed markers were separated into non-consecutive cycles, with minimal spectral overlap.

### Implementation and integration of the in situ Proximity Ligation Assay on Spatial Station

The isPLA was integrated as the first cycle of the automated workflow. Following completion of the full isPLA cycle, the deck layout and reagents were reconfigured during approximately 10 min, followed by initiation of the mIF protocol for a combined workflow. For the isPLA protocol, reagents were kept in a Microtube Rack (Hamilton, Cat# 6600409-01), while wash buffers (TBS-1, TBS, PBS) were stored in a 12-trough Reservoir (Parhelia Biosciences, Cat#40205). The automated isPLA workflow followed the manufacturer’s instructions for the NaveniBright PD1/PD-L1 BOND RX HRP kit (Navinci Diagnostics, Cat# 60032) with minor modification up until addition of their chromogenic detection at the Substrate development step, at which point the detection system was replaced with the Aluora™ 647 dye to facilitate a combined mIF and isPLA workflow. The modifications are listed below and a more detailed description of protocol parameters and deck setup can be found in the attached PSS run report **(Figure S3)**.

### Modifications to the in situ Proximity Ligation Assay for automated implementation and integration

Endogenous peroxidase activity was quenched with 3% hydrogen peroxide (Thermo Fisher Scientific, Cat# AS100HRP) for 30 min, followed by three washes in 1X TBS-T with 3 min incubations. All temperature dependent steps were performed at 38°C, except Enzyme reaction 2 which was kept at 36°C. Primary antibodies were diluted to a specified dilution **(Table 2)**, followed by 3 × 2 min washes in 1X TBS-T, while probe-conjugated secondary antibodies (Navinci Diagnostics, Cat# 60032) were diluted (1:40) and applied in two consecutive 60 min incubations, followed by extensive washes in 1X TBS-T (two rinses, 2 × 2 min incubation, 20 min incubation, one rinse, 20 min incubation, three rinses). Enzyme reaction 1 (1:20) (Navinci Diagnostics, Cat# 60032) was applied for probe ligation for 30 min, followed by 2 × 2 min washes in TBS-T, while Enzyme reaction 2 (1:20) (Navinci Diagnostics, Cat# 60032) was applied for amplification in two consecutive 45 min incubations. HRP-labeled oligonucleotides (1:400) (Navinci Diagnostics, Cat# 60032) were incubated for 30 min followed by 2 × 2 min washes in 1X TBS and one wash in 1X PBS. At this point, instead of adding the standard chromogenic detection provided by Navinci Diagnostics, the Aluora™ dye 647 (Thermo Fisher Scientific, Cat# AS100HRP) was incubated for 10 min, followed by 2 × 2 min washes with 1X PBS. Nuclear staining was performed with DAPI (Thermo Fisher Scientific, Cat# H3570) and applied twice for 4 min each, followed by two washes in 1X PBS with 2 min incubation, two rinses with 1X TBS-T, one 0.1X TBS-T rinse and a final 1X PBS wash. Samples then underwent HIER with 1X Tris-EDTA buffer (pH 9) (Abcam, Cat# ab93684) for 45 min at 90°C before initiating the automated mIF workflow. The technical negative control was processed identically, except for the addition of 1X PBS instead of primary antibody.

**Table 2.** Cycle order, protein markers and antibodies with corresponding Aluora™ dyes used in the isPLA part of the automated workflow.

| Cycle order | Marker | Supplier | Clone | Lot no. | Catalog no. | Aluora<sup>T</sup><br><sup>M</sup> dye | Ex/Em | Emission filter | Dilution | Exposure time (ms) |
| --- | --- | --- | --- | --- | --- | --- | --- | --- | --- | --- |
| 1 | isPLA (PD-1/PD-L1) | Navinci Diagnostics | Naveni PD-1<sup>a)</sup>, Naveni PD-L1<sup>b)</sup> | 08042503 | 50043, 50044 | Aluora 647 | 652/670 | Cy5 | 1:40 | 40 ms |

<sup>a)</sup> Based on clone EH33, Cell Signaling Technology; <sup>b)</sup> Based on clone SP142, Abcam RabMAb®

### Image acquisition

All samples were imaged on the Invitrogen™ EVOS™ Spatial Imaging System (Thermo Fisher Scientific) with a 20x objective using the same unmixing protocol. Exposure times are summarized in **Table 1 and 2**.

### Unmixing protocol setup

The unmixing algorithm requires distinct reference spectra for each fluorophore, which were derived from Single Color Controls (SCC), tissue samples stained with a single fluorophore. SCCs were acquired from FPPE bladder cancer tumor tissue samples for all markers except the Aluora 647 channel, for which an artificial SCC (Thermo Fisher Scientific) based on a FFPE tonsil tissue was used. All FFPE bladder cancer SCC were processed manually according to the manufacturer’s instructions for the Aluora™ Spatial Amplification kit (Thermo Fisher Scientific, Cat# AS100HRP). The unmixing protocol was set up by scanning each SCC in all relevant channels, starting with an unstained sample to capture tissue background. The EVOS™ Imaging Software generated an unmixing report for quality control of the spectral separation **(Figure S4)**.

### Image analysis

Multiplex immunofluorescence whole-slide images acquired on the EVOS S1000 were analyzed using the Python-based image analysis workflow PIPEX [20]. Image channels were extracted from OME-TIFF files as individual TIFF images while preserving spatial information.

For the EV5 bladder cancer whole-slide samples, preprocessing was applied to the DAPI channel to improve nuclear segmentation. For the bladder cancer sample and the tonsil sample, preprocessing of the DAPI channel was not required prior to segmentation.

Cell segmentation was performed in PIPEX using the StarDist2D (2D_versatile_fluo) algorithm for nuclear detection on the DAPI channel, followed by single-cell quantification of fluorescence marker intensities from the corresponding image channels. The resulting single-cell data were collected and exported as .csv and anndata files for downstream analysis.

Cell marker positivity was assigned based on marker-positivity rules (otsu-based thresholding) derived from the raw mean intensities of each channel. The cell phenotypes were then established by a hierarchical decision tree and then used to quantify cellular composition at both whole-sample and region-of-interest levels. Summary tables were generated for downstream visualization and analysis.

For the isPLA-stained samples, dot detection was performed in PIPEX using the Big-FISH spot detection module [21]. Dot detection was applied to the bladder cancer samples and the tonsil samples. Detection settings were optimized separately for each tissue type based on tissue morphology and isPLA signal characteristics. These tissue-specific settings were used to optimize within-sample dot detection and were not intended for direct comparison of absolute dot counts between tissue types. Detected spots were assigned to segmented cells to classify cells as isPLA-positive (> 1 dot detection) or isPLA-negative (< 1 dot detection).

For visualization and quality control, phenotype annotations, and isPLA-related outputs were exported for overlay in TissUUmaps (3.1) [22,23]. This enabled visual inspection of segmentation quality, phenotype assignment, and spatial localization of isPLA-positive cells in relation to the original fluorescence images and annotated regions.

### Data and code

The relevant code from the analysis can be found and accessed via the Spatial Proteomics Github webpage: https://github.com/NBISweden/sbpda/tree/main/Units/Spatial_Proteomics/Codes/Scripts

