## Supplemental Figure 1 for "A workflow for combined detection of protein interactions and cell types for translational studies"

### Supplementary Figure S1A

Figure_S1A_tonsil_master_table_vertical

| field | Whole_sample | GC_1 | GC_2 | GC_3 |
|---|---|---|---|---|
| tissue_section | Tonsil | Tonsil | Tonsil | Tonsil |
| total_cells | 29589 | 9895 | 9541 | 10153 |
| ispla_positive_cells | 5242 | 1754 | 2246 | 1242 |
| ispla_positive_fraction | 0.17716 | 0.177261 | 0.235405 | 0.122328 |
| ispla_positive_percent | 17.716 | 17.7261 | 23.5405 | 12.2328 |
| ispla_negative_cells | 24347 | 8141 | 7295 | 8911 |
| ispla_negative_fraction | 0.82284 | 0.822739 | 0.764595 | 0.877672 |
| ispla_negative_percent | 82.284 | 82.2739 | 76.4595 | 87.7672 |
| total_ispla_positive_immune_cells | 5215 | 1745 | 2235 | 1235 |
| cytotoxic_t_cell_cells | 2006 | 1034 | 282 | 690 |
| cytotoxic_t_cell_fraction_of_all_cells | 0.067795 | 0.104497 | 0.029557 | 0.06796 |
| cytotoxic_t_cell_percent_of_all_cells | 6.7795 | 10.4497 | 2.9557 | 6.796 |
| t_cell_cells | 7686 | 3051 | 1995 | 2640 |
| t_cell_fraction_of_all_cells | 0.259759 | 0.308338 | 0.209098 | 0.260022 |
| t_cell_percent_of_all_cells | 25.9759 | 30.8338 | 20.9098 | 26.0022 |
| other_immune_cells | 18172 | 5147 | 6677 | 6348 |
| other_immune_fraction_of_all_cells | 0.614147 | 0.520162 | 0.699822 | 0.625234 |
| other_immune_percent_of_all_cells | 61.4147 | 52.0162 | 69.9822 | 62.5234 |
| other_cells | 999 | 431 | 334 | 234 |
| other_fraction_of_all_cells | 0.033763 | 0.043557 | 0.035007 | 0.023047 |
| other_percent_of_all_cells | 3.3763 | 4.3557 | 3.5007 | 2.3047 |
| cytotoxic_t_cell_ispla_positive_cells | 329 | 147 | 52 | 130 |
| cytotoxic_t_cell_ispla_positive_fraction_within_phenotype | 0.164008 | 0.142166 | 0.184397 | 0.188406 |
| cytotoxic_t_cell_ispla_positive_percent_within_phenotype | 16.4008 | 14.2166 | 18.4397 | 18.8406 |
| cytotoxic_t_cell_ispla_negative_cells | 1677 | 887 | 230 | 560 |
| cytotoxic_t_cell_ispla_negative_fraction_within_phenotype | 0.835992 | 0.857834 | 0.815603 | 0.811594 |
| cytotoxic_t_cell_ispla_negative_percent_within_phenotype | 83.5992 | 85.7834 | 81.5603 | 81.1594 |
| t_cell_ispla_positive_cells | 2530 | 994 | 745 | 791 |
| t_cell_ispla_positive_fraction_within_phenotype | 0.32917 | 0.325795 | 0.373434 | 0.299621 |
| t_cell_ispla_positive_percent_within_phenotype | 32.917 | 32.5795 | 37.3434 | 29.9621 |
| t_cell_ispla_negative_cells | 5156 | 2057 | 1250 | 1849 |
| t_cell_ispla_negative_fraction_within_phenotype | 0.67083 | 0.674205 | 0.626566 | 0.700379 |
| t_cell_ispla_negative_percent_within_phenotype | 67.083 | 67.4205 | 62.6566 | 70.0379 |
| other_immune_ispla_positive_cells | 2356 | 604 | 1438 | 314 |
| other_immune_ispla_positive_fraction_within_phenotype | 0.12965 | 0.11735 | 0.215366 | 0.049464 |
| other_immune_ispla_positive_percent_within_phenotype | 12.965 | 11.735 | 21.5366 | 4.9464 |
| other_immune_ispla_negative_cells | 15816 | 4543 | 5239 | 6034 |
| other_immune_ispla_negative_fraction_within_phenotype | 0.87035 | 0.88265 | 0.784634 | 0.950536 |
| other_immune_ispla_negative_percent_within_phenotype | 87.035 | 88.265 | 78.4634 | 95.0536 |
| cytotoxic_t_cell_fraction_within_ispla_positive_immune_cells | 0.063087 | 0.084241 | 0.023266 | 0.105263 |
| cytotoxic_t_cell_percent_within_ispla_positive_immune_cells | 6.3087 | 8.4241 | 2.3266 | 10.5263 |
| t_cell_fraction_within_ispla_positive_immune_cells | 0.485139 | 0.569628 | 0.333333 | 0.640486 |
| t_cell_percent_within_ispla_positive_immune_cells | 48.5139 | 56.9628 | 33.3333 | 64.0486 |
| other_immune_fraction_within_ispla_positive_immune_cells | 0.451774 | 0.346132 | 0.6434 | 0.254251 |
| other_immune_percent_within_ispla_positive_immune_cells | 45.1774 | 34.6132 | 64.34 | 25.4251 |

#### Supplementary Figure S1B

Figure_S1B_bladder_TLS_master_table_vertical

| field | Whole_sample | TLS_1 | TLS_2 | TLS_3 |
|---|---|---|---|---|
| tissue_section | Bladder cancer | Bladder cancer | Bladder cancer | Bladder cancer |
| total_cells | 10559 | 4576 | 2723 | 3260 |
| ispla_positive_cells | 300 | 124 | 91 | 85 |
| ispla_positive_fraction | 0.028412 | 0.027098 | 0.033419 | 0.026074 |
| ispla_positive_percent | 2.8412 | 2.7098 | 3.3419 | 2.6074 |
| ispla_negative_cells | 10259 | 4452 | 2632 | 3175 |
| ispla_negative_fraction | 0.971588 | 0.972902 | 0.966581 | 0.973926 |
| ispla_negative_percent | 97.1588 | 97.2902 | 96.6581 | 97.3926 |
| total_ispla_positive_immune_cells | 135 | 40 | 53 | 42 |
| cytotoxic_t_cell_cells | 363 | 132 | 169 | 62 |
| cytotoxic_t_cell_fraction_of_all_cells | 0.034378 | 0.028846 | 0.062064 | 0.019018 |
| cytotoxic_t_cell_percent_of_all_cells | 3.4378 | 2.8846 | 6.2064 | 1.9018 |
| t_cell_cells | 481 | 194 | 213 | 74 |
| t_cell_fraction_of_all_cells | 0.045554 | 0.042395 | 0.078223 | 0.022699 |
| t_cell_percent_of_all_cells | 4.5554 | 4.2395 | 7.8223 | 2.2699 |
| other_immune_cells | 4022 | 1612 | 1064 | 1346 |
| other_immune_fraction_of_all_cells | 0.380907 | 0.352273 | 0.390746 | 0.412883 |
| other_immune_percent_of_all_cells | 38.0907 | 35.2273 | 39.0746 | 41.2883 |
| other_cells | 5443 | 2482 | 1223 | 1738 |
| other_fraction_of_all_cells | 0.515484 | 0.542395 | 0.449137 | 0.533129 |
| other_percent_of_all_cells | 51.5484 | 54.2395 | 44.9137 | 53.3129 |
| cytotoxic_t_cell_ispla_positive_cells | 12 | 4 | 3 | 5 |
| cytotoxic_t_cell_ispla_positive_fraction_within_phenotype | 0.033058 | 0.030303 | 0.017751 | 0.080645 |
| cytotoxic_t_cell_ispla_positive_percent_within_phenotype | 3.3058 | 3.0303 | 1.7751 | 8.0645 |
| cytotoxic_t_cell_ispla_negative_cells | 351 | 128 | 166 | 57 |
| cytotoxic_t_cell_ispla_negative_fraction_within_phenotype | 0.966942 | 0.969697 | 0.982249 | 0.919355 |
| cytotoxic_t_cell_ispla_negative_percent_within_phenotype | 96.6942 | 96.9697 | 98.2249 | 91.9355 |
| t_cell_ispla_positive_cells | 23 | 7 | 13 | 3 |
| t_cell_ispla_positive_fraction_within_phenotype | 0.047817 | 0.036082 | 0.061033 | 0.040541 |
| t_cell_ispla_positive_percent_within_phenotype | 4.7817 | 3.6082 | 6.1033 | 4.0541 |
| t_cell_ispla_negative_cells | 458 | 187 | 200 | 71 |
| t_cell_ispla_negative_fraction_within_phenotype | 0.952183 | 0.963918 | 0.938967 | 0.959459 |
| t_cell_ispla_negative_percent_within_phenotype | 95.2183 | 96.3918 | 93.8967 | 95.9459 |
| other_immune_ispla_positive_cells | 100 | 29 | 37 | 34 |
| other_immune_ispla_positive_fraction_within_phenotype | 0.024863 | 0.01799 | 0.034774 | 0.02526 |
| other_immune_ispla_positive_percent_within_phenotype | 2.4863 | 1.799 | 3.4774 | 2.526 |
| other_immune_ispla_negative_cells | 3922 | 1583 | 1027 | 1312 |
| other_immune_ispla_negative_fraction_within_phenotype | 0.975137 | 0.98201 | 0.965226 | 0.97474 |
| other_immune_ispla_negative_percent_within_phenotype | 97.5137 | 98.201 | 96.5226 | 97.474 |
| cytotoxic_t_cell_fraction_within_ispla_positive_immune_cells | 0.088889 | 0.1 | 0.056604 | 0.119048 |
| cytotoxic_t_cell_percent_within_ispla_positive_immune_cells | 8.8889 | 10.0 | 5.6604 | 11.9048 |
| t_cell_fraction_within_ispla_positive_immune_cells | 0.17037 | 0.175 | 0.245283 | 0.071429 |
| t_cell_percent_within_ispla_positive_immune_cells | 17.037 | 17.5 | 24.5283 | 7.1429 |
| other_immune_fraction_within_ispla_positive_immune_cells | 0.740741 | 0.725 | 0.698113 | 0.809524 |
| other_immune_percent_within_ispla_positive_immune_cells | 74.0741 | 72.5 | 69.8113 | 80.9524 |
