## Supplemental Figure 2 for "A workflow for combined detection of protein interactions and cell types for translational studies"

### Run Report

| | | | |
|---|---|---|---|
| Protocol Name: | Aluora Dyes 7 Abs v1.0 | Last Modified: | Feb 03, 2026 01:39:17 PM |
| Operator: | spu | Started on: | Feb 23, 2026 10:09:36 AM |
| Enclosure Status: | Present | Ended on: | Feb 24, 2026 08:32:33 AM |
| Serial Number: | PRPEM2207 | Software Version | 3.3.2.00005 |

### Completed

#### Deck Layout

| | |
|---|---|
| 1<br>Parhelia ST12 w/Sheath on HHC<br>Parhelia ST12 with Thermal Sheath<br>Hamilton Heater Cooler<br>Scan status: Incorrect (User confirmed) | 5<br>[empty]<br><br>Scan status: Occupied (User confirmed) |
| 2<br>[empty]<br><br>Scan status: Correct | 6<br>Tip Rack<br>300 µL Framed Conductive No Filter<br><br>Scan status: ConfirmationRequired (User confirmed) |
| 3<br>Plate<br>Seahorse - Agilent, 24, Deep Well, U-Bottom, 10mL, Clear, Polypropylene<br><br>Scan status: Correct | 7<br>[empty]<br><br>Scan status: Correct |
| 4<br>PBS and AR cycles 1-4<br>Seahorse - Agilent, 24, Deep Well, U-Bottom, 10mL, Clear, Polypropylene<br><br>Scan status: Correct | 8<br>H2O2_Blocking_Abs_HRP_Dyes<br>Seahorse - Agilent, 96, Deep Well, V-Bottom, 2mL, Clear, Polypropylene<br><br>Scan status: Incorrect (User confirmed) |

### Steps

○ Lighting Step

○ How Many Samples Step

1. #1 – Add H2O2 (does not repeat in other cycles) (Disabled)

2. H2O2 incubation (30 min) (Disabled)

3. Transfer Samples Step - ar buffer

   - Step Type: Transfer samples
   - Pipetting Tool: TwoChannel
   - Liquid Type: Aqueous
   - Aspirate Volume: 150
   - Dispense Volume: 150
   - Tips Name: 300 µL Framed Conductive No Filter
   - New Tips Each Transfer: True

4. Heat to ER temp (90C)

   - Step Type: Heat cool
   - Temperature: 90 °C

5. #1 ER Topoff 1/2 (mixing delay 9 min) Copy

   - Step Type: Replicate samples
   - Pipetting Tool: TwoChannel
   - Liquid Type: $Aq30-BlMuCh-multi-dispense-buffer-chamfered-cover-pad-blowout
   - Aspirate Volume: 50
   - Dispense Volume: 50
   - Tips Name: 300 µL Framed Conductive No Filter
   - New Tips Each Transfer: False

6. HHC Off Copy

   - Step Type: Heat cool
   - Temperature: Off

7. #1 ER Topoff 2/2 (mixing delay 8 min) Copy

   - Step Type: Replicate samples
   - Pipetting Tool: TwoChannel
   - Liquid Type: $Aq30-BlMuCh-multi-dispense-buffer-chamfered-cover-pad-blowout
   - Aspirate Volume: 50
   - Dispense Volume: 50
   - Tips Name: 300 µL Framed Conductive No Filter
   - New Tips Each Transfer: False

8. Set temp to 20C Copy

   - Step Type: Heat cool
   - Temperature: 20 °C

9. Transfer Samples Step - pbs

   - Step Type: Transfer samples
   - Pipetting Tool: TwoChannel
   - Liquid Type: Aqueous
   - Aspirate Volume: 150
   - Dispense Volume: 150
   - Tips Name: 300 µL Framed Conductive No Filter
   - New Tips Each Transfer: True

10. #1 – 3xWash and Blocking Buffer

    - Step Type: Hit picking
    - Pipetting tool: OneChannel
    - Liquid type: $AqBlMuCh-multi-dispense-buffer-chamfered-cover-pad-blowout10
    - Worklist file: #1 3XWash - Blocking Buffer.csv
    - Volume: From worklist
    - Tips Name: 300 µL Framed Conductive No Filter
    - New tips for each transfer: False

11. Blocking Buffer incubation (45 min)

    - Step Type: Pause
    - Duration: 2700 seconds

12. #1 – Add Primary antibody

    - Step Type: Replicate samples
    - Pipetting Tool: OneChannel
    - Liquid Type: $Aqueous-buffer-chamfered-cover-pad-1
    - Aspirate Volume: 120
    - Dispense Volume: 120
    - Tips Name: 300 µL Framed Conductive No Filter
    - New Tips Each Transfer: False

13. Primary antibody incubation (60 min)

    - Step Type: Pause
    - Duration: 3600 seconds

14. #1 – 3xWash and Secondary-HRP add

    - Step Type: Hit picking
    - Pipetting tool: TwoChannel
    - Liquid type: $AqBlMuCh-multi-dispense-buffer-chamfered-cover-pad-blowout10
    - Worklist file: #1 3XWash - HRP.csv
    - Volume: From worklist
    - Tips Name: 300 µL Framed Conductive No Filter

- New tips for each transfer: False

15. Secondary-HRP incubation step (45 min)

    - Step Type: Pause
    - Duration: 2100 seconds

16. #1 – 3xWash and Aluora dye

    - Step Type: Hit picking
    - Pipetting tool: TwoChannel
    - Liquid type: $AqBlMuCh-multi-dispense-buffer-chamfered-cover-pad-blowout10
    - Worklist file: #1 3XWash - Aluora Dye.csv
    - Volume: From worklist
    - Tips Name: 300 µL Framed Conductive No Filter
    - New tips for each transfer: False

17. Aluora Dye incubation step (10 min)

    - Step Type: Pause
    - Duration: 600 seconds

18. #1 – 3xWash and AR Buffer add

    - Step Type: Hit picking
    - Pipetting tool: TwoChannel
    - Liquid type: $AqBlMuCh-multi-dispense-buffer-chamfered-cover-pad-blowout10
    - Worklist file: #1 3XWash - AR.csv
    - Volume: From worklist
    - Tips Name: 300 µL Framed Conductive No Filter
    - New tips for each transfer: False

19. Heat to ER temp (90C)

    - Step Type: Heat cool
    - Temperature: 90 °C

20. #1 ER Topoff 1/2 (mixing delay 9 min)

    - Step Type: Replicate samples
    - Pipetting Tool: TwoChannel
    - Liquid Type: $Aq30-BlMuCh-multi-dispense-buffer-chamfered-cover-pad-blowout
    - Aspirate Volume: 50
    - Dispense Volume: 50
    - Tips Name: 300 µL Framed Conductive No Filter
    - New Tips Each Transfer: False

21. HHC Off

    - Step Type: Heat cool
    - Temperature: Off

22. #1 ER Topoff 2/2 (mixing delay 8 min)

    - Step Type: Replicate samples
    - Pipetting Tool: TwoChannel
    - Liquid Type: $Aq30-BlMuCh-multi-dispense-buffer-chamfered-cover-pad-blowout
    - Aspirate Volume: 50
    - Dispense Volume: 50
    - Tips Name: 300 µL Framed Conductive No Filter
    - New Tips Each Transfer: False

23. Set temp to 20C

    - Step Type: Heat cool
    - Temperature: 20 °C

24. ———#2 3X PBS Wash + Blocking Buffer———

    - Step Type: Hit picking
    - Pipetting tool: TwoChannel
    - Liquid type: $AqBlMuCh-multi-dispense-buffer-chamfered-cover-pad-blowout10
    - Worklist file: #2 3XWash - Blocking Buffer.csv
    - Volume: From worklist
    - Tips Name: 300 µL Framed Conductive No Filter
    - New tips for each transfer: False

25. Blocking Buffer incubation (45 min)

    - Step Type: Pause
    - Duration: 2700 seconds

26. #2 Add Primary antibody

    - Step Type: Replicate samples
    - Pipetting Tool: TwoChannel
    - Liquid Type: $Aq30-BlMuCh-multi-dispense-buffer-chamfered-cover-pad-blowout
    - Aspirate Volume: 120
    - Dispense Volume: 120
    - Tips Name: 300 µL Framed Conductive No Filter
    - New Tips Each Transfer: False

27. Primary antibody incubation (60 min)

    - Step Type: Pause
    - Duration: 3600 seconds

28. #2 3xWash and Secondary-HRP add

    - Step Type: Hit picking
    - Pipetting tool: TwoChannel
    - Liquid type: $AqBlMuCh-multi-dispense-buffer-chamfered-cover-pad-blowout10
    - Worklist file: #2 3XWash - HRP.csv
    - Volume: From worklist

- Tips Name: 300 µL Framed Conductive No Filter
- New tips for each transfer: False

29. Secondary-HRP incubation step (45 min)
    - Step Type: Pause
    - Duration: 2100 seconds
30. #2 3xWash and Aluora dye
    - Step Type: Hit picking
    - Pipetting tool: TwoChannel
    - Liquid type: $AqBlMuCh-multi-dispense-buffer-chamfered-cover-pad-blowout10
    - Worklist file: #2 3XWash - Aluora Dye.csv
    - Volume: From worklist
    - Tips Name: 300 µL Framed Conductive No Filter
    - New tips for each transfer: False
31. Aluora Dye incubation step (10 min)
    - Step Type: Pause
    - Duration: 600 seconds
32. #2 3xWash and AR Buffer add
    - Step Type: Hit picking
    - Pipetting tool: TwoChannel
    - Liquid type: $Aqueous-buffer-chamfered-cover-pad-1
    - Worklist file: #2 3XWash - AR.csv
    - Volume: From worklist
    - Tips Name: 300 µL Framed Conductive No Filter
    - New tips for each transfer: False
33. Heat to ER temp (90C)
    - Step Type: Heat cool
    - Temperature: 90 °C
34. #2 ER Topoff 1/2 (mixing delay 9 min)
    - Step Type: Replicate samples
    - Pipetting Tool: TwoChannel
    - Liquid Type: $Aq30-BlMuCh-multi-dispense-buffer-chamfered-cover-pad-blowout
    - Aspirate Volume: 50
    - Dispense Volume: 50
    - Tips Name: 300 µL Framed Conductive No Filter
    - New Tips Each Transfer: False
35. HHC Off
    - Step Type: Heat cool
    - Temperature: Off
36. #2 ER Topoff 2/2 (mixing delay 8 min)
    - Step Type: Replicate samples
    - Pipetting Tool: TwoChannel
    - Liquid Type: $Aq30-BlMuCh-multi-dispense-buffer-chamfered-cover-pad-blowout
    - Aspirate Volume: 50
    - Dispense Volume: 50
    - Tips Name: 300 µL Framed Conductive No Filter
    - New Tips Each Transfer: False
37. Set temp to 20C
    - Step Type: Heat cool
    - Temperature: 20 °C
38. ———#3 3X PBS Wash + Blocking Buffer———
    - Step Type: Hit picking
    - Pipetting tool: TwoChannel
    - Liquid type: $AqBlMuCh-multi-dispense-buffer-chamfered-cover-pad-blowout10
    - Worklist file: #3 3XWash - Blocking Buffer.csv
    - Volume: From worklist
    - Tips Name: 300 µL Framed Conductive No Filter
    - New tips for each transfer: False
39. Blocking Buffer incubation (45 min)
    - Step Type: Pause
    - Duration: 2700 seconds
40. #3 Add Primary antibody
    - Step Type: Replicate samples
    - Pipetting Tool: TwoChannel
    - Liquid Type: $Aqueous-buffer-chamfered-cover-pad-1
    - Aspirate Volume: 120
    - Dispense Volume: 120
    - Tips Name: 300 µL Framed Conductive No Filter
    - New Tips Each Transfer: False
41. Primary antibody incubation (60 min)
    - Step Type: Pause
    - Duration: 3600 seconds
42. #3 3xWash and Secondary-HRP add
    - Step Type: Hit picking
    - Pipetting tool: TwoChannel
    - Liquid type: $AqBlMuCh-multi-dispense-buffer-chamfered-cover-pad-blowout10
    - Worklist file: #3 3XWash - HRP.csv

- Volume: From worklist
- Tips Name: 300 µL Framed Conductive No Filter
- New tips for each transfer: False

43. Secondary-HRP incubation step (45 min)

- Step Type: Pause
- Duration: 2100 seconds

44. #3 3xWash and Aluora dye

- Step Type: Hit picking
- Pipetting tool: TwoChannel
- Liquid type: $AqBIMuCh-multi-dispense-buffer-chamfered-cover-pad-blowout10
- Worklist file: #3 3XWash - Aluora Dye.csv
- Volume: From worklist
- Tips Name: 300 µL Framed Conductive No Filter
- New tips for each transfer: False

45. Aluora Dye incubation step (10 min)

- Step Type: Pause
- Duration: 600 seconds

46. #3 3xWash and AR Buffer add

- Step Type: Hit picking
- Pipetting tool: TwoChannel
- Liquid type: $AqBIMuCh-multi-dispense-buffer-chamfered-cover-pad-blowout10
- Worklist file: #3 3XWash - AR.csv
- Volume: From worklist
- Tips Name: 300 µL Framed Conductive No Filter
- New tips for each transfer: False

47. Heat to ER temp (90C)

- Step Type: Heat cool
- Temperature: 90 °C

48. #3 ER Topoff 1/2 (mixing delay 9 min)

- Step Type: Replicate samples
- Pipetting Tool: TwoChannel
- Liquid Type: $Aq30-BIMuCh-multi-dispense-buffer-chamfered-cover-pad-blowout
- Aspirate Volume: 50
- Dispense Volume: 50
- Tips Name: 300 µL Framed Conductive No Filter
- New Tips Each Transfer: False

49. HHC Off

- Step Type: Heat cool
- Temperature: Off

50. #3 ER Topoff 2/2 (mixing delay 8 min)

- Step Type: Replicate samples
- Pipetting Tool: TwoChannel
- Liquid Type: $Aq30-BIMuCh-multi-dispense-buffer-chamfered-cover-pad-blowout
- Aspirate Volume: 50
- Dispense Volume: 50
- Tips Name: 300 µL Framed Conductive No Filter
- New Tips Each Transfer: False

51. Set temp to 20C

- Step Type: Heat cool
- Temperature: 20 °C

52. ———#4 3X PBS Wash + Blocking Buffer———

- Step Type: Hit picking
- Pipetting tool: TwoChannel
- Liquid type: $AqBIMuCh-multi-dispense-buffer-chamfered-cover-pad-blowout10
- Worklist file: #4 3XWash - Blocking Buffer.csv
- Volume: From worklist
- Tips Name: 300 µL Framed Conductive No Filter
- New tips for each transfer: False

53. Blocking Buffer incubation (45 min)

- Step Type: Pause
- Duration: 2700 seconds

54. #4 Add Primary antibody

- Step Type: Replicate samples
- Pipetting Tool: TwoChannel
- Liquid Type: $Aq30-BIMuCh-multi-dispense-buffer-chamfered-cover-pad-blowout
- Aspirate Volume: 120
- Dispense Volume: 120
- Tips Name: 300 µL Framed Conductive No Filter
- New Tips Each Transfer: False

55. Primary antibody incubation (60 min)

- Step Type: Pause
- Duration: 3600 seconds

56. #4 3xWash and Secondary-HRP add

- Step Type: Hit picking
- Pipetting tool: TwoChannel
- Liquid type: $AqBIMuCh-multi-dispense-buffer-chamfered-cover-pad-blowout10

- Worklist file: #4 3XWash - HRP.csv
- Volume: From worklist
- Tips Name: 300 µL Framed Conductive No Filter
- New tips for each transfer: False

57. Secondary-HRP incubation step (45 min)

- Step Type: Pause
- Duration: 2100 seconds

58. #4 3xWash and Aluora dye

- Step Type: Hit picking
- Pipetting tool: TwoChannel
- Liquid type: $AqBlMuCh-multi-dispense-buffer-chamfered-cover-pad-blowout10
- Worklist file: #4 3XWash - Aluora Dye.csv
- Volume: From worklist
- Tips Name: 300 µL Framed Conductive No Filter
- New tips for each transfer: False

59. Aluora Dye incubation step (10 min)

- Step Type: Pause
- Duration: 600 seconds

60. #4 3xWash and AR Buffer add

- Step Type: Hit picking
- Pipetting tool: TwoChannel
- Liquid type: $AqBlMuCh-multi-dispense-buffer-chamfered-cover-pad-blowout10
- Worklist file: #4 3XWash - AR.csv
- Volume: From worklist
- Tips Name: 300 µL Framed Conductive No Filter
- New tips for each transfer: False

61. Heat to ER temp (90C)

- Step Type: Heat cool
- Temperature: 90 °C

62. #4 ER Topoff 1/2 (mixing delay 9 min)

- Step Type: Replicate samples
- Pipetting Tool: TwoChannel
- Liquid Type: $Aq30-BlMuCh-multi-dispense-buffer-chamfered-cover-pad-blowout
- Aspirate Volume: 50
- Dispense Volume: 50
- Tips Name: 300 µL Framed Conductive No Filter
- New Tips Each Transfer: False

63. HHC Off

- Step Type: Heat cool
- Temperature: Off

64. #4 ER Topoff 2/2 (mixing delay 8 min)

- Step Type: Replicate samples
- Pipetting Tool: TwoChannel
- Liquid Type: $Aq30-BlMuCh-multi-dispense-buffer-chamfered-cover-pad-blowout
- Aspirate Volume: 50
- Dispense Volume: 50
- Tips Name: 300 µL Framed Conductive No Filter
- New Tips Each Transfer: False

65. Set temp to 20C

- Step Type: Heat cool
- Temperature: 20 °C

66. ———#5 3X PBS Wash + Blocking Buffer———

- Step Type: Hit picking
- Pipetting tool: TwoChannel
- Liquid type: $AqBlMuCh-multi-dispense-buffer-chamfered-cover-pad-blowout10
- Worklist file: #5 3XWash - Blocking Buffer.csv
- Volume: From worklist
- Tips Name: 300 µL Framed Conductive No Filter
- New tips for each transfer: False

67. Blocking Buffer incubation (45 min)

- Step Type: Pause
- Duration: 2700 seconds

68. #5 Add Primary antibody

- Step Type: Replicate samples
- Pipetting Tool: TwoChannel
- Liquid Type: $Aq30-BlMuCh-multi-dispense-buffer-chamfered-cover-pad-blowout
- Aspirate Volume: 120
- Dispense Volume: 120
- Tips Name: 300 µL Framed Conductive No Filter
- New Tips Each Transfer: False

69. Primary antibody incubation (60 min)

- Step Type: Pause
- Duration: 3600 seconds

70. #5 3xWash and Secondary-HRP add

- Step Type: Hit picking
- Pipetting tool: TwoChannel

- Liquid type: $AqBIMuCh-multi-dispense-buffer-chamfered-cover-pad-blowout10
- Worklist file: #5 3XWash - HRP.csv
- Volume: From worklist
- Tips Name: 300 µL Framed Conductive No Filter
- New tips for each transfer: False

71. Secondary-HRP incubation step (45 min)

    - Step Type: Pause
    - Duration: 2100 seconds

72. #5 3xWash and Aluora dye

    - Step Type: Hit picking
    - Pipetting tool: TwoChannel
    - Liquid type: $AqBIMuCh-multi-dispense-buffer-chamfered-cover-pad-blowout10
    - Worklist file: #5 3XWash - Aluora Dye.csv
    - Volume: From worklist
    - Tips Name: 300 µL Framed Conductive No Filter
    - New tips for each transfer: False

73. Aluora Dye incubation step (10 min)

    - Step Type: Pause
    - Duration: 600 seconds

74. #5 3xWash and AR Buffer add (Disabled)

75. Heat to ER temp (90C) (Disabled)

76. #5 ER Topoff 1/2 (mixing delay 9 min) (Disabled)

77. HHC Off (Disabled)

78. #5 ER Topoff 2/2 (mixing delay 8 min) (Disabled)

79. Set temp to 20C (Disabled)

80. ———#6 3X PBS Wash + Blocking Buffer——— (Disabled)

81. Blocking Buffer incubation (45 min) (Disabled)

82. #6 Add Primary antibody (Disabled)

83. Primary antibody incubation (60 min) (Disabled)

84. #6 3xWash and Secondary-HRP add (Disabled)

85. Secondary-HRP incubation step (45 min) (Disabled)

86. #6 3xWash and Aluora dye (Disabled)

87. Aluora Dye incubation step (10 min) (Disabled)

88. #6 3xWash and AR Buffer add (Disabled)

89. Heat to ER temp (90C) (Disabled)

90. #6 ER Topoff 1/2 (mixing delay 9 min) (Disabled)

91. HHC Off (Disabled)

92. #6 ER Topoff 2/2 (mixing delay 8 min) (Disabled)

93. Set temp to 20C (Disabled)

94. ———#7 3X PBS Wash + Blocking Buffer——— (Disabled)

95. Blocking Buffer incubation (45 min) (Disabled)

96. #7 Add Primary antibody (Disabled)

97. Primary antibody incubation (60 min) (Disabled)

98. #7 3xWash and Secondary-HRP add (Disabled)

99. Secondary-HRP incubation step (45 min) (Disabled)

100. #7 3xWash and Aluora dye (Disabled)

101. #7 3xWash

     - Step Type: Hit picking
     - Pipetting tool: TwoChannel
     - Liquid type: $AqBIMuCh-multi-dispense-buffer-chamfered-cover-pad-blowout10
     - Worklist file: #7 3XWash.csv
     - Volume: From worklist
     - Tips Name: 300 µL Framed Conductive No Filter
     - New tips for each transfer: False

102. Aluora Dye incubation step (10 min) (Disabled)

103. Cool to 4C

     - Step Type: Heat cool
     - Temperature: 4 °C

104. Hold at 6C until user ends

     - Step Type: Pause
     - Duration: Pause until user resumes run.

### Liquids

[none]

### Tubes

[none]

### Protocol Setup

○ Lighting

- Deck Lights: Disable After Deck Verification
- Error Lighting Disabled: No

○ How Many Samples Step

- Step Type: How many samples
- Message: Select Tissue Slides locations
- Deck Positions: 1:Parhelia ST12 w/Sheath on HHC
- Selected Targets
    - Plate 1: A1

### Activity

**3. Transfer Samples Step _ ar buffer [10:09:33 AM - 10:09:54 AM]**

| | | Aspirate | | | | | | Dispense | | | | | |
|---|---|---|---|---|---|---|---|---|---|---|---|---|---|
| Channel | Liquid Type | Labware | Volume | Well | Start (CET) | End (CET) | Status | Labware | Volume | Well | Start (CET) | End (CET) | Status |
| 1 | Aqueous | 4 : PBS and AR cyc… | 150 | A6 | 10:09:35 AM | 10:09:43 AM | Ok | 1 : Parhelia ST12 … | 150 | A1 | 10:09:43 AM | 10:09:48 AM | Ok |

**4. Heat to ER temp (90C) [10:09:54 AM - 10:20:23 AM]**

**5. #1 ER Topoff 1/2 (mixing delay 9 min) Copy [10:20:23 AM - 10:29:59 AM]**

| | | Aspirate | | | | | | Dispense | | | | | |
|---|---|---|---|---|---|---|---|---|---|---|---|---|---|
| Channel | Liquid Type | Labware | Volume | Well | Start (CET) | End (CET) | Status | Labware | Volume | Well | Start (CET) | End (CET) | Status |
| 1 | $Aq30-BlMuCh-multi-dispense-buffer-chamfered-cover-pad-blowout | 4 : PBS and AR cyc… | 50 | A6 | 10:20:28 AM | 10:29:43 AM | Ok | 1 : Parhelia ST12 … | 50 | A1 | 10:29:43 AM | 10:29:54 AM | Ok |

**6. HHC Off Copy [10:29:59 AM - 10:30:02 AM]**

**7. #1 ER Topoff 2/2 (mixing delay 8 min) Copy [10:30:02 AM - 10:38:37 AM]**

| | | Aspirate | | | | | | Dispense | | | | | |
|---|---|---|---|---|---|---|---|---|---|---|---|---|---|
| Channel | Liquid Type | Labware | Volume | Well | Start (CET) | End (CET) | Status | Labware | Volume | Well | Start (CET) | End (CET) | Status |
| 1 | $Aq30-BlMuCh-multi-dispense-buffer-chamfered-cover-pad-blowout | 4 : PBS and AR cyc… | 50 | A6 | 10:30:07 AM | 10:38:21 AM | Ok | 1 : Parhelia ST12 … | 50 | A1 | 10:38:21 AM | 10:38:32 AM | Ok |

**8. Set temp to 20C Copy [10:38:37 AM - 10:48:25 AM]**

**9. Transfer Samples Step _ pbs [10:48:25 AM - 10:48:48 AM]**

| | | Aspirate | | | | | | Dispense | | | | | |
|---|---|---|---|---|---|---|---|---|---|---|---|---|---|
| Channel | Liquid Type | Labware | Volume | Well | Start (CET) | End (CET) | Status | Labware | Volume | Well | Start (CET) | End (CET) | Status |
| 1 | Aqueous | 3 : Plate | 150 | B4 | 10:48:30 AM | 10:48:38 AM | Ok | 1 : Parhelia ST12 … | 150 | A1 | 10:48:38 AM | 10:48:43 AM | Ok |

**10. #1 -- 3xWash and Blocking Buffer [10:48:48 AM - 10:50:09 AM]**

| | | Aspirate | | | | | | Dispense | | | | | |
|---|---|---|---|---|---|---|---|---|---|---|---|---|---|
| Channel | Liquid Type | Labware | Volume | Well | Start (CET) | End (CET) | Status | Labware | Volume | Well | Start (CET) | End (CET) | Status |
| 2 | $AqBlMuCh-multi-dispense-buffer-chamfered-cover-pad-blowout10 | 4 : PBS and AR cyc… | 150 | A1 | 10:48:53 AM | 10:49:01 AM | Ok | 1 : Parhelia ST12 … | 150 | A1 | 10:49:01 AM | 10:49:11 AM | Ok |
| 2 | $AqBlMuCh-multi-dispense-buffer-chamfered-cover-pad-blowout10 | 4 : PBS and AR cyc… | 150 | A1 | 10:49:11 AM | 10:49:19 AM | Ok | 1 : Parhelia ST12 … | 150 | A1 | 10:49:19 AM | 10:49:29 AM | Ok |
| 2 | $AqBlMuCh-multi-dispense-buffer-chamfered-cover-pad-blowout10 | 4 : PBS and AR cyc… | 150 | A1 | 10:49:29 AM | 10:49:37 AM | Ok | 1 : Parhelia ST12 … | 150 | A1 | 10:49:37 AM | 10:49:47 AM | Ok |
| 2 | $AqBlMuCh-multi-dispense-buffer-chamfered-cover-pad-blowout10 | 8 : H2O2_Blocking_… | 120 | B1 | 10:49:47 AM | 10:49:55 AM | Ok | 1 : Parhelia ST12 … | 120 | A1 | 10:49:55 AM | 10:50:04 AM | Ok |

**11. Blocking Buffer incubation (45 min) [10:50:09 AM - 11:35:09 AM]**

**12. #1 – Add Primary antibody [11:35:09 AM - 11:35:33 AM]**

| | | Aspirate | | | | | | Dispense | | | | | |
|---|---|---|---|---|---|---|---|---|---|---|---|---|---|
| Channel | Liquid Type | Labware | Volume | Well | Start (CET) | End (CET) | Status | Labware | Volume | Well | Start (CET) | End (CET) | Status |
| 2 | $Aqueous-buffer-chamfered-cover-pad-1 | 8 : H2O2_Blocking_… | 120 | C1 | 11:35:14 AM | 11:35:20 AM | Ok | 1 : Parhelia ST12 … | 120 | A1 | 11:35:20 AM | 11:35:28 AM | Ok |

**13. Primary antibody incubation (60 min) [11:35:33 AM - 12:35:33 PM]**

**14. #1 – 3xWash and Secondary-HRP add [12:35:33 PM - 12:36:53 PM]**

| | | Aspirate | | | | | | Dispense | | | | | |
|---|---|---|---|---|---|---|---|---|---|---|---|---|---|
| Channel | Liquid Type | Labware | Volume | Well | Start (CET) | End (CET) | Status | Labware | Volume | Well | Start (CET) | End (CET) | Status |
| 1 | $AqBlMuCh-multi-dispense-buffer-chamfered-cover-pad-blowout10 | 4 : PBS and AR cyc… | 150 | A1 | 12:35:38 PM | 12:35:52 PM | Ok | 1 : Parhelia ST12 … | 150 | A1 | 12:35:52 PM | 12:36:13 PM | Ok |
| 2 | $AqBlMuCh-multi-dispense-buffer-chamfered-cover-pad-blowout10 | 4 : PBS and AR cyc… | 150 | A1 | 12:35:38 PM | 12:35:52 PM | Ok | 1 : Parhelia ST12 … | 150 | A1 | 12:35:52 PM | 12:36:13 PM | Ok |
| 1 | $AqBlMuCh-multi-dispense-buffer-chamfered-cover-pad-blowout10 | 4 : PBS and AR cyc… | 150 | A1 | 12:36:13 PM | 12:36:28 PM | Ok | 1 : Parhelia ST12 … | 150 | A1 | 12:36:28 PM | 12:36:48 PM | Ok |
| 2 | $AqBlMuCh-multi-dispense-buffer-chamfered-cover-pad-blowout10 | 8 : H2O2_Blocking_… | 120 | D1 | 12:36:13 PM | 12:36:28 PM | Ok | 1 : Parhelia ST12 … | 120 | A1 | 12:36:28 PM | 12:36:48 PM | Ok |

**15. Secondary-HRP incubation step (45 min) [12:36:53 PM - 1:11:53 PM]**

**16. #1 – 3xWash and Aluora dye [1:11:53 PM - 1:13:12 PM]**

| | | Aspirate | | | | | | Dispense | | | | | |
|---|---|---|---|---|---|---|---|---|---|---|---|---|---|
| Channel | Liquid Type | Labware | Volume | Well | Start (CET) | End (CET) | Status | Labware | Volume | Well | Start (CET) | End (CET) | Status |
| 1 | $AqBlMuCh-multi-dispense-buffer-chamfered-cover-pad-blowout10 | 4 : PBS and AR cyc… | 150 | A1 | 1:11:58 PM | 1:12:12 PM | Ok | 1 : Parhelia ST12 … | 150 | A1 | 1:12:12 PM | 1:12:33 PM | Ok |
| 2 | $AqBlMuCh-multi-dispense-buffer-chamfered-cover-pad-blowout10 | 4 : PBS and AR cyc… | 150 | A1 | 1:11:58 PM | 1:12:12 PM | Ok | 1 : Parhelia ST12 … | 150 | A1 | 1:12:12 PM | 1:12:33 PM | Ok |
| 1 | $AqBlMuCh-multi-dispense-buffer-chamfered-cover-pad-blowout10 | 4 : PBS and AR cyc… | 150 | A1 | 1:12:33 PM | 1:12:47 PM | Ok | 1 : Parhelia ST12 … | 150 | A1 | 1:12:47 PM | 1:13:07 PM | Ok |
| 2 | $AqBlMuCh-multi-dispense-buffer-chamfered-cover-pad-blowout10 | 8 : H2O2_Blocking_… | 120 | E1 | 1:12:33 PM | 1:12:47 PM | Ok | 1 : Parhelia ST12 … | 120 | A1 | 1:12:47 PM | 1:13:07 PM | Ok |

**17. Aluora Dye incubation step (10 min) [1:13:12 PM - 1:23:12 PM]**

**18. #1 – 3xWash and AR Buffer add [1:23:12 PM - 1:24:33 PM]**

| | | Aspirate | | | | | | Dispense | | | | | |
|---|---|---|---|---|---|---|---|---|---|---|---|---|---|
| Channel | Liquid Type | Labware | Volume | Well | Start (CET) | End (CET) | Status | Labware | Volume | Well | Start (CET) | End (CET) | Status |
| 1 | $AqBlMuCh-multi-dispense-buffer-chamfered-cover-pad-blowout10 | 4 : PBS and AR cyc… | 150 | A1 | 1:23:17 PM | 1:23:31 PM | Ok | 1 : Parhelia ST12 … | 150 | A1 | 1:23:31 PM | 1:23:52 PM | Ok |
| 2 | $AqBlMuCh-multi-dispense-buffer-chamfered-cover-pad-blowout10 | 4 : PBS and AR cyc… | 150 | A1 | 1:23:17 PM | 1:23:31 PM | Ok | 1 : Parhelia ST12 … | 150 | A1 | 1:23:31 PM | 1:23:52 PM | Ok |
| 1 | $AqBlMuCh-multi-dispense-buffer-chamfered-cover-pad-blowout10 | 4 : PBS and AR cyc… | 150 | A1 | 1:23:52 PM | 1:24:07 PM | Ok | 1 : Parhelia ST12 … | 150 | A1 | 1:24:07 PM | 1:24:28 PM | Ok |
| 2 | $AqBlMuCh-multi-dispense-buffer-chamfered-cover-pad-blowout10 | 4 : PBS and AR cyc… | 150 | A6 | 1:23:52 PM | 1:24:07 PM | Ok | 1 : Parhelia ST12 … | 150 | A1 | 1:24:07 PM | 1:24:28 PM | Ok |

**19. Heat to ER temp (90C) [1:24:33 PM - 1:35:42 PM]**

**20. #1 ER Topoff 1/2 (mixing delay 9 min) [1:35:42 PM - 1:45:19 PM]**

| Channel | Liquid Type | Aspirate Labware | Volume | Well | Start (CET) | End (CET) | Status | Dispense Labware | Volume | Well | Start (CET) | End (CET) | Status |
|---|---|---|---|---|---|---|---|---|---|---|---|---|---|
| 1 | $Aq30-BlMuCh-multi-dispense-buffer-chamfered-cover-pad-blowout | 4 : PBS and AR cyc… | 50 | A6 | 1:35:47 PM | 1:45:02 PM | Ok | 1 : Parhelia ST12 … | 50 | A1 | 1:45:02 PM | 1:45:14 PM | Ok |

**21. HHC Off [1:45:19 PM - 1:45:22 PM]**

**22. #1 ER Topoff 2/2 (mixing delay 8 min) [1:45:22 PM - 1:53:57 PM]**

| Channel | Liquid Type | Aspirate Labware | Volume | Well | Start (CET) | End (CET) | Status | Dispense Labware | Volume | Well | Start (CET) | End (CET) | Status |
|---|---|---|---|---|---|---|---|---|---|---|---|---|---|
| 1 | $Aq30-BlMuCh-multi-dispense-buffer-chamfered-cover-pad-blowout | 4 : PBS and AR cyc… | 50 | A6 | 1:45:27 PM | 1:53:41 PM | Ok | 1 : Parhelia ST12 … | 50 | A1 | 1:53:41 PM | 1:53:53 PM | Ok |

**23. Set temp to 20C [1:53:57 PM - 2:04:29 PM]**

**24. --------#2 3X PBS Wash + Blocking Buffer-------- [2:04:29 PM - 2:05:49 PM]**

| Channel | Liquid Type | Aspirate Labware | Volume | Well | Start (CET) | End (CET) | Status | Dispense Labware | Volume | Well | Start (CET) | End (CET) | Status |
|---|---|---|---|---|---|---|---|---|---|---|---|---|---|
| 1 | $AqBlMuCh-multi-dispense-buffer-chamfered-cover-pad-blowout10 | 4 : PBS and AR cyc… | 150 | A2 | 2:04:34 PM | 2:04:48 PM | Ok | 1 : Parhelia ST12 … | 150 | A1 | 2:04:48 PM | 2:05:09 PM | Ok |
| 2 | $AqBlMuCh-multi-dispense-buffer-chamfered-cover-pad-blowout10 | 4 : PBS and AR cyc… | 150 | A2 | 2:04:34 PM | 2:04:48 PM | Ok | 1 : Parhelia ST12 … | 150 | A1 | 2:04:48 PM | 2:05:09 PM | Ok |
| 1 | $AqBlMuCh-multi-dispense-buffer-chamfered-cover-pad-blowout10 | 4 : PBS and AR cyc… | 150 | A2 | 2:05:10 PM | 2:05:24 PM | Ok | 1 : Parhelia ST12 … | 150 | A1 | 2:05:24 PM | 2:05:44 PM | Ok |
| 2 | $AqBlMuCh-multi-dispense-buffer-chamfered-cover-pad-blowout10 | 8 : H2O2_Blocking_… | 120 | B2 | 2:05:10 PM | 2:05:24 PM | Ok | 1 : Parhelia ST12 … | 120 | A1 | 2:05:24 PM | 2:05:44 PM | Ok |

**25. Blocking Buffer incubation (45 min) [2:05:49 PM - 2:50:49 PM]**

**26. #2 Add Primary antibody [2:50:49 PM - 2:51:17 PM]**

| Channel | Liquid Type | Aspirate Labware | Volume | Well | Start (CET) | End (CET) | Status | Dispense Labware | Volume | Well | Start (CET) | End (CET) | Status |
|---|---|---|---|---|---|---|---|---|---|---|---|---|---|
| 1 | $Aq30-BlMuCh-multi-dispense-buffer-chamfered-cover-pad-blowout | 8 : H2O2_Blocking_… | 120 | C2 | 2:50:55 PM | 2:51:02 PM | Ok | 1 : Parhelia ST12 … | 120 | A1 | 2:51:02 PM | 2:51:11 PM | Ok |

**27. Primary antibody incubation (60 min) [2:51:17 PM - 3:51:17 PM]**

**28. #2 3xWash and Secondary-HRP add [3:51:17 PM - 3:52:27 PM]**

| Channel | Liquid Type | Aspirate Labware | Volume | Well | Start (CET) | End (CET) | Status | Dispense Labware | Volume | Well | Start (CET) | End (CET) | Status |
|---|---|---|---|---|---|---|---|---|---|---|---|---|---|
| 1 | $AqBlMuCh-multi-dispense-buffer-chamfered-cover-pad-blowout10 | 4 : PBS and AR cyc… | 150 | A2 | 3:51:22 PM | 3:51:35 PM | Ok | 1 : Parhelia ST12 … | 150 | A1 | 3:51:35 PM | 3:51:53 PM | Ok |
| 2 | $AqBlMuCh-multi-dispense-buffer-chamfered-cover-pad-blowout10 | 4 : PBS and AR cyc… | 150 | A2 | 3:51:22 PM | 3:51:35 PM | Ok | 1 : Parhelia ST12 … | 150 | A1 | 3:51:35 PM | 3:51:53 PM | Ok |
| 1 | $AqBlMuCh-multi-dispense-buffer-chamfered-cover-pad-blowout10 | 4 : PBS and AR cyc… | 150 | A2 | 3:51:53 PM | 3:52:06 PM | Ok | 1 : Parhelia ST12 … | 150 | A1 | 3:52:06 PM | 3:52:22 PM | Ok |
| 2 | $AqBlMuCh-multi-dispense-buffer-chamfered-cover-pad-blowout10 | 8 : H2O2_Blocking_… | 120 | D2 | 3:51:53 PM | 3:52:06 PM | Ok | 1 : Parhelia ST12 … | 120 | A1 | 3:52:06 PM | 3:52:22 PM | Ok |

**29. Secondary-HRP incubation step (45 min) [3:52:27 PM - 4:27:27 PM]**

**30. #2 3xWash and Aluora dye [4:27:27 PM - 4:28:47 PM]**

| Channel | Liquid Type | Aspirate Labware | Volume | Well | Start (CET) | End (CET) | Status | Dispense Labware | Volume | Well | Start (CET) | End (CET) | Status |
|---|---|---|---|---|---|---|---|---|---|---|---|---|---|

| Channel | Liquid Type | Aspirate Labware | Volume | Well | Start (CET) | End (CET) | Status | Dispense Labware | Volume | Well | Start (CET) | End (CET) | Status |
|---|---|---|---|---|---|---|---|---|---|---|---|---|---|
| 1 | $AqBlMuCh-multi-dispense-buffer-chamfered-cover-pad-blowout10 | 4 : PBS and AR cyc… | 150 | A2 | 4:27:32 PM | 4:27:46 PM | Ok | 1 : Parhelia ST12 … | 150 | A1 | 4:27:46 PM | 4:28:08 PM | Ok |
| 2 | $AqBlMuCh-multi-dispense-buffer-chamfered-cover-pad-blowout10 | 4 : PBS and AR cyc… | 150 | A2 | 4:27:32 PM | 4:27:46 PM | Ok | 1 : Parhelia ST12 … | 150 | A1 | 4:27:46 PM | 4:28:08 PM | Ok |
| 1 | $AqBlMuCh-multi-dispense-buffer-chamfered-cover-pad-blowout10 | 4 : PBS and AR cyc… | 150 | A2 | 4:28:08 PM | 4:28:22 PM | Ok | 1 : Parhelia ST12 … | 150 | A1 | 4:28:22 PM | 4:28:42 PM | Ok |
| 2 | $AqBlMuCh-multi-dispense-buffer-chamfered-cover-pad-blowout10 | 8 : H2O2_Blocking_… | 120 | E2 | 4:28:08 PM | 4:28:22 PM | Ok | 1 : Parhelia ST12 … | 120 | A1 | 4:28:22 PM | 4:28:42 PM | Ok |

**31. Aluora Dye incubation step (10 min) [4:28:47 PM - 4:38:47 PM]**

**32. #2 3xWash and AR Buffer add [4:38:47 PM - 4:39:57 PM]**

| | | Aspirate | | | | | | Dispense | | | | | |
|---|---|---|---|---|---|---|---|---|---|---|---|---|---|
| Channel | Liquid Type | Labware | Volume | Well | Start (CET) | End (CET) | Status | Labware | Volume | Well | Start (CET) | End (CET) | Status |
| 1 | $Aqueous-buffer-chamfered-cover-pad-1 | 4 : PBS and AR cyc… | 150 | A2 | 4:38:52 PM | 4:39:05 PM | Ok | 1 : Parhelia ST12 … | 150 | A1 | 4:39:05 PM | 4:39:22 PM | Ok |
| 2 | $Aqueous-buffer-chamfered-cover-pad-1 | 4 : PBS and AR cyc… | 150 | A2 | 4:38:52 PM | 4:39:05 PM | Ok | 1 : Parhelia ST12 … | 150 | A1 | 4:39:05 PM | 4:39:22 PM | Ok |
| 1 | $Aqueous-buffer-chamfered-cover-pad-1 | 4 : PBS and AR cyc… | 150 | A2 | 4:39:22 PM | 4:39:35 PM | Ok | 1 : Parhelia ST12 … | 150 | A1 | 4:39:35 PM | 4:39:52 PM | Ok |
| 2 | $Aqueous-buffer-chamfered-cover-pad-1 | 4 : PBS and AR cyc… | 150 | B6 | 4:39:22 PM | 4:39:35 PM | Ok | 1 : Parhelia ST12 … | 150 | A1 | 4:39:35 PM | 4:39:52 PM | Ok |

**33. Heat to ER temp (90C) [4:39:57 PM - 4:51:42 PM]**

**34. #2 ER Topoff 1/2 (mixing delay 9 min) [4:51:42 PM - 5:01:18 PM]**

| | | Aspirate | | | | | | Dispense | | | | | |
|---|---|---|---|---|---|---|---|---|---|---|---|---|---|
| Channel | Liquid Type | Labware | Volume | Well | Start (CET) | End (CET) | Status | Labware | Volume | Well | Start (CET) | End (CET) | Status |
| 1 | $Aq30-BlMuCh-multi-dispense-buffer-chamfered-cover-pad-blowout | 4 : PBS and AR cyc… | 50 | B6 | 4:51:47 PM | 5:01:02 PM | Ok | 1 : Parhelia ST12 … | 50 | A1 | 5:01:02 PM | 5:01:13 PM | Ok |

**35. HHC Off [5:01:18 PM - 5:01:21 PM]**

**36. #2 ER Topoff 2/2 (mixing delay 8 min) [5:01:21 PM - 5:09:57 PM]**

| | | Aspirate | | | | | | Dispense | | | | | |
|---|---|---|---|---|---|---|---|---|---|---|---|---|---|
| Channel | Liquid Type | Labware | Volume | Well | Start (CET) | End (CET) | Status | Labware | Volume | Well | Start (CET) | End (CET) | Status |
| 1 | $Aq30-BlMuCh-multi-dispense-buffer-chamfered-cover-pad-blowout | 4 : PBS and AR cyc… | 50 | B6 | 5:01:26 PM | 5:09:40 PM | Ok | 1 : Parhelia ST12 … | 50 | A1 | 5:09:40 PM | 5:09:52 PM | Ok |

**37. Set temp to 20C [5:09:57 PM - 5:20:32 PM]**

**38. ———#3 3X PBS Wash + Blocking Buffer——— [5:20:32 PM - 5:21:52 PM]**

| | | Aspirate | | | | | | Dispense | | | | | |
|---|---|---|---|---|---|---|---|---|---|---|---|---|---|
| Channel | Liquid Type | Labware | Volume | Well | Start (CET) | End (CET) | Status | Labware | Volume | Well | Start (CET) | End (CET) | Status |
| 1 | $AqBlMuCh-multi-dispense-buffer-chamfered-cover-pad-blowout10 | 4 : PBS and AR cyc… | 150 | A3 | 5:20:37 PM | 5:20:51 PM | Ok | 1 : Parhelia ST12 … | 150 | A1 | 5:20:51 PM | 5:21:12 PM | Ok |
| 2 | $AqBlMuCh-multi-dispense-buffer-chamfered-cover-pad-blowout10 | 4 : PBS and AR cyc… | 150 | A3 | 5:20:37 PM | 5:20:51 PM | Ok | 1 : Parhelia ST12 … | 150 | A1 | 5:20:51 PM | 5:21:12 PM | Ok |
| 1 | $AqBlMuCh-multi-dispense-buffer-chamfered-cover-pad-blowout10 | 4 : PBS and AR cyc… | 150 | A3 | 5:21:12 PM | 5:21:27 PM | Ok | 1 : Parhelia ST12 … | 150 | A1 | 5:21:27 PM | 5:21:47 PM | Ok |
| 2 | $AqBlMuCh-multi-dispense-buffer-chamfered-cover-pad-blowout10 | 8 : H2O2_Blocking_… | 120 | B3 | 5:21:12 PM | 5:21:27 PM | Ok | 1 : Parhelia ST12 … | 120 | A1 | 5:21:27 PM | 5:21:47 PM | Ok |

**39. Blocking Buffer incubation (45 min) [5:21:52 PM - 6:06:52 PM]**

**40. #3 Add Primary antibody [6:06:52 PM - 6:07:16 PM]**

| Channel | Liquid Type | Aspirate Labware | Volume | Well | Start (CET) | End (CET) | Status | Dispense Labware | Volume | Well | Start (CET) | End (CET) | Status |
|---|---|---|---|---|---|---|---|---|---|---|---|---|---|
| 1 | $Aqueous-buffer-chamfered-cover-pad-1 | 8 : H2O2_Blocking_… | 120 | C3 | 6:06:58 PM | 6:07:04 PM | Ok | 1 : Parhelia ST12 … | 120 | A1 | 6:07:04 PM | 6:07:12 PM | Ok |

**41. Primary antibody incubation (60 min) [6:07:16 PM - 7:07:16 PM]**

**42. #3 3xWash and Secondary-HRP add [7:07:16 PM - 7:08:27 PM]**

| Channel | Liquid Type | Aspirate Labware | Volume | Well | Start (CET) | End (CET) | Status | Dispense Labware | Volume | Well | Start (CET) | End (CET) | Status |
|---|---|---|---|---|---|---|---|---|---|---|---|---|---|
| 1 | $AqBlMuCh-multi-dispense-buffer-chamfered-cover-pad-blowout10 | 4 : PBS and AR cyc… | 150 | A3 | 7:07:22 PM | 7:07:35 PM | Ok | 1 : Parhelia ST12 … | 150 | A1 | 7:07:35 PM | 7:07:52 PM | Ok |
| 2 | $AqBlMuCh-multi-dispense-buffer-chamfered-cover-pad-blowout10 | 4 : PBS and AR cyc… | 150 | A3 | 7:07:22 PM | 7:07:35 PM | Ok | 1 : Parhelia ST12 … | 150 | A1 | 7:07:35 PM | 7:07:52 PM | Ok |
| 1 | $AqBlMuCh-multi-dispense-buffer-chamfered-cover-pad-blowout10 | 4 : PBS and AR cyc… | 150 | A3 | 7:07:52 PM | 7:08:05 PM | Ok | 1 : Parhelia ST12 … | 150 | A1 | 7:08:05 PM | 7:08:22 PM | Ok |
| 2 | $AqBlMuCh-multi-dispense-buffer-chamfered-cover-pad-blowout10 | 8 : H2O2_Blocking_… | 120 | D3 | 7:07:52 PM | 7:08:05 PM | Ok | 1 : Parhelia ST12 … | 120 | A1 | 7:08:05 PM | 7:08:22 PM | Ok |

**43. Secondary-HRP incubation step (45 min) [7:08:27 PM - 7:43:27 PM]**

**44. #3 3xWash and Aluora dye [7:43:27 PM - 7:44:46 PM]**

| Channel | Liquid Type | Aspirate Labware | Volume | Well | Start (CET) | End (CET) | Status | Dispense Labware | Volume | Well | Start (CET) | End (CET) | Status |
|---|---|---|---|---|---|---|---|---|---|---|---|---|---|
| 1 | $AqBlMuCh-multi-dispense-buffer-chamfered-cover-pad-blowout10 | 4 : PBS and AR cyc… | 150 | A3 | 7:43:32 PM | 7:43:46 PM | Ok | 1 : Parhelia ST12 … | 150 | A1 | 7:43:46 PM | 7:44:07 PM | Ok |
| 2 | $AqBlMuCh-multi-dispense-buffer-chamfered-cover-pad-blowout10 | 4 : PBS and AR cyc… | 150 | A3 | 7:43:32 PM | 7:43:46 PM | Ok | 1 : Parhelia ST12 … | 150 | A1 | 7:43:46 PM | 7:44:07 PM | Ok |
| 1 | $AqBlMuCh-multi-dispense-buffer-chamfered-cover-pad-blowout10 | 4 : PBS and AR cyc… | 150 | A3 | 7:44:07 PM | 7:44:21 PM | Ok | 1 : Parhelia ST12 … | 150 | A1 | 7:44:21 PM | 7:44:41 PM | Ok |
| 2 | $AqBlMuCh-multi-dispense-buffer-chamfered-cover-pad-blowout10 | 8 : H2O2_Blocking_… | 120 | E3 | 7:44:07 PM | 7:44:21 PM | Ok | 1 : Parhelia ST12 … | 120 | A1 | 7:44:21 PM | 7:44:41 PM | Ok |

**45. Aluora Dye incubation step (10 min) [7:44:46 PM - 7:54:46 PM]**

**46. #3 3xWash and AR Buffer add [7:54:47 PM - 7:56:08 PM]**

| Channel | Liquid Type | Aspirate Labware | Volume | Well | Start (CET) | End (CET) | Status | Dispense Labware | Volume | Well | Start (CET) | End (CET) | Status |
|---|---|---|---|---|---|---|---|---|---|---|---|---|---|
| 1 | $AqBlMuCh-multi-dispense-buffer-chamfered-cover-pad-blowout10 | 4 : PBS and AR cyc… | 150 | A3 | 7:54:52 PM | 7:55:06 PM | Ok | 1 : Parhelia ST12 … | 150 | A1 | 7:55:06 PM | 7:55:27 PM | Ok |
| 2 | $AqBlMuCh-multi-dispense-buffer-chamfered-cover-pad-blowout10 | 4 : PBS and AR cyc… | 150 | A3 | 7:54:52 PM | 7:55:06 PM | Ok | 1 : Parhelia ST12 … | 150 | A1 | 7:55:06 PM | 7:55:27 PM | Ok |
| 1 | $AqBlMuCh-multi-dispense-buffer-chamfered-cover-pad-blowout10 | 4 : PBS and AR cyc… | 150 | A3 | 7:55:27 PM | 7:55:42 PM | Ok | 1 : Parhelia ST12 … | 150 | A1 | 7:55:42 PM | 7:56:03 PM | Ok |
| 2 | $AqBlMuCh-multi-dispense-buffer-chamfered-cover-pad-blowout10 | 4 : PBS and AR cyc… | 150 | C6 | 7:55:27 PM | 7:55:42 PM | Ok | 1 : Parhelia ST12 … | 150 | A1 | 7:55:42 PM | 7:56:03 PM | Ok |

**47. Heat to ER temp (90C) [7:56:08 PM - 8:07:56 PM]**

**48. #3 ER Topoff 1/2 (mixing delay 9 min) [8:07:56 PM - 8:17:33 PM]**

| Channel | Liquid Type | Aspirate Labware | Volume | Well | Start (CET) | End (CET) | Status | Dispense Labware | Volume | Well | Start (CET) | End (CET) | Status |
|---|---|---|---|---|---|---|---|---|---|---|---|---|---|

| | | | | | | | | | | | | | |
|---|---|---|---|---|---|---|---|---|---|---|---|---|---|
| 1 | $Aq30-BIMuCh-multi-dispense-buffer-chamfered-cover-pad-blowout | 4 : PBS and AR cyc… | 50 | C6 | 8:08:01 PM | 8:17:16 PM | Ok | 1 : Parhelia ST12 … | 50 | A1 | 8:17:16 PM | 8:17:28 PM | Ok |

**49. HHC Off [8:17:33 PM - 8:17:36 PM]**

**50. #3 ER Topoff 2/2 (mixing delay 8 min) [8:17:36 PM - 8:26:12 PM]**

| | | Aspirate | | | | | | Dispense | | | | | |
|---|---|---|---|---|---|---|---|---|---|---|---|---|---|
| Channel | Liquid Type | Labware | Volume | Well | Start (CET) | End (CET) | Status | Labware | Volume | Well | Start (CET) | End (CET) | Status |
| 1 | $Aq30-BIMuCh-multi-dispense-buffer-chamfered-cover-pad-blowout | 4 : PBS and AR cyc… | 50 | C6 | 8:17:41 PM | 8:25:55 PM | Ok | 1 : Parhelia ST12 … | 50 | A1 | 8:25:55 PM | 8:26:07 PM | Ok |

**51. Set temp to 20C [8:26:12 PM - 8:36:52 PM]**

**52. --------#4 3X PBS Wash + Blocking Buffer-------- [8:36:52 PM - 8:38:12 PM]**

| | | Aspirate | | | | | | Dispense | | | | | |
|---|---|---|---|---|---|---|---|---|---|---|---|---|---|
| Channel | Liquid Type | Labware | Volume | Well | Start (CET) | End (CET) | Status | Labware | Volume | Well | Start (CET) | End (CET) | Status |
| 1 | $AqBIMuCh-multi-dispense-buffer-chamfered-cover-pad-blowout10 | 4 : PBS and AR cyc… | 150 | A4 | 8:36:58 PM | 8:37:12 PM | Ok | 1 : Parhelia ST12 … | 150 | A1 | 8:37:12 PM | 8:37:33 PM | Ok |
| 2 | $AqBIMuCh-multi-dispense-buffer-chamfered-cover-pad-blowout10 | 4 : PBS and AR cyc… | 150 | A4 | 8:36:58 PM | 8:37:12 PM | Ok | 1 : Parhelia ST12 … | 150 | A1 | 8:37:12 PM | 8:37:33 PM | Ok |
| 1 | $AqBIMuCh-multi-dispense-buffer-chamfered-cover-pad-blowout10 | 4 : PBS and AR cyc… | 150 | A4 | 8:37:33 PM | 8:37:47 PM | Ok | 1 : Parhelia ST12 … | 150 | A1 | 8:37:47 PM | 8:38:07 PM | Ok |
| 2 | $AqBIMuCh-multi-dispense-buffer-chamfered-cover-pad-blowout10 | 8 : H2O2_Blocking_… | 120 | B4 | 8:37:33 PM | 8:37:47 PM | Ok | 1 : Parhelia ST12 … | 120 | A1 | 8:37:47 PM | 8:38:07 PM | Ok |

**53. Blocking Buffer incubation (45 min) [8:38:12 PM - 9:23:12 PM]**

**54. #4 Add Primary antibody [9:23:12 PM - 9:23:39 PM]**

| | | Aspirate | | | | | | Dispense | | | | | |
|---|---|---|---|---|---|---|---|---|---|---|---|---|---|
| Channel | Liquid Type | Labware | Volume | Well | Start (CET) | End (CET) | Status | Labware | Volume | Well | Start (CET) | End (CET) | Status |
| 1 | $Aq30-BIMuCh-multi-dispense-buffer-chamfered-cover-pad-blowout | 8 : H2O2_Blocking_… | 120 | C4 | 9:23:17 PM | 9:23:24 PM | Ok | 1 : Parhelia ST12 … | 120 | A1 | 9:23:24 PM | 9:23:34 PM | Ok |

**55. Primary antibody incubation (60 min) [9:23:39 PM - 10:23:39 PM]**

**56. #4 3xWash and Secondary-HRP add [10:23:39 PM - 10:24:49 PM]**

| | | Aspirate | | | | | | Dispense | | | | | |
|---|---|---|---|---|---|---|---|---|---|---|---|---|---|
| Channel | Liquid Type | Labware | Volume | Well | Start (CET) | End (CET) | Status | Labware | Volume | Well | Start (CET) | End (CET) | Status |
| 1 | $AqBIMuCh-multi-dispense-buffer-chamfered-cover-pad-blowout10 | 4 : PBS and AR cyc… | 150 | A4 | 10:23:44 PM | 10:23:57 PM | Ok | 1 : Parhelia ST12 … | 150 | A1 | 10:23:57 PM | 10:24:14 PM | Ok |
| 2 | $AqBIMuCh-multi-dispense-buffer-chamfered-cover-pad-blowout10 | 4 : PBS and AR cyc… | 150 | A4 | 10:23:44 PM | 10:23:57 PM | Ok | 1 : Parhelia ST12 … | 150 | A1 | 10:23:57 PM | 10:24:14 PM | Ok |
| 1 | $AqBIMuCh-multi-dispense-buffer-chamfered-cover-pad-blowout10 | 4 : PBS and AR cyc… | 150 | A4 | 10:24:14 PM | 10:24:28 PM | Ok | 1 : Parhelia ST12 … | 150 | A1 | 10:24:28 PM | 10:24:44 PM | Ok |
| 2 | $AqBIMuCh-multi-dispense-buffer-chamfered-cover-pad-blowout10 | 8 : H2O2_Blocking_… | 120 | D4 | 10:24:14 PM | 10:24:28 PM | Ok | 1 : Parhelia ST12 … | 120 | A1 | 10:24:28 PM | 10:24:44 PM | Ok |

**57. Secondary-HRP incubation step (45 min) [10:24:49 PM - 10:59:49 PM]**

**58. #4 3xWash and Aluora dye [10:59:49 PM - 11:01:09 PM]**

| | | Aspirate | | | | | | Dispense | | | | | |
|---|---|---|---|---|---|---|---|---|---|---|---|---|---|
| Channel | Liquid Type | Labware | Volume | Well | Start (CET) | End (CET) | Status | Labware | Volume | Well | Start (CET) | End (CET) | Status |

| Channel | Liquid Type | Aspirate Labware | Volume | Well | Start (CET) | End (CET) | Status | Dispense Labware | Volume | Well | Start (CET) | End (CET) | Status |
|---|---|---|---|---|---|---|---|---|---|---|---|---|---|
| 1 | $AqBIMuCh-multi-dispense-buffer-chamfered-cover-pad-blowout10 | 4 : PBS and AR cyc… | 150 | A4 | 10:59:55 PM | 11:00:09 PM | Ok | 1 : Parhelia ST12 … | 150 | A1 | 11:00:09 PM | 11:00:30 PM | Ok |
| 2 | $AqBIMuCh-multi-dispense-buffer-chamfered-cover-pad-blowout10 | 4 : PBS and AR cyc… | 150 | A4 | 10:59:55 PM | 11:00:09 PM | Ok | 1 : Parhelia ST12 … | 150 | A1 | 11:00:09 PM | 11:00:30 PM | Ok |
| 1 | $AqBIMuCh-multi-dispense-buffer-chamfered-cover-pad-blowout10 | 4 : PBS and AR cyc… | 150 | A4 | 11:00:30 PM | 11:00:44 PM | Ok | 1 : Parhelia ST12 … | 150 | A1 | 11:00:44 PM | 11:01:04 PM | Ok |
| 2 | $AqBIMuCh-multi-dispense-buffer-chamfered-cover-pad-blowout10 | 8 : H2O2_Blocking_… | 120 | E4 | 11:00:30 PM | 11:00:44 PM | Ok | 1 : Parhelia ST12 … | 120 | A1 | 11:00:44 PM | 11:01:04 PM | Ok |

**59. Aluora Dye incubation step (10 min) [11:01:09 PM - 11:11:09 PM]**

**60. #4 3xWash and AR Buffer add [11:11:09 PM - 11:12:30 PM]**

| | | Aspirate | | | | | | Dispense | | | | | |
|---|---|---|---|---|---|---|---|---|---|---|---|---|---|
| Channel | Liquid Type | Labware | Volume | Well | Start (CET) | End (CET) | Status | Labware | Volume | Well | Start (CET) | End (CET) | Status |
| 1 | $AqBIMuCh-multi-dispense-buffer-chamfered-cover-pad-blowout10 | 4 : PBS and AR cyc… | 150 | A4 | 11:11:14 PM | 11:11:28 PM | Ok | 1 : Parhelia ST12 … | 150 | A1 | 11:11:28 PM | 11:11:49 PM | Ok |
| 2 | $AqBIMuCh-multi-dispense-buffer-chamfered-cover-pad-blowout10 | 4 : PBS and AR cyc… | 150 | A4 | 11:11:14 PM | 11:11:28 PM | Ok | 1 : Parhelia ST12 … | 150 | A1 | 11:11:28 PM | 11:11:49 PM | Ok |
| 1 | $AqBIMuCh-multi-dispense-buffer-chamfered-cover-pad-blowout10 | 4 : PBS and AR cyc… | 150 | A4 | 11:11:49 PM | 11:12:04 PM | Ok | 1 : Parhelia ST12 … | 150 | A1 | 11:12:04 PM | 11:12:25 PM | Ok |
| 2 | $AqBIMuCh-multi-dispense-buffer-chamfered-cover-pad-blowout10 | 4 : PBS and AR cyc… | 150 | D6 | 11:11:49 PM | 11:12:04 PM | Ok | 1 : Parhelia ST12 … | 150 | A1 | 11:12:04 PM | 11:12:25 PM | Ok |

**61. Heat to ER temp (90C) [11:12:30 PM - 11:24:28 PM]**

**62. #4 ER Topoff 1/2 (mixing delay 9 min) [11:24:28 PM - 11:34:05 PM]**

| | | Aspirate | | | | | | Dispense | | | | | |
|---|---|---|---|---|---|---|---|---|---|---|---|---|---|
| Channel | Liquid Type | Labware | Volume | Well | Start (CET) | End (CET) | Status | Labware | Volume | Well | Start (CET) | End (CET) | Status |
| 1 | $Aq30-BIMuCh-multi-dispense-buffer-chamfered-cover-pad-blowout | 4 : PBS and AR cyc… | 50 | D6 | 11:24:33 PM | 11:33:48 PM | Ok | 1 : Parhelia ST12 … | 50 | A1 | 11:33:48 PM | 11:34:00 PM | Ok |

**63. HHC Off [11:34:06 PM - 11:34:09 PM]**

**64. #4 ER Topoff 2/2 (mixing delay 8 min) [11:34:09 PM - 11:42:45 PM]**

| | | Aspirate | | | | | | Dispense | | | | | |
|---|---|---|---|---|---|---|---|---|---|---|---|---|---|
| Channel | Liquid Type | Labware | Volume | Well | Start (CET) | End (CET) | Status | Labware | Volume | Well | Start (CET) | End (CET) | Status |
| 1 | $Aq30-BIMuCh-multi-dispense-buffer-chamfered-cover-pad-blowout | 4 : PBS and AR cyc… | 50 | D6 | 11:34:14 PM | 11:42:28 PM | Ok | 1 : Parhelia ST12 … | 50 | A1 | 11:42:28 PM | 11:42:40 PM | Ok |

**65. Set temp to 20C [11:42:45 PM - 11:53:46 PM]**

**66. ---------#5 3X PBS Wash + Blocking Buffer--------- [11:53:46 PM - 11:55:05 PM]**

| | | Aspirate | | | | | | Dispense | | | | | |
|---|---|---|---|---|---|---|---|---|---|---|---|---|---|
| Channel | Liquid Type | Labware | Volume | Well | Start (CET) | End (CET) | Status | Labware | Volume | Well | Start (CET) | End (CET) | Status |
| 1 | $AqBIMuCh-multi-dispense-buffer-chamfered-cover-pad-blowout10 | 3 : Plate | 150 | A1 | 11:53:52 PM | 11:54:05 PM | Ok | 1 : Parhelia ST12 … | 150 | A1 | 11:54:05 PM | 11:54:26 PM | Ok |
| 2 | $AqBIMuCh-multi-dispense-buffer-chamfered-cover-pad-blowout10 | 3 : Plate | 150 | A1 | 11:53:52 PM | 11:54:05 PM | Ok | 1 : Parhelia ST12 … | 150 | A1 | 11:54:05 PM | 11:54:26 PM | Ok |

| Channel | Liquid Type | Aspirate Labware | Volume | Well | Start (CET) | End (CET) | Status | Dispense Labware | Volume | Well | Start (CET) | End (CET) | Status |
|---|---|---|---|---|---|---|---|---|---|---|---|---|---|
| 1 | $AqBlMuCh-multi-dispense-buffer-chamfered-cover-pad-blowout10 | 3 : Plate | 150 | A1 | 11:54:26 PM | 11:54:40 PM | Ok | 1 : Parhelia ST12 … | 150 | A1 | 11:54:40 PM | 11:55:00 PM | Ok |
| 2 | $AqBlMuCh-multi-dispense-buffer-chamfered-cover-pad-blowout10 | 8 : H2O2_Blocking_… | 120 | B5 | 11:54:26 PM | 11:54:40 PM | Ok | 1 : Parhelia ST12 … | 120 | A1 | 11:54:40 PM | 11:55:00 PM | Ok |

**67. Blocking Buffer incubation (45 min) [11:55:05 PM - 12:40:05 AM]**

**68. #5 Add Primary antibody [12:40:05 AM - 12:40:32 AM]**

| | | Aspirate | | | | | | Dispense | | | | | |
|---|---|---|---|---|---|---|---|---|---|---|---|---|---|
| Channel | Liquid Type | Labware | Volume | Well | Start (CET) | End (CET) | Status | Labware | Volume | Well | Start (CET) | End (CET) | Status |
| 1 | $Aq30-BlMuCh-multi-dispense-buffer-chamfered-cover-pad-blowout | 8 : H2O2_Blocking_… | 120 | C5 | 12:40:10 AM | 12:40:17 AM | Ok | 1 : Parhelia ST12 … | 120 | A1 | 12:40:17 AM | 12:40:26 AM | Ok |

**69. Primary antibody incubation (60 min) [12:40:32 AM - 1:40:32 AM]**

**70. #5 3xWash and Secondary-HRP add [1:40:32 AM - 1:41:41 AM]**

| | | Aspirate | | | | | | Dispense | | | | | |
|---|---|---|---|---|---|---|---|---|---|---|---|---|---|
| Channel | Liquid Type | Labware | Volume | Well | Start (CET) | End (CET) | Status | Labware | Volume | Well | Start (CET) | End (CET) | Status |
| 1 | $AqBlMuCh-multi-dispense-buffer-chamfered-cover-pad-blowout10 | 3 : Plate | 150 | A1 | 1:40:37 AM | 1:40:50 AM | Ok | 1 : Parhelia ST12 … | 150 | A1 | 1:40:50 AM | 1:41:07 AM | Ok |
| 2 | $AqBlMuCh-multi-dispense-buffer-chamfered-cover-pad-blowout10 | 3 : Plate | 150 | A1 | 1:40:37 AM | 1:40:50 AM | Ok | 1 : Parhelia ST12 … | 150 | A1 | 1:40:50 AM | 1:41:07 AM | Ok |
| 1 | $AqBlMuCh-multi-dispense-buffer-chamfered-cover-pad-blowout10 | 3 : Plate | 150 | A1 | 1:41:07 AM | 1:41:20 AM | Ok | 1 : Parhelia ST12 … | 150 | A1 | 1:41:20 AM | 1:41:36 AM | Ok |
| 2 | $AqBlMuCh-multi-dispense-buffer-chamfered-cover-pad-blowout10 | 8 : H2O2_Blocking_… | 120 | D5 | 1:41:07 AM | 1:41:20 AM | Ok | 1 : Parhelia ST12 … | 120 | A1 | 1:41:20 AM | 1:41:36 AM | Ok |

**71. Secondary-HRP incubation step (45 min) [1:41:41 AM - 2:16:41 AM]**

**72. #5 3xWash and Aluora dye [2:16:41 AM - 2:17:59 AM]**

| | | Aspirate | | | | | | Dispense | | | | | |
|---|---|---|---|---|---|---|---|---|---|---|---|---|---|
| Channel | Liquid Type | Labware | Volume | Well | Start (CET) | End (CET) | Status | Labware | Volume | Well | Start (CET) | End (CET) | Status |
| 1 | $AqBlMuCh-multi-dispense-buffer-chamfered-cover-pad-blowout10 | 3 : Plate | 150 | A1 | 2:16:46 AM | 2:17:00 AM | Ok | 1 : Parhelia ST12 … | 150 | A1 | 2:17:00 AM | 2:17:20 AM | Ok |
| 2 | $AqBlMuCh-multi-dispense-buffer-chamfered-cover-pad-blowout10 | 3 : Plate | 150 | A1 | 2:16:46 AM | 2:17:00 AM | Ok | 1 : Parhelia ST12 … | 150 | A1 | 2:17:00 AM | 2:17:20 AM | Ok |
| 1 | $AqBlMuCh-multi-dispense-buffer-chamfered-cover-pad-blowout10 | 3 : Plate | 150 | A1 | 2:17:20 AM | 2:17:34 AM | Ok | 1 : Parhelia ST12 … | 150 | A1 | 2:17:34 AM | 2:17:54 AM | Ok |
| 2 | $AqBlMuCh-multi-dispense-buffer-chamfered-cover-pad-blowout10 | 8 : H2O2_Blocking_… | 120 | E5 | 2:17:20 AM | 2:17:34 AM | Ok | 1 : Parhelia ST12 … | 120 | A1 | 2:17:35 AM | 2:17:54 AM | Ok |

**73. Aluora Dye incubation step (10 min) [2:17:59 AM - 2:27:59 AM]**

**101. #7 3xWash [2:27:59 AM - 2:29:02 AM]**

| | | Aspirate | | | | | | Dispense | | | | | |
|---|---|---|---|---|---|---|---|---|---|---|---|---|---|
| Channel | Liquid Type | Labware | Volume | Well | Start (CET) | End (CET) | Status | Labware | Volume | Well | Start (CET) | End (CET) | Status |
| 1 | $AqBlMuCh-multi-dispense-buffer-chamfered-cover-pad-blowout10 | 3 : Plate | 150 | A3 | 2:28:04 AM | 2:28:18 AM | Ok | 1 : Parhelia ST12 … | 150 | A1 | 2:28:18 AM | 2:28:39 AM | Ok |
| 2 | $AqBlMuCh-multi-dispense-buffer-chamfered-cover-pad-blowout10 | 3 : Plate | 150 | A3 | 2:28:04 AM | 2:28:18 AM | Ok | 1 : Parhelia ST12 … | 150 | A1 | 2:28:18 AM | 2:28:39 AM | Ok |

| | | | | | | | | | | | | | |
|---|---|---|---|---|---|---|---|---|---|---|---|---|---|
| 1 | $AqBIMuCh-multi-dispense-buffer-chamfered-cover-pad-blowout10 | 3 : Plate | 150 | A3 | 2:28:39 AM | 2:28:47 AM | Ok | 1 : Parhelia ST12 … | 150 | A1 | 2:28:47 AM | 2:28:57 AM | Ok |

**103. Cool to 4C [2:29:02 AM - 2:40:05 AM]**

**104. Hold at 6C until user ends [2:40:05 AM - 8:32:30 AM]**

### Pauses

Pause 1: Wait Step Initiated Pause

- Length of pause: 2700 seconds
- Start of pause: Monday, February 23, 2026 10:50:09 AM
- End of pause: Monday, February 23, 2026 11:35:09 AM

Pause 2: Wait Step Initiated Pause

- Length of pause: 3600 seconds
- Start of pause: Monday, February 23, 2026 11:35:33 AM
- End of pause: Monday, February 23, 2026 12:35:33 PM

Pause 3: Wait Step Initiated Pause

- Length of pause: 2100 seconds
- Start of pause: Monday, February 23, 2026 12:36:53 PM
- End of pause: Monday, February 23, 2026 1:11:53 PM

Pause 4: Wait Step Initiated Pause

- Length of pause: 600 seconds
- Start of pause: Monday, February 23, 2026 1:13:12 PM
- End of pause: Monday, February 23, 2026 1:23:12 PM

Pause 5: Wait Step Initiated Pause

- Length of pause: 2700 seconds
- Start of pause: Monday, February 23, 2026 2:05:49 PM
- End of pause: Monday, February 23, 2026 2:50:49 PM

Pause 6: Wait Step Initiated Pause

- Length of pause: 3600 seconds
- Start of pause: Monday, February 23, 2026 2:51:17 PM
- End of pause: Monday, February 23, 2026 3:51:17 PM

Pause 7: Wait Step Initiated Pause

- Length of pause: 2100 seconds
- Start of pause: Monday, February 23, 2026 3:52:27 PM
- End of pause: Monday, February 23, 2026 4:27:27 PM

Pause 8: Wait Step Initiated Pause

- Length of pause: 600 seconds
- Start of pause: Monday, February 23, 2026 4:28:47 PM
- End of pause: Monday, February 23, 2026 4:38:47 PM

Pause 9: Wait Step Initiated Pause

- Length of pause: 2700 seconds
- Start of pause: Monday, February 23, 2026 5:21:52 PM
- End of pause: Monday, February 23, 2026 6:06:52 PM

Pause 10: Wait Step Initiated Pause

- Length of pause: 3600 seconds
- Start of pause: Monday, February 23, 2026 6:07:16 PM
- End of pause: Monday, February 23, 2026 7:07:16 PM

Pause 11: Wait Step Initiated Pause

- Length of pause: 2100 seconds
- Start of pause: Monday, February 23, 2026 7:08:27 PM
- End of pause: Monday, February 23, 2026 7:43:27 PM

Pause 12: Wait Step Initiated Pause

- Length of pause: 600 seconds
- Start of pause: Monday, February 23, 2026 7:44:47 PM
- End of pause: Monday, February 23, 2026 7:54:47 PM

Pause 13: Wait Step Initiated Pause

- Length of pause: 2700 seconds
- Start of pause: Monday, February 23, 2026 8:38:12 PM
- End of pause: Monday, February 23, 2026 9:23:12 PM

Pause 14: Wait Step Initiated Pause

- Length of pause: 3600 seconds
- Start of pause: Monday, February 23, 2026 9:23:39 PM
- End of pause: Monday, February 23, 2026 10:23:39 PM

Pause 15: Wait Step Initiated Pause

- Length of pause: 2100 seconds
- Start of pause: Monday, February 23, 2026 10:24:49 PM
- End of pause: Monday, February 23, 2026 10:59:49 PM

Pause 16: Wait Step Initiated Pause

- Length of pause: 600 seconds
- Start of pause: Monday, February 23, 2026 11:01:09 PM
- End of pause: Monday, February 23, 2026 11:11:09 PM

Pause 17: Wait Step Initiated Pause

- Length of pause: 2700 seconds
- Start of pause: Monday, February 23, 2026 11:55:05 PM
- End of pause: Tuesday, February 24, 2026 12:40:05 AM

Pause 18: Wait Step Initiated Pause

- Length of pause: 3600 seconds
- Start of pause: Tuesday, February 24, 2026 12:40:32 AM
- End of pause: Tuesday, February 24, 2026 1:40:32 AM

Pause 19: Wait Step Initiated Pause

- Length of pause: 2100 seconds
- Start of pause: Tuesday, February 24, 2026 1:41:41 AM
- End of pause: Tuesday, February 24, 2026 2:16:41 AM

Pause 20: Wait Step Initiated Pause

- Length of pause: 600 seconds
- Start of pause: Tuesday, February 24, 2026 2:17:59 AM
- End of pause: Tuesday, February 24, 2026 2:27:59 AM

Pause 21: Step Initiated Pause

- Length of pause: 21142 seconds
- Start of pause: Tuesday, February 24, 2026 2:40:07 AM
- End of pause: Tuesday, February 24, 2026 8:32:29 AM

### Door Events

[none]

### Error Messages

[none]

### Deck Snapshot

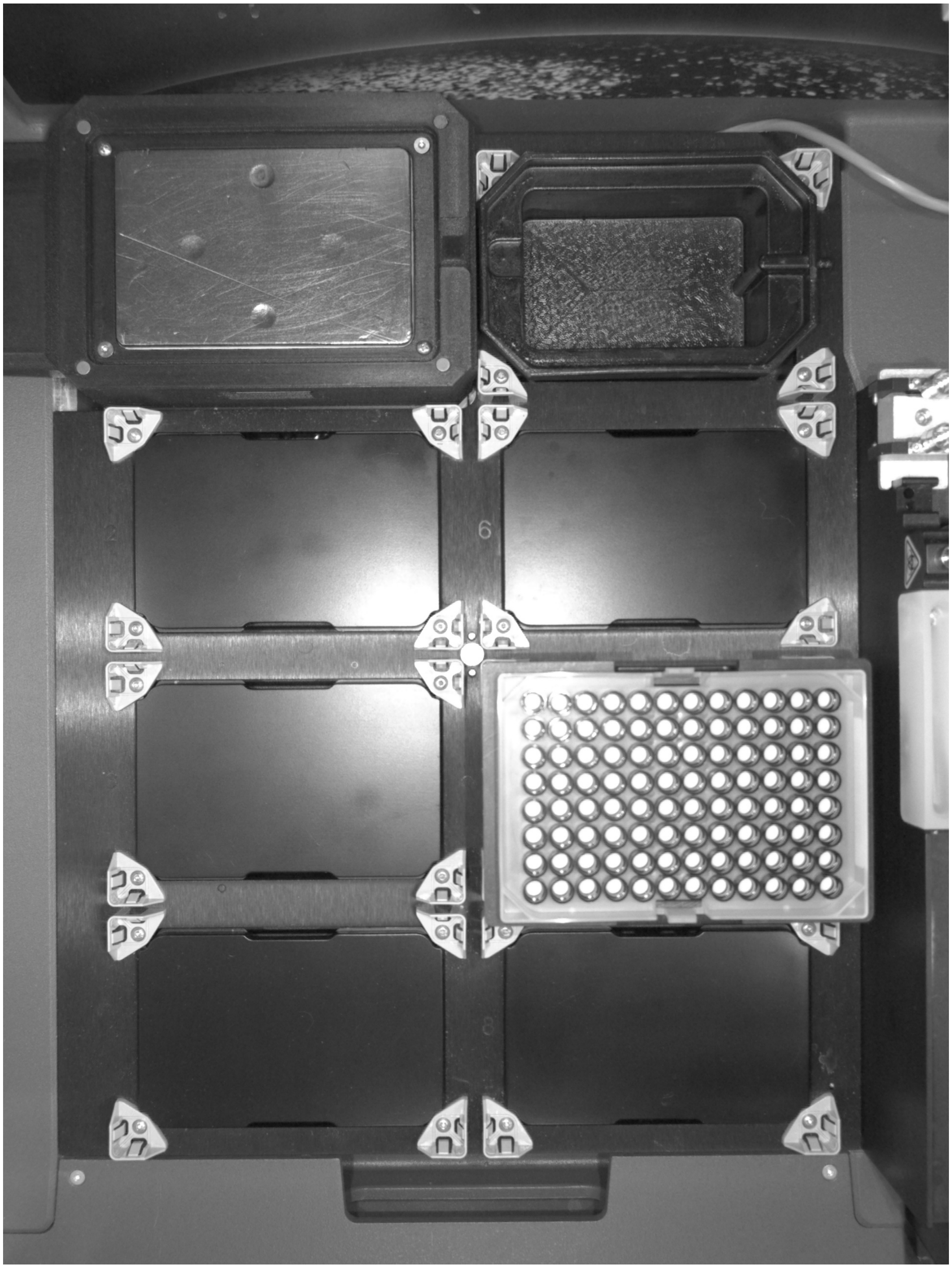

Signature: _______________________________

Operator: spu
