## Supplemental Figure 3 for "A workflow for combined detection of protein interactions and cell types for translational studies"

### Run Report

| | | | |
|---|---|---|---|
| Protocol Name: | isPLA PD1/PD-L1 HRP (LO) | Last Modified: | Feb 19, 2026 09:59:51 AM |
| Operator: | spu | Started on: | Feb 26, 2026 10:42:27 AM |
| Enclosure Status: | Present | Ended on: | Feb 27, 2026 09:08:04 AM |
| Serial Number: | PRPEM2207 | Software Version | 3.3.2.00005 |

### Completed

#### Deck Layout

| | |
|---|---|
| 1<br>Plate<br>Parhelia ST12 with Thermal Sheath<br>Hamilton Heater Cooler<br>Scan status: Incorrect (User confirmed) | 5<br>[empty]<br><br>Scan status: Correct |
| 2<br>[empty]<br><br>Scan status: Correct | 6<br>[empty]<br><br>Scan status: Correct |
| 3<br>Tip Rack<br>300 µL Framed Conductive No Filter<br><br>Scan status: ConfirmationRequired (User confirmed) | 7<br>Microtube Rack<br>Hamilton, 24, Microtube Rack, 6600409-01<br>See Protocol Tubes<br>Scan status: ConfirmationRequired (User confirmed) |
| 4<br>Reservoir<br>Seahorse-Agilent, Reservoir, V-Bottom, 12 Col, 252mL, Clear PP<br><br>Scan status: Correct | 8<br>[empty]<br><br>Scan status: Correct |

### Steps

○ Lighting Step

○ How Many Samples Step

1. 1. Apply Peroxide Block

   - Step Type: Add reagent
   - Pipetting Tool: TwoChannel
   - Liquid Name: peroxide
   - Aspirate Volume: 120
   - Dispense Volume: 120
   - Tips Name: 300 µL Framed Conductive No Filter
   - New tips for each transfer: False

2. Incubate - 30 min

   - Step Type: Pause
   - Duration: 1800 seconds

3. Wash - 1X TBST

   - Step Type: Add reagent
   - Pipetting Tool: TwoChannel
   - Liquid Name: 1X TBST
   - Aspirate Volume: 150
   - Dispense Volume: 150
   - Tips Name: 300 µL Framed Conductive No Filter
   - New tips for each transfer: False

4. Incubate - 1 min

   - Step Type: Pause
   - Duration: 60 seconds

5. Wash - 1X TBST

   - Step Type: Add reagent
   - Pipetting Tool: TwoChannel
   - Liquid Name: 1X TBST
   - Aspirate Volume: 150
   - Dispense Volume: 150
   - Tips Name: 300 µL Framed Conductive No Filter
   - New tips for each transfer: False

6. Incubate - 1 min

   - Step Type: Pause
   - Duration: 60 seconds

7. Wash - 1X TBST

   - Step Type: Add reagent
   - Pipetting Tool: TwoChannel
   - Liquid Name: 1X TBST
   - Aspirate Volume: 150
   - Dispense Volume: 150
   - Tips Name: 300 µL Framed Conductive No Filter
   - New tips for each transfer: False

8. Incubate - 1 min

   - Step Type: Pause
   - Duration: 60 seconds

9. Set temp to 38C

   - Step Type: Heat cool
   - Temperature: 38 °C

10. 2. Apply Blocking Solution

   - Step Type: Add reagent
   - Pipetting Tool: TwoChannel
   - Liquid Name: blocking solution
   - Aspirate Volume: 120
   - Dispense Volume: 120
   - Tips Name: 300 µL Framed Conductive No Filter
   - New tips for each transfer: False

11. Blocking incubation - 60 min

   - Step Type: Pause
   - Duration: 3600 seconds

12. Wash - 1X TBST

   - Step Type: Add reagent
   - Pipetting Tool: TwoChannel
   - Liquid Name: 1X TBST
   - Aspirate Volume: 150
   - Dispense Volume: 150
   - Tips Name: 300 µL Framed Conductive No Filter
   - New tips for each transfer: False

13. Incubate - 3 min

   - Step Type: Pause
   - Duration: 180 seconds

14. Wash - 1X TBST

- Step Type: Add reagent
- Pipetting Tool: TwoChannel
- Liquid Name: 1X TBST
- Aspirate Volume: 150
- Dispense Volume: 150
- Tips Name: 300 µL Framed Conductive No Filter
- New tips for each transfer: False

15. Incubate - 3 min

- Step Type: Pause
- Duration: 180 seconds

16. 3. Add neg control (PBS)

- Step Type: Add reagent
- Pipetting Tool: TwoChannel
- Liquid Name: neg control
- Aspirate Volume: 120
- Dispense Volume: 120
- Tips Name: 300 µL Framed Conductive No Filter
- New tips for each transfer: False

17. 4. Apply Antibody - PD1/PD-L1

- Step Type: Add reagent
- Pipetting Tool: OneChannel
- Liquid Name: Antibody
- Aspirate Volume: 120
- Dispense Volume: 120
- Tips Name: 300 µL Framed Conductive No Filter
- New tips for each transfer: False

18. Primary Ab incubation- 60 min

- Step Type: Pause
- Duration: 3600 seconds

19. Wash - 1X TBST

- Step Type: Add reagent
- Pipetting Tool: TwoChannel
- Liquid Name: 1X TBST
- Aspirate Volume: 150
- Dispense Volume: 150
- Tips Name: 300 µL Framed Conductive No Filter
- New tips for each transfer: False

20. Incubation - 2 min

- Step Type: Pause
- Duration: 120 seconds

21. Wash - 1X TBST

- Step Type: Add reagent
- Pipetting Tool: TwoChannel
- Liquid Name: 1X TBST
- Aspirate Volume: 150
- Dispense Volume: 150
- Tips Name: 300 µL Framed Conductive No Filter
- New tips for each transfer: False

22. Incubation - 2 min

- Step Type: Pause
- Duration: 120 seconds

23. Wash - 1X TBST

- Step Type: Add reagent
- Pipetting Tool: TwoChannel
- Liquid Name: 1X TBST
- Aspirate Volume: 150
- Dispense Volume: 150
- Tips Name: 300 µL Framed Conductive No Filter
- New tips for each transfer: False

24. Incubation - 2 min

- Step Type: Pause
- Duration: 120 seconds

25. 5. Apply Navenibody M1 and R2

- Step Type: Add reagent
- Pipetting Tool: OneChannel
- Liquid Name: Navenibody
- Aspirate Volume: 120
- Dispense Volume: 120
- Tips Name: 300 µL Framed Conductive No Filter
- New tips for each transfer: False

26. M1 and R2 incubation - 60 min

- Step Type: Pause
- Duration: 3540 seconds

27. 5. Apply Navenibody M1 and R2

- Step Type: Add reagent
- Pipetting Tool: OneChannel
- Liquid Name: Navenibody

- Aspirate Volume: 120
- Dispense Volume: 120
- Tips Name: 300 µL Framed Conductive No Filter
- New tips for each transfer: False

28. M1 and R2 incubation - 60 min
    - Step Type: Pause
    - Duration: 3540 seconds
29. Wash - 1X TBST
    - Step Type: Add reagent
    - Pipetting Tool: TwoChannel
    - Liquid Name: 1X TBST
    - Aspirate Volume: 150
    - Dispense Volume: 150
    - Tips Name: 300 µL Framed Conductive No Filter
    - New tips for each transfer: False
30. Wash - 1X TBST
    - Step Type: Add reagent
    - Pipetting Tool: TwoChannel
    - Liquid Name: 1X TBST
    - Aspirate Volume: 150
    - Dispense Volume: 150
    - Tips Name: 300 µL Framed Conductive No Filter
    - New tips for each transfer: False
31. Wash - 1X TBST
    - Step Type: Add reagent
    - Pipetting Tool: TwoChannel
    - Liquid Name: 1X TBST
    - Aspirate Volume: 150
    - Dispense Volume: 150
    - Tips Name: 300 µL Framed Conductive No Filter
    - New tips for each transfer: False
32. Incubate - 2 min
    - Step Type: Pause
    - Duration: 120 seconds
33. Wash - 1X TBST
    - Step Type: Add reagent
    - Pipetting Tool: TwoChannel
    - Liquid Name: 1X TBST
    - Aspirate Volume: 150
    - Dispense Volume: 150
    - Tips Name: 300 µL Framed Conductive No Filter
    - New tips for each transfer: False
34. Incubate - 2 min
    - Step Type: Pause
    - Duration: 120 seconds
35. Wash - 1X TBST
    - Step Type: Add reagent
    - Pipetting Tool: TwoChannel
    - Liquid Name: 1X TBST
    - Aspirate Volume: 150
    - Dispense Volume: 150
    - Tips Name: 300 µL Framed Conductive No Filter
    - New tips for each transfer: False
36. Incubate - 20 min
    - Step Type: Pause
    - Duration: 1200 seconds
37. Wash - 1X TBST
    - Step Type: Add reagent
    - Pipetting Tool: TwoChannel
    - Liquid Name: 1X TBST
    - Aspirate Volume: 150
    - Dispense Volume: 150
    - Tips Name: 300 µL Framed Conductive No Filter
    - New tips for each transfer: False
38. Wash - 1X TBST
    - Step Type: Add reagent
    - Pipetting Tool: TwoChannel
    - Liquid Name: 1X TBST
    - Aspirate Volume: 150
    - Dispense Volume: 150
    - Tips Name: 300 µL Framed Conductive No Filter
    - New tips for each transfer: False
39. Incubate - 20 min
    - Step Type: Pause
    - Duration: 1200 seconds
40. Wash - 1X TBST
    - Step Type: Add reagent

- Pipetting Tool: TwoChannel
- Liquid Name: 1X TBST
- Aspirate Volume: 150
- Dispense Volume: 150
- Tips Name: 300 µL Framed Conductive No Filter
- New tips for each transfer: False

41. Wash - 1X TBST

- Step Type: Add reagent
- Pipetting Tool: TwoChannel
- Liquid Name: 1X TBST
- Aspirate Volume: 150
- Dispense Volume: 150
- Tips Name: 300 µL Framed Conductive No Filter
- New tips for each transfer: False

42. Wash - 1X TBST

- Step Type: Add reagent
- Pipetting Tool: TwoChannel
- Liquid Name: 1X TBST
- Aspirate Volume: 150
- Dispense Volume: 150
- Tips Name: 300 µL Framed Conductive No Filter
- New tips for each transfer: False

43. 6. Apply Reaction 1

- Step Type: Add reagent
- Pipetting Tool: TwoChannel
- Liquid Name: Reaction 1
- Aspirate Volume: 120
- Dispense Volume: 120
- Tips Name: 300 µL Framed Conductive No Filter
- New tips for each transfer: False

44. Reaction 1 incubatiom - 30 min

- Step Type: Pause
- Duration: 1800 seconds

45. Wash - 1X TBST

- Step Type: Add reagent
- Pipetting Tool: TwoChannel
- Liquid Name: 1X TBST
- Aspirate Volume: 150
- Dispense Volume: 150
- Tips Name: 300 µL Framed Conductive No Filter
- New tips for each transfer: False

46. Incubation - 2 min

- Step Type: Pause
- Duration: 120 seconds

47. Wash - 1X TBST

- Step Type: Add reagent
- Pipetting Tool: TwoChannel
- Liquid Name: 1X TBST
- Aspirate Volume: 150
- Dispense Volume: 150
- Tips Name: 300 µL Framed Conductive No Filter
- New tips for each transfer: False

48. Incubation - 2 min

- Step Type: Pause
- Duration: 120 seconds

49. Heat/Cool Step 36 C

- Step Type: Heat cool
- Temperature: 36 °C

50. 7. Apply Reaction 2

- Step Type: Add reagent
- Pipetting Tool: TwoChannel
- Liquid Name: Reaction 2
- Aspirate Volume: 120
- Dispense Volume: 120
- Tips Name: 300 µL Framed Conductive No Filter
- New tips for each transfer: False

51. R2 amplification - 45 min

- Step Type: Pause
- Duration: 2700 seconds

52. 7. Apply Reaction 2

- Step Type: Add reagent
- Pipetting Tool: TwoChannel
- Liquid Name: Reaction 2
- Aspirate Volume: 120
- Dispense Volume: 120
- Tips Name: 300 µL Framed Conductive No Filter
- New tips for each transfer: False

53. R2 amplification - 45 min

- Step Type: Pause
- Duration: 2700 seconds

54. Wash - 1X TBS

- Step Type: Add reagent
- Pipetting Tool: TwoChannel
- Liquid Name: tbs
- Aspirate Volume: 150
- Dispense Volume: 150
- Tips Name: 300 µL Framed Conductive No Filter
- New tips for each transfer: False

55. Incubation - 5 min

- Step Type: Pause
- Duration: 300 seconds

56. Wash - 1X TBS

- Step Type: Add reagent
- Pipetting Tool: TwoChannel
- Liquid Name: tbs
- Aspirate Volume: 150
- Dispense Volume: 150
- Tips Name: 300 µL Framed Conductive No Filter
- New tips for each transfer: False

57. Incubation - 5 min

- Step Type: Pause
- Duration: 300 seconds

58. Wash - 1X TBS 0.1

- Step Type: Add reagent
- Pipetting Tool: TwoChannel
- Liquid Name: tbs 0.1
- Aspirate Volume: 150
- Dispense Volume: 150
- Tips Name: 300 µL Framed Conductive No Filter
- New tips for each transfer: False

59. Incubation - 5 min

- Step Type: Pause
- Duration: 300 seconds

60. Cool to RT

- Step Type: Heat cool
- Temperature: 23 °C

61. Wait - 5 min equilibration

- Step Type: Pause
- Duration: 300 seconds

62. 8. Apply HRP

- Step Type: Add reagent
- Pipetting Tool: TwoChannel
- Liquid Name: HRP
- Aspirate Volume: 120
- Dispense Volume: 120
- Tips Name: 300 µL Framed Conductive No Filter
- New tips for each transfer: False

63. HRP incubation - 30 min

- Step Type: Pause
- Duration: 1800 seconds

64. Wash - 1X TBS

- Step Type: Add reagent
- Pipetting Tool: TwoChannel
- Liquid Name: tbs
- Aspirate Volume: 150
- Dispense Volume: 150
- Tips Name: 300 µL Framed Conductive No Filter
- New tips for each transfer: False

65. Incubation - 2 min

- Step Type: Pause
- Duration: 120 seconds

66. Wash - 1X TBS

- Step Type: Add reagent
- Pipetting Tool: TwoChannel
- Liquid Name: tbs
- Aspirate Volume: 150
- Dispense Volume: 150
- Tips Name: 300 µL Framed Conductive No Filter
- New tips for each transfer: False

67. Incubation - 2 min

- Step Type: Pause
- Duration: 120 seconds

68. Wash - PBS

- Step Type: Add reagent
- Pipetting Tool: TwoChannel
- Liquid Name: PBS
- Aspirate Volume: 150
- Dispense Volume: 150
- Tips Name: 300 µL Framed Conductive No Filter
- New tips for each transfer: False

69. 9. Apply Detection aluora

- Step Type: Add reagent
- Pipetting Tool: TwoChannel
- Liquid Name: Detection aluora
- Aspirate Volume: 120
- Dispense Volume: 120
- Tips Name: 300 µL Framed Conductive No Filter
- New tips for each transfer: False

70. Aluora incubation - 10 min

- Step Type: Pause
- Duration: 600 seconds

71. Wash - PBS

- Step Type: Add reagent
- Pipetting Tool: TwoChannel
- Liquid Name: PBS
- Aspirate Volume: 150
- Dispense Volume: 150
- Tips Name: 300 µL Framed Conductive No Filter
- New tips for each transfer: False

72. Incubation - 2 min

- Step Type: Pause
- Duration: 120 seconds

73. Wash - PBS

- Step Type: Add reagent
- Pipetting Tool: TwoChannel
- Liquid Name: PBS
- Aspirate Volume: 150
- Dispense Volume: 150
- Tips Name: 300 µL Framed Conductive No Filter
- New tips for each transfer: False

74. Incubation - 2 min

- Step Type: Pause
- Duration: 120 seconds

75. 9. Apply DAPI

- Step Type: Add reagent
- Pipetting Tool: TwoChannel
- Liquid Name: DAPI
- Aspirate Volume: 120
- Dispense Volume: 120
- Tips Name: 300 µL Framed Conductive No Filter
- New tips for each transfer: False

76. DAPI incubation- 4 min

- Step Type: Pause
- Duration: 240 seconds

77. 9. Apply DAPI

- Step Type: Add reagent
- Pipetting Tool: TwoChannel
- Liquid Name: DAPI
- Aspirate Volume: 120
- Dispense Volume: 120
- Tips Name: 300 µL Framed Conductive No Filter
- New tips for each transfer: False

78. DAPI incubation- 4 min

- Step Type: Pause
- Duration: 240 seconds

79. Wash - PBS

- Step Type: Add reagent
- Pipetting Tool: TwoChannel
- Liquid Name: PBS
- Aspirate Volume: 150
- Dispense Volume: 150
- Tips Name: 300 µL Framed Conductive No Filter
- New tips for each transfer: False

80. Wait - 2 min

- Step Type: Pause
- Duration: 120 seconds

81. Wash - PBS

- Step Type: Add reagent
- Pipetting Tool: TwoChannel
- Liquid Name: PBS

- Aspirate Volume: 150
- Dispense Volume: 150
- Tips Name: 300 µL Framed Conductive No Filter
- New tips for each transfer: False

82. 1X TBST

- Step Type: Add reagent
- Pipetting Tool: TwoChannel
- Liquid Name: 1X TBST
- Aspirate Volume: 150
- Dispense Volume: 150
- Tips Name: 300 µL Framed Conductive No Filter
- New tips for each transfer: False

83. 1X TBST

- Step Type: Add reagent
- Pipetting Tool: TwoChannel
- Liquid Name: 1X TBST
- Aspirate Volume: 150
- Dispense Volume: 150
- Tips Name: 300 µL Framed Conductive No Filter
- New tips for each transfer: False

84. Wait - 2 min

- Step Type: Pause
- Duration: 120 seconds

85. 0.1X TBST

- Step Type: Add reagent
- Pipetting Tool: TwoChannel
- Liquid Name: 0.1X TBST
- Aspirate Volume: 150
- Dispense Volume: 150
- Tips Name: 300 µL Framed Conductive No Filter
- New tips for each transfer: False

86. 0.1X TBST

- Step Type: Add reagent
- Pipetting Tool: TwoChannel
- Liquid Name: 0.1X TBST
- Aspirate Volume: 150
- Dispense Volume: 150
- Tips Name: 300 µL Framed Conductive No Filter
- New tips for each transfer: False

87. Wash - PBS

- Step Type: Add reagent
- Pipetting Tool: TwoChannel
- Liquid Name: PBS
- Aspirate Volume: 150
- Dispense Volume: 150
- Tips Name: 300 µL Framed Conductive No Filter
- New tips for each transfer: False

88. Cool to 4C

- Step Type: Heat cool
- Temperature: 4 °C

89. Wait until user resumes

- Step Type: Pause
- Duration: Pause until user resumes run.

### Liquids

| Key | Name | Type | Location(s) |
|---|---|---|---|
| Liquid 1 | PBS | $Aqueous-buffer-chamfered-cover-pad-1 | Site 4 - A8; |
| Liquid 14 | Detection aluora | $Aqueous-buffer-chamfered-cover-pad-1 | Site 7 - C3; |
| Liquid 15 | DAPI | $Aqueous-multi-dispense-buffer-chamfered-cover-pad-1 | Site 7 - D3; |
| Liquid 2 | tbs 0.1 | $Aqueous-buffer-chamfered-cover-pad-1 | Site 4 - A3; |
| Liquid 3 | tbs | Aqueous | Site 4 - A4; |
| Liquid 5 | 1X TBST | $AqBlMuCh-multi-dispense-buffer-chamfered-cover-pad-blowout10 | Site 4 - A5; Site 4 - A6; |
| Liquid 6 | peroxide | $Aqueous-buffer-chamfered-cover-pad-1 | Site 7 - A5; |
| Liquid 7 | blocking solution | $Aqueous-multi-dispense-buffer-chamfered-cover-pad-1 | Site 7 - A1; |
| Liquid 4 | 0.1X TBST | $Aqueous-multi-dispense-buffer-chamfered-cover-pad-1 | Site 4 - A7; |
| Liquid 9 | Antibody | $Aqueous-buffer-chamfered-cover-pad-1 | Site 7 - B1; |
| Liquid 10 | Navenibody | $Aqueous-buffer-chamfered-cover-pad-1 | Site 7 - C1; |
| Liquid 11 | Reaction 1 | $Aqueous-multi-dispense-buffer-chamfered-cover-pad-1 | Site 7 - D1; |
| Liquid 12 | Reaction 2 | $Aqueous-buffer-chamfered-cover-pad-1 | Site 7 - A3; |
| Liquid 13 | HRP | $Aqueous-buffer-chamfered-cover-pad-1 | Site 7 - B3; |
| Liquid 8 | neg control | Aqueous | Site 7 - B5; |

### Tubes

| Labware Name | Site | Tube Name | Location(s) |
|---|---|---|---|
| Microtube Rack | 7 | Eppendorf, Safe-Lock Tube, 1.5 mL | A1; A3; A5; B1; B3; B5; C1; C3; D1; D3; |

### Protocol Setup

○ Lighting

- Deck Lights: Always Enable
- Error Lighting Disabled: No

○ How Many Samples Step

- Step Type: How many samples
- Message: how many samples
- Deck Positions: 1:Plate
- Selected Targets
  - Plate 1: A1, B1

### Activity

**1. 1. Apply Peroxide Block [10:42:23 AM - 10:42:51 AM]**

| | | Aspirate | | | | | | Dispense | | | | | |
|---|---|---|---|---|---|---|---|---|---|---|---|---|---|
| Channel | Liquid Type | Labware | Volume | Well | Start (CET) | End (CET) | Status | Labware | Volume | Well | Start (CET) | End (CET) | Status |
| 1 | $Aqueous-buffer-chamfered-cover-pad-1 | 7 : Microtube Rack | 120 | A5 | 10:42:26 AM | 10:42:38 AM | Ok | 1 : Plate | 120 | A1 | 10:42:38 AM | 10:42:45 AM | Ok |
| 2 | $Aqueous-buffer-chamfered-cover-pad-1 | 7 : Microtube Rack | 120 | A5 | 10:42:26 AM | 10:42:38 AM | Ok | 1 : Plate | 120 | B1 | 10:42:38 AM | 10:42:45 AM | Ok |

**2. Incubate - 30 min [10:42:51 AM - 11:12:51 AM]**

**3. Wash - 1X TBST [11:12:51 AM - 11:13:17 AM]**

| | | Aspirate | | | | | | Dispense | | | | | |
|---|---|---|---|---|---|---|---|---|---|---|---|---|---|
| Channel | Liquid Type | Labware | Volume | Well | Start (CET) | End (CET) | Status | Labware | Volume | Well | Start (CET) | End (CET) | Status |
| 1 | $AqBlMuCh-multi-dispense-buffer-chamfered-cover-pad-blowout10 | 4 : Reservoir | 150 | A5 | 11:12:56 AM | 11:13:04 AM | Ok | 1 : Plate | 150 | A1 | 11:13:04 AM | 11:13:13 AM | Ok |
| 2 | $AqBlMuCh-multi-dispense-buffer-chamfered-cover-pad-blowout10 | 4 : Reservoir | 150 | A5 | 11:12:56 AM | 11:13:04 AM | Ok | 1 : Plate | 150 | B1 | 11:13:04 AM | 11:13:13 AM | Ok |

**4. Incubate - 1 min [11:13:17 AM - 11:14:17 AM]**

**5. Wash - 1X TBST [11:14:17 AM - 11:14:44 AM]**

| | | Aspirate | | | | | | Dispense | | | | | |
|---|---|---|---|---|---|---|---|---|---|---|---|---|---|
| Channel | Liquid Type | Labware | Volume | Well | Start (CET) | End (CET) | Status | Labware | Volume | Well | Start (CET) | End (CET) | Status |
| 1 | $AqBlMuCh-multi-dispense-buffer-chamfered-cover-pad-blowout10 | 4 : Reservoir | 150 | A5 | 11:14:22 AM | 11:14:30 AM | Ok | 1 : Plate | 150 | A1 | 11:14:30 AM | 11:14:39 AM | Ok |
| 2 | $AqBlMuCh-multi-dispense-buffer-chamfered-cover-pad-blowout10 | 4 : Reservoir | 150 | A5 | 11:14:22 AM | 11:14:30 AM | Ok | 1 : Plate | 150 | B1 | 11:14:30 AM | 11:14:39 AM | Ok |

**6. Incubate - 1 min [11:14:44 AM - 11:15:44 AM]**

**7. Wash - 1X TBST [11:15:44 AM - 11:16:11 AM]**

| | | Aspirate | | | | | | Dispense | | | | | |
|---|---|---|---|---|---|---|---|---|---|---|---|---|---|
| Channel | Liquid Type | Labware | Volume | Well | Start (CET) | End (CET) | Status | Labware | Volume | Well | Start (CET) | End (CET) | Status |
| 1 | $AqBlMuCh-multi-dispense-buffer-chamfered-cover-pad-blowout10 | 4 : Reservoir | 150 | A5 | 11:15:49 AM | 11:15:57 AM | Ok | 1 : Plate | 150 | A1 | 11:15:57 AM | 11:16:06 AM | Ok |
| 2 | $AqBlMuCh-multi-dispense-buffer-chamfered-cover-pad-blowout10 | 4 : Reservoir | 150 | A5 | 11:15:49 AM | 11:15:57 AM | Ok | 1 : Plate | 150 | B1 | 11:15:57 AM | 11:16:06 AM | Ok |

**8. Incubate - 1 min [11:16:11 AM - 11:17:11 AM]**

**9. Set temp to 38C [11:17:11 AM - 11:17:48 AM]**

**10. 2. Apply Blocking Solution [11:17:48 AM - 11:18:21 AM]**

| Channel | Liquid Type | Aspirate<br>Labware | Volume | Well | Start (CET) | End (CET) | Status | Dispense<br>Labware | Volume | Well | Start (CET) | End (CET) | Status |
|---|---|---|---|---|---|---|---|---|---|---|---|---|---|
| 1 | $Aqueous-multi-dispense-buffer-chamfered-cover-pad-1 | 7 : Microtube Rack | 120 | A1 | 11:17:54 AM | 11:18:07 AM | Ok | 1 : Plate | 120 | A1 | 11:18:07 AM | 11:18:16 AM | Ok |
| 2 | $Aqueous-multi-dispense-buffer-chamfered-cover-pad-1 | 7 : Microtube Rack | 120 | A1 | 11:17:54 AM | 11:18:07 AM | Ok | 1 : Plate | 120 | B1 | 11:18:07 AM | 11:18:16 AM | Ok |

**11. Blocking incubation - 60 min [11:18:21 AM - 12:18:21 PM]**

**12. Wash - 1X TBST [12:18:21 PM - 12:18:47 PM]**

| Channel | Liquid Type | Aspirate<br>Labware | Volume | Well | Start (CET) | End (CET) | Status | Dispense<br>Labware | Volume | Well | Start (CET) | End (CET) | Status |
|---|---|---|---|---|---|---|---|---|---|---|---|---|---|
| 1 | $AqBlMuCh-multi-dispense-buffer-chamfered-cover-pad-blowout10 | 4 : Reservoir | 150 | A5 | 12:18:26 PM | 12:18:34 PM | Ok | 1 : Plate | 150 | A1 | 12:18:34 PM | 12:18:43 PM | Ok |
| 2 | $AqBlMuCh-multi-dispense-buffer-chamfered-cover-pad-blowout10 | 4 : Reservoir | 150 | A5 | 12:18:26 PM | 12:18:34 PM | Ok | 1 : Plate | 150 | B1 | 12:18:34 PM | 12:18:43 PM | Ok |

**13. Incubate - 3 min [12:18:47 PM - 12:21:47 PM]**

**14. Wash - 1X TBST [12:21:47 PM - 12:22:15 PM]**

| Channel | Liquid Type | Aspirate<br>Labware | Volume | Well | Start (CET) | End (CET) | Status | Dispense<br>Labware | Volume | Well | Start (CET) | End (CET) | Status |
|---|---|---|---|---|---|---|---|---|---|---|---|---|---|
| 1 | $AqBlMuCh-multi-dispense-buffer-chamfered-cover-pad-blowout10 | 4 : Reservoir | 150 | A5 | 12:21:53 PM | 12:22:01 PM | Ok | 1 : Plate | 150 | A1 | 12:22:01 PM | 12:22:09 PM | Ok |
| 2 | $AqBlMuCh-multi-dispense-buffer-chamfered-cover-pad-blowout10 | 4 : Reservoir | 150 | A5 | 12:21:53 PM | 12:22:01 PM | Ok | 1 : Plate | 150 | B1 | 12:22:01 PM | 12:22:09 PM | Ok |

**15. Incubate - 3 min [12:22:15 PM - 12:25:15 PM]**

**16. 3. Add neg control (PBS) [12:25:15 PM - 12:25:45 PM]**

| Channel | Liquid Type | Aspirate<br>Labware | Volume | Well | Start (CET) | End (CET) | Status | Dispense<br>Labware | Volume | Well | Start (CET) | End (CET) | Status |
|---|---|---|---|---|---|---|---|---|---|---|---|---|---|
| 1 | Aqueous | 7 : Microtube Rack | 160 | B5 | 12:25:20 PM | 12:25:29 PM | Ok | 7 : Microtube Rack | 20 | B5 | 12:25:29 PM | 12:25:32 PM | Ok |
| | | | | | | | | 1 : Plate | 120 | B1 | 12:25:32 PM | 12:25:37 PM | Ok |
| | | | | | | | | 7 : Microtube Rack | 20 | B5 | 12:25:37 PM | 12:25:41 PM | Ok |

**17. 4. Apply Antibody - PD1/PD-L1 [12:25:45 PM - 12:26:09 PM]**

| Channel | Liquid Type | Aspirate<br>Labware | Volume | Well | Start (CET) | End (CET) | Status | Dispense<br>Labware | Volume | Well | Start (CET) | End (CET) | Status |
|---|---|---|---|---|---|---|---|---|---|---|---|---|---|
| 2 | $Aqueous-buffer-chamfered-cover-pad-1 | 7 : Microtube Rack | 120 | B1 | 12:25:50 PM | 12:25:57 PM | Ok | 1 : Plate | 120 | A1 | 12:25:57 PM | 12:26:04 PM | Ok |

**18. Primary Ab incubation- 60 min [12:26:09 PM - 1:26:09 PM]**

**19. Wash - 1X TBST [1:26:09 PM - 1:26:37 PM]**

| Channel | Liquid Type | Aspirate<br>Labware | Volume | Well | Start (CET) | End (CET) | Status | Dispense<br>Labware | Volume | Well | Start (CET) | End (CET) | Status |
|---|---|---|---|---|---|---|---|---|---|---|---|---|---|
| 1 | $AqBlMuCh-multi-dispense-buffer-chamfered-cover-pad-blowout10 | 4 : Reservoir | 150 | A5 | 1:26:15 PM | 1:26:22 PM | Ok | 1 : Plate | 150 | A1 | 1:26:22 PM | 1:26:31 PM | Ok |
| 2 | $AqBlMuCh-multi-dispense-buffer-chamfered-cover-pad-blowout10 | 4 : Reservoir | 150 | A5 | 1:26:15 PM | 1:26:22 PM | Ok | 1 : Plate | 150 | B1 | 1:26:22 PM | 1:26:31 PM | Ok |

**20. Incubation - 2 min [1:26:37 PM - 1:28:37 PM]**

**21. Wash - 1X TBST [1:28:37 PM - 1:29:03 PM]**

| Channel | Liquid Type | Aspirate<br>Labware | Volume | Well | Start (CET) | End (CET) | Status | Dispense<br>Labware | Volume | Well | Start (CET) | End (CET) | Status |
|---|---|---|---|---|---|---|---|---|---|---|---|---|---|
| 1 | $AqBlMuCh-multi-dispense-buffer-chamfered-cover-pad-blowout10 | 4 : Reservoir | 150 | A5 | 1:28:42 PM | 1:28:50 PM | Ok | 1 : Plate | 150 | A1 | 1:28:50 PM | 1:28:59 PM | Ok |

| Channel | Liquid Type | Aspirate Labware | Volume | Well | Start (CET) | End (CET) | Status | Dispense Labware | Volume | Well | Start (CET) | End (CET) | Status |
|---|---|---|---|---|---|---|---|---|---|---|---|---|---|
| 2 | $AqBIMuCh-multi-dispense-buffer-chamfered-cover-pad-blowout10 | 4 : Reservoir | 150 | A5 | 1:28:42 PM | 1:28:50 PM | Ok | 1 : Plate | 150 | B1 | 1:28:50 PM | 1:28:59 PM | Ok |

**22. Incubation - 2 min [1:29:03 PM - 1:31:03 PM]**

**23. Wash - 1X TBST [1:31:03 PM - 1:31:30 PM]**

| | | **Aspirate** | | | | | | **Dispense** | | | | | |
|---|---|---|---|---|---|---|---|---|---|---|---|---|---|
| Channel | Liquid Type | Labware | Volume | Well | Start (CET) | End (CET) | Status | Labware | Volume | Well | Start (CET) | End (CET) | Status |
| 1 | $AqBIMuCh-multi-dispense-buffer-chamfered-cover-pad-blowout10 | 4 : Reservoir | 150 | A5 | 1:31:09 PM | 1:31:16 PM | Ok | 1 : Plate | 150 | A1 | 1:31:16 PM | 1:31:25 PM | Ok |
| 2 | $AqBIMuCh-multi-dispense-buffer-chamfered-cover-pad-blowout10 | 4 : Reservoir | 150 | A5 | 1:31:09 PM | 1:31:16 PM | Ok | 1 : Plate | 150 | B1 | 1:31:16 PM | 1:31:25 PM | Ok |

**24. Incubation - 2 min [1:31:30 PM - 1:33:30 PM]**

**25. 5. Apply Navenibody M1 and R2 [1:33:30 PM - 1:34:09 PM]**

| | | **Aspirate** | | | | | | **Dispense** | | | | | |
|---|---|---|---|---|---|---|---|---|---|---|---|---|---|
| Channel | Liquid Type | Labware | Volume | Well | Start (CET) | End (CET) | Status | Labware | Volume | Well | Start (CET) | End (CET) | Status |
| 2 | $Aqueous-buffer-chamfered-cover-pad-1 | 7 : Microtube Rack | 120 | C1 | 1:33:35 PM | 1:33:42 PM | Ok | 1 : Plate | 120 | A1 | 1:33:42 PM | 1:33:49 PM | Ok |
| 2 | $Aqueous-buffer-chamfered-cover-pad-1 | 7 : Microtube Rack | 120 | C1 | 1:33:49 PM | 1:33:56 PM | Ok | 1 : Plate | 120 | B1 | 1:33:56 PM | 1:34:03 PM | Ok |

**26. M1 and R2 incubation - 60 min [1:34:09 PM - 2:33:09 PM]**

**27. 5. Apply Navenibody M1 and R2 [2:33:09 PM - 2:33:45 PM]**

| | | **Aspirate** | | | | | | **Dispense** | | | | | |
|---|---|---|---|---|---|---|---|---|---|---|---|---|---|
| Channel | Liquid Type | Labware | Volume | Well | Start (CET) | End (CET) | Status | Labware | Volume | Well | Start (CET) | End (CET) | Status |
| 2 | $Aqueous-buffer-chamfered-cover-pad-1 | 7 : Microtube Rack | 120 | C1 | 2:33:14 PM | 2:33:20 PM | Ok | 1 : Plate | 120 | A1 | 2:33:20 PM | 2:33:27 PM | Ok |
| 2 | $Aqueous-buffer-chamfered-cover-pad-1 | 7 : Microtube Rack | 120 | C1 | 2:33:27 PM | 2:33:33 PM | Ok | 1 : Plate | 120 | B1 | 2:33:33 PM | 2:33:41 PM | Ok |

**28. M1 and R2 incubation - 60 min [2:33:46 PM - 3:32:45 PM]**

**29. Wash - 1X TBST [3:32:46 PM - 3:33:12 PM]**

| | | **Aspirate** | | | | | | **Dispense** | | | | | |
|---|---|---|---|---|---|---|---|---|---|---|---|---|---|
| Channel | Liquid Type | Labware | Volume | Well | Start (CET) | End (CET) | Status | Labware | Volume | Well | Start (CET) | End (CET) | Status |
| 1 | $AqBIMuCh-multi-dispense-buffer-chamfered-cover-pad-blowout10 | 4 : Reservoir | 150 | A5 | 3:32:51 PM | 3:32:59 PM | Ok | 1 : Plate | 150 | A1 | 3:32:59 PM | 3:33:07 PM | Ok |
| 2 | $AqBIMuCh-multi-dispense-buffer-chamfered-cover-pad-blowout10 | 4 : Reservoir | 150 | A5 | 3:32:51 PM | 3:32:59 PM | Ok | 1 : Plate | 150 | B1 | 3:32:59 PM | 3:33:07 PM | Ok |

**30. Wash - 1X TBST [3:33:12 PM - 3:33:39 PM]**

| | | **Aspirate** | | | | | | **Dispense** | | | | | |
|---|---|---|---|---|---|---|---|---|---|---|---|---|---|
| Channel | Liquid Type | Labware | Volume | Well | Start (CET) | End (CET) | Status | Labware | Volume | Well | Start (CET) | End (CET) | Status |
| 1 | $AqBIMuCh-multi-dispense-buffer-chamfered-cover-pad-blowout10 | 4 : Reservoir | 150 | A5 | 3:33:17 PM | 3:33:25 PM | Ok | 1 : Plate | 150 | A1 | 3:33:25 PM | 3:33:33 PM | Ok |
| 2 | $AqBIMuCh-multi-dispense-buffer-chamfered-cover-pad-blowout10 | 4 : Reservoir | 150 | A5 | 3:33:17 PM | 3:33:25 PM | Ok | 1 : Plate | 150 | B1 | 3:33:25 PM | 3:33:33 PM | Ok |

**31. Wash - 1X TBST [3:33:39 PM - 3:34:05 PM]**

| | | **Aspirate** | | | | | | **Dispense** | | | | | |
|---|---|---|---|---|---|---|---|---|---|---|---|---|---|
| Channel | Liquid Type | Labware | Volume | Well | Start (CET) | End (CET) | Status | Labware | Volume | Well | Start (CET) | End (CET) | Status |
| 1 | $AqBIMuCh-multi-dispense-buffer-chamfered-cover-pad-blowout10 | 4 : Reservoir | 150 | A5 | 3:33:44 PM | 3:33:52 PM | Ok | 1 : Plate | 150 | A1 | 3:33:52 PM | 3:34:00 PM | Ok |

| Channel | Liquid Type | Aspirate Labware | Volume | Well | Start (CET) | End (CET) | Status | Dispense Labware | Volume | Well | Start (CET) | End (CET) | Status |
|---|---|---|---|---|---|---|---|---|---|---|---|---|---|
| 2 | $AqBIMuCh-multi-dispense-buffer-chamfered-cover-pad-blowout10 | 4 : Reservoir | 150 | A5 | 3:33:44 PM | 3:33:52 PM | Ok | 1 : Plate | 150 | B1 | 3:33:52 PM | 3:34:00 PM | Ok |

**32. Incubate - 2 min [3:34:05 PM - 3:36:05 PM]**

**33. Wash - 1X TBST [3:36:05 PM - 3:36:31 PM]**

| | | **Aspirate** | | | | | | **Dispense** | | | | | |
|---|---|---|---|---|---|---|---|---|---|---|---|---|---|
| Channel | Liquid Type | Labware | Volume | Well | Start (CET) | End (CET) | Status | Labware | Volume | Well | Start (CET) | End (CET) | Status |
| 1 | $AqBIMuCh-multi-dispense-buffer-chamfered-cover-pad-blowout10 | 4 : Reservoir | 150 | A5 | 3:36:10 PM | 3:36:18 PM | Ok | 1 : Plate | 150 | A1 | 3:36:18 PM | 3:36:26 PM | Ok |
| 2 | $AqBIMuCh-multi-dispense-buffer-chamfered-cover-pad-blowout10 | 4 : Reservoir | 150 | A5 | 3:36:10 PM | 3:36:18 PM | Ok | 1 : Plate | 150 | B1 | 3:36:18 PM | 3:36:26 PM | Ok |

**34. Incubate - 2 min [3:36:31 PM - 3:38:31 PM]**

**35. Wash - 1X TBST [3:38:31 PM - 3:38:58 PM]**

| | | **Aspirate** | | | | | | **Dispense** | | | | | |
|---|---|---|---|---|---|---|---|---|---|---|---|---|---|
| Channel | Liquid Type | Labware | Volume | Well | Start (CET) | End (CET) | Status | Labware | Volume | Well | Start (CET) | End (CET) | Status |
| 1 | $AqBIMuCh-multi-dispense-buffer-chamfered-cover-pad-blowout10 | 4 : Reservoir | 150 | A5 | 3:38:36 PM | 3:38:44 PM | Ok | 1 : Plate | 150 | A1 | 3:38:44 PM | 3:38:53 PM | Ok |
| 2 | $AqBIMuCh-multi-dispense-buffer-chamfered-cover-pad-blowout10 | 4 : Reservoir | 150 | A5 | 3:38:36 PM | 3:38:44 PM | Ok | 1 : Plate | 150 | B1 | 3:38:44 PM | 3:38:53 PM | Ok |

**36. Incubate - 20 min [3:38:58 PM - 3:58:58 PM]**

**37. Wash - 1X TBST [3:58:58 PM - 3:59:24 PM]**

| | | **Aspirate** | | | | | | **Dispense** | | | | | |
|---|---|---|---|---|---|---|---|---|---|---|---|---|---|
| Channel | Liquid Type | Labware | Volume | Well | Start (CET) | End (CET) | Status | Labware | Volume | Well | Start (CET) | End (CET) | Status |
| 1 | $AqBIMuCh-multi-dispense-buffer-chamfered-cover-pad-blowout10 | 4 : Reservoir | 150 | A5 | 3:59:03 PM | 3:59:11 PM | Ok | 1 : Plate | 150 | A1 | 3:59:11 PM | 3:59:20 PM | Ok |
| 2 | $AqBIMuCh-multi-dispense-buffer-chamfered-cover-pad-blowout10 | 4 : Reservoir | 150 | A5 | 3:59:03 PM | 3:59:11 PM | Ok | 1 : Plate | 150 | B1 | 3:59:11 PM | 3:59:20 PM | Ok |

**38. Wash - 1X TBST [3:59:24 PM - 3:59:51 PM]**

| | | **Aspirate** | | | | | | **Dispense** | | | | | |
|---|---|---|---|---|---|---|---|---|---|---|---|---|---|
| Channel | Liquid Type | Labware | Volume | Well | Start (CET) | End (CET) | Status | Labware | Volume | Well | Start (CET) | End (CET) | Status |
| 1 | $AqBIMuCh-multi-dispense-buffer-chamfered-cover-pad-blowout10 | 4 : Reservoir | 150 | A5 | 3:59:29 PM | 3:59:37 PM | Ok | 1 : Plate | 150 | A1 | 3:59:37 PM | 3:59:46 PM | Ok |
| 2 | $AqBIMuCh-multi-dispense-buffer-chamfered-cover-pad-blowout10 | 4 : Reservoir | 150 | A5 | 3:59:29 PM | 3:59:37 PM | Ok | 1 : Plate | 150 | B1 | 3:59:37 PM | 3:59:46 PM | Ok |

**39. Incubate - 20 min [3:59:51 PM - 4:19:51 PM]**

**40. Wash - 1X TBST [4:19:51 PM - 4:20:17 PM]**

| | | **Aspirate** | | | | | | **Dispense** | | | | | |
|---|---|---|---|---|---|---|---|---|---|---|---|---|---|
| Channel | Liquid Type | Labware | Volume | Well | Start (CET) | End (CET) | Status | Labware | Volume | Well | Start (CET) | End (CET) | Status |
| 1 | $AqBIMuCh-multi-dispense-buffer-chamfered-cover-pad-blowout10 | 4 : Reservoir | 150 | A5 | 4:19:56 PM | 4:20:04 PM | Ok | 1 : Plate | 150 | A1 | 4:20:04 PM | 4:20:12 PM | Ok |
| 2 | $AqBIMuCh-multi-dispense-buffer-chamfered-cover-pad-blowout10 | 4 : Reservoir | 150 | A5 | 4:19:56 PM | 4:20:04 PM | Ok | 1 : Plate | 150 | B1 | 4:20:04 PM | 4:20:12 PM | Ok |

**41. Wash - 1X TBST [4:20:17 PM - 4:20:44 PM]**

| | | **Aspirate** | | | | | | **Dispense** | | | | | |
|---|---|---|---|---|---|---|---|---|---|---|---|---|---|
| Channel | Liquid Type | Labware | Volume | Well | Start (CET) | End (CET) | Status | Labware | Volume | Well | Start (CET) | End (CET) | Status |

| | | | | | | | | | | | | | |
|---|---|---|---|---|---|---|---|---|---|---|---|---|---|
| 1 | $AqBIMuCh-multi-dispense-buffer-chamfered-cover-pad-blowout10 | 4 : Reservoir | 150 | A5 | 4:20:23 PM | 4:20:30 PM | Ok | 1 : Plate | 150 | A1 | 4:20:30 PM | 4:20:39 PM | Ok |
| 2 | $AqBIMuCh-multi-dispense-buffer-chamfered-cover-pad-blowout10 | 4 : Reservoir | 150 | A5 | 4:20:23 PM | 4:20:30 PM | Ok | 1 : Plate | 150 | B1 | 4:20:30 PM | 4:20:39 PM | Ok |

**42. Wash - 1X TBST [4:20:44 PM - 4:21:10 PM]**

| | | Aspirate | | | | | | Dispense | | | | | |
|---|---|---|---|---|---|---|---|---|---|---|---|---|---|
| Channel | Liquid Type | Labware | Volume | Well | Start (CET) | End (CET) | Status | Labware | Volume | Well | Start (CET) | End (CET) | Status |
| 1 | $AqBIMuCh-multi-dispense-buffer-chamfered-cover-pad-blowout10 | 4 : Reservoir | 150 | A5 | 4:20:49 PM | 4:20:57 PM | Ok | 1 : Plate | 150 | A1 | 4:20:57 PM | 4:21:05 PM | Ok |
| 2 | $AqBIMuCh-multi-dispense-buffer-chamfered-cover-pad-blowout10 | 4 : Reservoir | 150 | A5 | 4:20:49 PM | 4:20:57 PM | Ok | 1 : Plate | 150 | B1 | 4:20:57 PM | 4:21:05 PM | Ok |

**43. 6. Apply Reaction 1 [4:21:10 PM - 4:21:41 PM]**

| | | Aspirate | | | | | | Dispense | | | | | |
|---|---|---|---|---|---|---|---|---|---|---|---|---|---|
| Channel | Liquid Type | Labware | Volume | Well | Start (CET) | End (CET) | Status | Labware | Volume | Well | Start (CET) | End (CET) | Status |
| 1 | $Aqueous-multi-dispense-buffer-chamfered-cover-pad-1 | 7 : Microtube Rack | 120 | D1 | 4:21:16 PM | 4:21:27 PM | Ok | 1 : Plate | 120 | A1 | 4:21:27 PM | 4:21:35 PM | Ok |
| 2 | $Aqueous-multi-dispense-buffer-chamfered-cover-pad-1 | 7 : Microtube Rack | 120 | D1 | 4:21:16 PM | 4:21:27 PM | Ok | 1 : Plate | 120 | B1 | 4:21:27 PM | 4:21:35 PM | Ok |

**44. Reaction 1 incubatiom - 30 min [4:21:41 PM - 4:51:41 PM]**

**45. Wash - 1X TBST [4:51:41 PM - 4:52:07 PM]**

| | | Aspirate | | | | | | Dispense | | | | | |
|---|---|---|---|---|---|---|---|---|---|---|---|---|---|
| Channel | Liquid Type | Labware | Volume | Well | Start (CET) | End (CET) | Status | Labware | Volume | Well | Start (CET) | End (CET) | Status |
| 1 | $AqBIMuCh-multi-dispense-buffer-chamfered-cover-pad-blowout10 | 4 : Reservoir | 150 | A5 | 4:51:46 PM | 4:51:54 PM | Ok | 1 : Plate | 150 | A1 | 4:51:54 PM | 4:52:02 PM | Ok |
| 2 | $AqBIMuCh-multi-dispense-buffer-chamfered-cover-pad-blowout10 | 4 : Reservoir | 150 | A5 | 4:51:46 PM | 4:51:54 PM | Ok | 1 : Plate | 150 | B1 | 4:51:54 PM | 4:52:02 PM | Ok |

**46. Incubation - 2 min [4:52:07 PM - 4:54:07 PM]**

**47. Wash - 1X TBST [4:54:07 PM - 4:54:34 PM]**

| | | Aspirate | | | | | | Dispense | | | | | |
|---|---|---|---|---|---|---|---|---|---|---|---|---|---|
| Channel | Liquid Type | Labware | Volume | Well | Start (CET) | End (CET) | Status | Labware | Volume | Well | Start (CET) | End (CET) | Status |
| 1 | $AqBIMuCh-multi-dispense-buffer-chamfered-cover-pad-blowout10 | 4 : Reservoir | 150 | A5 | 4:54:12 PM | 4:54:20 PM | Ok | 1 : Plate | 150 | A1 | 4:54:20 PM | 4:54:29 PM | Ok |
| 2 | $AqBIMuCh-multi-dispense-buffer-chamfered-cover-pad-blowout10 | 4 : Reservoir | 150 | A5 | 4:54:12 PM | 4:54:20 PM | Ok | 1 : Plate | 150 | B1 | 4:54:20 PM | 4:54:29 PM | Ok |

**48. Incubation - 2 min [4:54:34 PM - 4:56:34 PM]**

**49. Heat/Cool Step 36 C [4:56:34 PM - 4:56:44 PM]**

**50. 7. Apply Reaction 2 [4:56:44 PM - 4:57:16 PM]**

| | | Aspirate | | | | | | Dispense | | | | | |
|---|---|---|---|---|---|---|---|---|---|---|---|---|---|
| Channel | Liquid Type | Labware | Volume | Well | Start (CET) | End (CET) | Status | Labware | Volume | Well | Start (CET) | End (CET) | Status |
| 1 | $Aqueous-buffer-chamfered-cover-pad-1 | 7 : Microtube Rack | 120 | A3 | 4:56:49 PM | 4:57:02 PM | Ok | 1 : Plate | 120 | A1 | 4:57:02 PM | 4:57:10 PM | Ok |
| 2 | $Aqueous-buffer-chamfered-cover-pad-1 | 7 : Microtube Rack | 120 | A3 | 4:56:49 PM | 4:57:02 PM | Ok | 1 : Plate | 120 | B1 | 4:57:02 PM | 4:57:10 PM | Ok |

**51. R2 amplification - 45 min [4:57:16 PM - 5:42:16 PM]**

**52. 7. Apply Reaction 2 [5:42:16 PM - 5:42:46 PM]**

| | | Aspirate | | | | | | Dispense | | | | | |
|---|---|---|---|---|---|---|---|---|---|---|---|---|---|
| Channel | Liquid Type | Labware | Volume | Well | Start (CET) | End (CET) | Status | Labware | Volume | Well | Start (CET) | End (CET) | Status |

| | | | | | | | | | | | | | |
|---|---|---|---|---|---|---|---|---|---|---|---|---|---|
| 1 | $Aqueous-buffer-chamfered-cover-pad-1 | 7 : Microtube Rack | 120 | A3 | 5:42:21 PM | 5:42:33 PM | Ok | 1 : Plate | 120 | A1 | 5:42:33 PM | 5:42:41 PM | Ok |
| 2 | $Aqueous-buffer-chamfered-cover-pad-1 | 7 : Microtube Rack | 120 | A3 | 5:42:21 PM | 5:42:33 PM | Ok | 1 : Plate | 120 | B1 | 5:42:33 PM | 5:42:41 PM | Ok |

**53. R2 amplification - 45 min [5:42:46 PM - 6:27:46 PM]**

**54. Wash - 1X TBS [6:27:46 PM - 6:28:10 PM]**

| | | **Aspirate** | | | | | | **Dispense** | | | | | |
|---|---|---|---|---|---|---|---|---|---|---|---|---|---|
| Channel | Liquid Type | Labware | Volume | Well | Start (CET) | End (CET) | Status | Labware | Volume | Well | Start (CET) | End (CET) | Status |
| 1 | Aqueous | 4 : Reservoir | 150 | A4 | 6:27:51 PM | 6:28:00 PM | Ok | 1 : Plate | 150 | A1 | 6:28:00 PM | 6:28:05 PM | Ok |
| 2 | Aqueous | 4 : Reservoir | 150 | A4 | 6:27:51 PM | 6:28:00 PM | Ok | 1 : Plate | 150 | B1 | 6:28:00 PM | 6:28:05 PM | Ok |

**55. Incubation - 5 min [6:28:10 PM - 6:33:10 PM]**

**56. Wash - 1X TBS [6:33:10 PM - 6:33:34 PM]**

| | | **Aspirate** | | | | | | **Dispense** | | | | | |
|---|---|---|---|---|---|---|---|---|---|---|---|---|---|
| Channel | Liquid Type | Labware | Volume | Well | Start (CET) | End (CET) | Status | Labware | Volume | Well | Start (CET) | End (CET) | Status |
| 1 | Aqueous | 4 : Reservoir | 150 | A4 | 6:33:15 PM | 6:33:23 PM | Ok | 1 : Plate | 150 | A1 | 6:33:23 PM | 6:33:28 PM | Ok |
| 2 | Aqueous | 4 : Reservoir | 150 | A4 | 6:33:15 PM | 6:33:23 PM | Ok | 1 : Plate | 150 | B1 | 6:33:23 PM | 6:33:28 PM | Ok |

**57. Incubation - 5 min [6:33:34 PM - 6:38:34 PM]**

**58. Wash - 1X TBS 0.1 [6:38:34 PM - 6:39:00 PM]**

| | | **Aspirate** | | | | | | **Dispense** | | | | | |
|---|---|---|---|---|---|---|---|---|---|---|---|---|---|
| Channel | Liquid Type | Labware | Volume | Well | Start (CET) | End (CET) | Status | Labware | Volume | Well | Start (CET) | End (CET) | Status |
| 1 | $Aqueous-buffer-chamfered-cover-pad-1 | 4 : Reservoir | 150 | A3 | 6:38:39 PM | 6:38:47 PM | Ok | 1 : Plate | 150 | A1 | 6:38:47 PM | 6:38:55 PM | Ok |
| 2 | $Aqueous-buffer-chamfered-cover-pad-1 | 4 : Reservoir | 150 | A3 | 6:38:39 PM | 6:38:47 PM | Ok | 1 : Plate | 150 | B1 | 6:38:47 PM | 6:38:55 PM | Ok |

**59. Incubation - 5 min [6:39:00 PM - 6:44:00 PM]**

**60. Cool to RT [6:44:00 PM - 6:45:22 PM]**

**61. Wait - 5 min equilibration [6:45:22 PM - 6:50:22 PM]**

**62. 8. Apply HRP [6:50:23 PM - 6:50:52 PM]**

| | | **Aspirate** | | | | | | **Dispense** | | | | | |
|---|---|---|---|---|---|---|---|---|---|---|---|---|---|
| Channel | Liquid Type | Labware | Volume | Well | Start (CET) | End (CET) | Status | Labware | Volume | Well | Start (CET) | End (CET) | Status |
| 1 | $Aqueous-buffer-chamfered-cover-pad-1 | 7 : Microtube Rack | 120 | B3 | 6:50:28 PM | 6:50:39 PM | Ok | 1 : Plate | 120 | A1 | 6:50:39 PM | 6:50:47 PM | Ok |
| 2 | $Aqueous-buffer-chamfered-cover-pad-1 | 7 : Microtube Rack | 120 | B3 | 6:50:28 PM | 6:50:39 PM | Ok | 1 : Plate | 120 | B1 | 6:50:39 PM | 6:50:47 PM | Ok |

**63. HRP incubation - 30 min [6:50:52 PM - 7:20:52 PM]**

**64. Wash - 1X TBS [7:20:52 PM - 7:21:16 PM]**

| | | **Aspirate** | | | | | | **Dispense** | | | | | |
|---|---|---|---|---|---|---|---|---|---|---|---|---|---|
| Channel | Liquid Type | Labware | Volume | Well | Start (CET) | End (CET) | Status | Labware | Volume | Well | Start (CET) | End (CET) | Status |
| 1 | Aqueous | 4 : Reservoir | 150 | A4 | 7:20:57 PM | 7:21:05 PM | Ok | 1 : Plate | 150 | A1 | 7:21:05 PM | 7:21:11 PM | Ok |
| 2 | Aqueous | 4 : Reservoir | 150 | A4 | 7:20:57 PM | 7:21:05 PM | Ok | 1 : Plate | 150 | B1 | 7:21:05 PM | 7:21:11 PM | Ok |

**65. Incubation - 2 min [7:21:16 PM - 7:23:16 PM]**

**66. Wash - 1X TBS [7:23:16 PM - 7:23:40 PM]**

| | | **Aspirate** | | | | | | **Dispense** | | | | | |
|---|---|---|---|---|---|---|---|---|---|---|---|---|---|
| Channel | Liquid Type | Labware | Volume | Well | Start (CET) | End (CET) | Status | Labware | Volume | Well | Start (CET) | End (CET) | Status |
| 1 | Aqueous | 4 : Reservoir | 150 | A4 | 7:23:21 PM | 7:23:30 PM | Ok | 1 : Plate | 150 | A1 | 7:23:30 PM | 7:23:35 PM | Ok |
| 2 | Aqueous | 4 : Reservoir | 150 | A4 | 7:23:21 PM | 7:23:30 PM | Ok | 1 : Plate | 150 | B1 | 7:23:30 PM | 7:23:35 PM | Ok |

**67. Incubation - 2 min [7:23:40 PM - 7:25:40 PM]**

**68. Wash - PBS [7:25:40 PM - 7:26:07 PM]**

| | | **Aspirate** | | | | | | **Dispense** | | | | | |
|---|---|---|---|---|---|---|---|---|---|---|---|---|---|
| Channel | Liquid Type | Labware | Volume | Well | Start (CET) | End (CET) | Status | Labware | Volume | Well | Start (CET) | End (CET) | Status |
| 1 | $Aqueous-buffer-chamfered-cover-pad-1 | 4 : Reservoir | 150 | A8 | 7:25:45 PM | 7:25:53 PM | Ok | 1 : Plate | 150 | A1 | 7:25:53 PM | 7:26:02 PM | Ok |
| 2 | $Aqueous-buffer-chamfered-cover-pad-1 | 4 : Reservoir | 150 | A8 | 7:25:45 PM | 7:25:53 PM | Ok | 1 : Plate | 150 | B1 | 7:25:53 PM | 7:26:02 PM | Ok |

**69. 9. Apply Detection aluora [7:26:07 PM - 7:26:38 PM]**

| | | **Aspirate** | | | | | | **Dispense** | | | | | |
|---|---|---|---|---|---|---|---|---|---|---|---|---|---|
| Channel | Liquid Type | Labware | Volume | Well | Start (CET) | End (CET) | Status | Labware | Volume | Well | Start (CET) | End (CET) | Status |
| 1 | $Aqueous-buffer-chamfered-cover-pad-1 | 7 : Microtube Rack | 120 | C3 | 7:26:12 PM | 7:26:24 PM | Ok | 1 : Plate | 120 | A1 | 7:26:24 PM | 7:26:32 PM | Ok |

| Channel | Liquid Type | Aspirate Labware | Volume | Well | Start (CET) | End (CET) | Status | Dispense Labware | Volume | Well | Start (CET) | End (CET) | Status |
|---|---|---|---|---|---|---|---|---|---|---|---|---|---|
| 2 | $Aqueous-buffer-chamfered-cover-pad-1 | 7 : Microtube Rack | 120 | C3 | 7:26:12 PM | 7:26:24 PM | Ok | 1 : Plate | 120 | B1 | 7:26:24 PM | 7:26:32 PM | Ok |

**70. Aluora incubation - 10 min [7:26:38 PM - 7:36:38 PM]**

**71. Wash - PBS [7:36:38 PM - 7:37:05 PM]**

| | | **Aspirate** | | | | | | **Dispense** | | | | | |
|---|---|---|---|---|---|---|---|---|---|---|---|---|---|
| Channel | Liquid Type | Labware | Volume | Well | Start (CET) | End (CET) | Status | Labware | Volume | Well | Start (CET) | End (CET) | Status |
| 1 | $Aqueous-buffer-chamfered-cover-pad-1 | 4 : Reservoir | 150 | A8 | 7:36:43 PM | 7:36:51 PM | Ok | 1 : Plate | 150 | A1 | 7:36:51 PM | 7:37:00 PM | Ok |
| 2 | $Aqueous-buffer-chamfered-cover-pad-1 | 4 : Reservoir | 150 | A8 | 7:36:43 PM | 7:36:51 PM | Ok | 1 : Plate | 150 | B1 | 7:36:51 PM | 7:37:00 PM | Ok |

**72. Incubation - 2 min [7:37:05 PM - 7:39:05 PM]**

**73. Wash - PBS [7:39:05 PM - 7:39:32 PM]**

| | | **Aspirate** | | | | | | **Dispense** | | | | | |
|---|---|---|---|---|---|---|---|---|---|---|---|---|---|
| Channel | Liquid Type | Labware | Volume | Well | Start (CET) | End (CET) | Status | Labware | Volume | Well | Start (CET) | End (CET) | Status |
| 1 | $Aqueous-buffer-chamfered-cover-pad-1 | 4 : Reservoir | 150 | A8 | 7:39:10 PM | 7:39:18 PM | Ok | 1 : Plate | 150 | A1 | 7:39:18 PM | 7:39:27 PM | Ok |
| 2 | $Aqueous-buffer-chamfered-cover-pad-1 | 4 : Reservoir | 150 | A8 | 7:39:10 PM | 7:39:18 PM | Ok | 1 : Plate | 150 | B1 | 7:39:18 PM | 7:39:27 PM | Ok |

**74. Incubation - 2 min [7:39:32 PM - 7:41:32 PM]**

**75. 9. Apply DAPI [7:41:32 PM - 7:42:06 PM]**

| | | **Aspirate** | | | | | | **Dispense** | | | | | |
|---|---|---|---|---|---|---|---|---|---|---|---|---|---|
| Channel | Liquid Type | Labware | Volume | Well | Start (CET) | End (CET) | Status | Labware | Volume | Well | Start (CET) | End (CET) | Status |
| 1 | $Aqueous-multi-dispense-buffer-chamfered-cover-pad-1 | 7 : Microtube Rack | 120 | D3 | 7:41:37 PM | 7:41:53 PM | Ok | 1 : Plate | 120 | A1 | 7:41:53 PM | 7:42:01 PM | Ok |
| 2 | $Aqueous-multi-dispense-buffer-chamfered-cover-pad-1 | 7 : Microtube Rack | 120 | D3 | 7:41:37 PM | 7:41:53 PM | Ok | 1 : Plate | 120 | B1 | 7:41:53 PM | 7:42:01 PM | Ok |

**76. DAPI incubation- 4 min [7:42:06 PM - 7:46:06 PM]**

**77. 9. Apply DAPI [7:46:06 PM - 7:46:40 PM]**

| | | **Aspirate** | | | | | | **Dispense** | | | | | |
|---|---|---|---|---|---|---|---|---|---|---|---|---|---|
| Channel | Liquid Type | Labware | Volume | Well | Start (CET) | End (CET) | Status | Labware | Volume | Well | Start (CET) | End (CET) | Status |
| 1 | $Aqueous-multi-dispense-buffer-chamfered-cover-pad-1 | 7 : Microtube Rack | 120 | D3 | 7:46:11 PM | 7:46:27 PM | Ok | 1 : Plate | 120 | A1 | 7:46:27 PM | 7:46:36 PM | Ok |
| 2 | $Aqueous-multi-dispense-buffer-chamfered-cover-pad-1 | 7 : Microtube Rack | 120 | D3 | 7:46:11 PM | 7:46:27 PM | Ok | 1 : Plate | 120 | B1 | 7:46:27 PM | 7:46:36 PM | Ok |

**78. DAPI incubation- 4 min [7:46:40 PM - 7:50:40 PM]**

**79. Wash - PBS [7:50:40 PM - 7:51:07 PM]**

| | | **Aspirate** | | | | | | **Dispense** | | | | | |
|---|---|---|---|---|---|---|---|---|---|---|---|---|---|
| Channel | Liquid Type | Labware | Volume | Well | Start (CET) | End (CET) | Status | Labware | Volume | Well | Start (CET) | End (CET) | Status |
| 1 | $Aqueous-buffer-chamfered-cover-pad-1 | 4 : Reservoir | 150 | A8 | 7:50:46 PM | 7:50:53 PM | Ok | 1 : Plate | 150 | A1 | 7:50:53 PM | 7:51:02 PM | Ok |
| 2 | $Aqueous-buffer-chamfered-cover-pad-1 | 4 : Reservoir | 150 | A8 | 7:50:46 PM | 7:50:53 PM | Ok | 1 : Plate | 150 | B1 | 7:50:53 PM | 7:51:02 PM | Ok |

**80. Wait - 2 min [7:51:07 PM - 7:53:07 PM]**

**81. Wash - PBS [7:53:08 PM - 7:53:35 PM]**

| | | **Aspirate** | | | | | | **Dispense** | | | | | |
|---|---|---|---|---|---|---|---|---|---|---|---|---|---|
| Channel | Liquid Type | Labware | Volume | Well | Start (CET) | End (CET) | Status | Labware | Volume | Well | Start (CET) | End (CET) | Status |
| 1 | $Aqueous-buffer-chamfered-cover-pad-1 | 4 : Reservoir | 150 | A8 | 7:53:12 PM | 7:53:21 PM | Ok | 1 : Plate | 150 | A1 | 7:53:21 PM | 7:53:30 PM | Ok |
| 2 | $Aqueous-buffer-chamfered-cover-pad-1 | 4 : Reservoir | 150 | A8 | 7:53:12 PM | 7:53:21 PM | Ok | 1 : Plate | 150 | B1 | 7:53:21 PM | 7:53:30 PM | Ok |

**82. 1X TBST [7:53:35 PM - 7:54:02 PM]**

| | | **Aspirate** | | | | | | **Dispense** | | | | | |
|---|---|---|---|---|---|---|---|---|---|---|---|---|---|
| Channel | Liquid Type | Labware | Volume | Well | Start (CET) | End (CET) | Status | Labware | Volume | Well | Start (CET) | End (CET) | Status |
| 1 | $AqBlMuCh-multi-dispense-buffer-chamfered-cover-pad-blowout10 | 4 : Reservoir | 150 | A5 | 7:53:40 PM | 7:53:48 PM | Ok | 1 : Plate | 150 | A1 | 7:53:48 PM | 7:53:57 PM | Ok |

| 2 | $AqBlMuCh-multi-dispense-buffer-chamfered-cover-pad-blowout10 | 4 : Reservoir | 150 | A5 | 7:53:40 PM | 7:53:48 PM | Ok | 1 : Plate | 150 | B1 | 7:53:48 PM | 7:53:57 PM | Ok |
|---|---|---|---|---|---|---|---|---|---|---|---|---|---|

**83. 1X TBST [7:54:02 PM - 7:54:29 PM]**

| | | Aspirate | | | | | | Dispense | | | | | |
|---|---|---|---|---|---|---|---|---|---|---|---|---|---|
| Channel | Liquid Type | Labware | Volume | Well | Start (CET) | End (CET) | Status | Labware | Volume | Well | Start (CET) | End (CET) | Status |
| 1 | $AqBlMuCh-multi-dispense-buffer-chamfered-cover-pad-blowout10 | 4 : Reservoir | 150 | A5 | 7:54:07 PM | 7:54:15 PM | Ok | 1 : Plate | 150 | A1 | 7:54:15 PM | 7:54:24 PM | Ok |
| 2 | $AqBlMuCh-multi-dispense-buffer-chamfered-cover-pad-blowout10 | 4 : Reservoir | 150 | A5 | 7:54:07 PM | 7:54:15 PM | Ok | 1 : Plate | 150 | B1 | 7:54:15 PM | 7:54:24 PM | Ok |

**84. Wait - 2 min [7:54:29 PM - 7:56:29 PM]**

**85. 0.1X TBST [7:56:29 PM - 7:56:56 PM]**

| | | Aspirate | | | | | | Dispense | | | | | |
|---|---|---|---|---|---|---|---|---|---|---|---|---|---|
| Channel | Liquid Type | Labware | Volume | Well | Start (CET) | End (CET) | Status | Labware | Volume | Well | Start (CET) | End (CET) | Status |
| 1 | $Aqueous-multi-dispense-buffer-chamfered-cover-pad-1 | 4 : Reservoir | 150 | A7 | 7:56:34 PM | 7:56:42 PM | Ok | 1 : Plate | 150 | A1 | 7:56:42 PM | 7:56:51 PM | Ok |
| 2 | $Aqueous-multi-dispense-buffer-chamfered-cover-pad-1 | 4 : Reservoir | 150 | A7 | 7:56:34 PM | 7:56:42 PM | Ok | 1 : Plate | 150 | B1 | 7:56:42 PM | 7:56:51 PM | Ok |

**86. 0.1X TBST [7:56:56 PM - 7:57:23 PM]**

| | | Aspirate | | | | | | Dispense | | | | | |
|---|---|---|---|---|---|---|---|---|---|---|---|---|---|
| Channel | Liquid Type | Labware | Volume | Well | Start (CET) | End (CET) | Status | Labware | Volume | Well | Start (CET) | End (CET) | Status |
| 1 | $Aqueous-multi-dispense-buffer-chamfered-cover-pad-1 | 4 : Reservoir | 150 | A7 | 7:57:01 PM | 7:57:09 PM | Ok | 1 : Plate | 150 | A1 | 7:57:09 PM | 7:57:18 PM | Ok |
| 2 | $Aqueous-multi-dispense-buffer-chamfered-cover-pad-1 | 4 : Reservoir | 150 | A7 | 7:57:01 PM | 7:57:09 PM | Ok | 1 : Plate | 150 | B1 | 7:57:09 PM | 7:57:18 PM | Ok |

**87. Wash - PBS [7:57:23 PM - 7:57:50 PM]**

| | | Aspirate | | | | | | Dispense | | | | | |
|---|---|---|---|---|---|---|---|---|---|---|---|---|---|
| Channel | Liquid Type | Labware | Volume | Well | Start (CET) | End (CET) | Status | Labware | Volume | Well | Start (CET) | End (CET) | Status |
| 1 | $Aqueous-buffer-chamfered-cover-pad-1 | 4 : Reservoir | 150 | A8 | 7:57:28 PM | 7:57:36 PM | Ok | 1 : Plate | 150 | A1 | 7:57:36 PM | 7:57:45 PM | Ok |
| 2 | $Aqueous-buffer-chamfered-cover-pad-1 | 4 : Reservoir | 150 | A8 | 7:57:28 PM | 7:57:36 PM | Ok | 1 : Plate | 150 | B1 | 7:57:36 PM | 7:57:45 PM | Ok |

**88. Cool to 4C [7:57:50 PM - 8:06:39 PM]**

**89. Wait until user resumes [8:06:39 PM - 9:08:00 AM]**

### Pauses

Pause 1: Wait Step Initiated Pause

- Length of pause: 1800 seconds
- Start of pause: Thursday, February 26, 2026 10:42:51 AM
- End of pause: Thursday, February 26, 2026 11:12:51 AM

Pause 2: Wait Step Initiated Pause

- Length of pause: 60 seconds
- Start of pause: Thursday, February 26, 2026 11:13:17 AM
- End of pause: Thursday, February 26, 2026 11:14:17 AM

Pause 3: Wait Step Initiated Pause

- Length of pause: 60 seconds
- Start of pause: Thursday, February 26, 2026 11:14:44 AM
- End of pause: Thursday, February 26, 2026 11:15:44 AM

Pause 4: Wait Step Initiated Pause

- Length of pause: 60 seconds
- Start of pause: Thursday, February 26, 2026 11:16:11 AM
- End of pause: Thursday, February 26, 2026 11:17:11 AM

Pause 5: Wait Step Initiated Pause

- Length of pause: 3600 seconds
- Start of pause: Thursday, February 26, 2026 11:18:21 AM
- End of pause: Thursday, February 26, 2026 12:18:21 PM

Pause 6: Wait Step Initiated Pause

- Length of pause: 180 seconds
- Start of pause: Thursday, February 26, 2026 12:18:47 PM
- End of pause: Thursday, February 26, 2026 12:21:47 PM

Pause 7: Wait Step Initiated Pause

- Length of pause: 180 seconds
- Start of pause: Thursday, February 26, 2026 12:22:15 PM
- End of pause: Thursday, February 26, 2026 12:25:15 PM

Pause 8: Wait Step Initiated Pause

- Length of pause: 3600 seconds
- Start of pause: Thursday, February 26, 2026 12:26:09 PM
- End of pause: Thursday, February 26, 2026 1:26:09 PM

Pause 9: Wait Step Initiated Pause

- Length of pause: 120 seconds
- Start of pause: Thursday, February 26, 2026 1:26:37 PM
- End of pause: Thursday, February 26, 2026 1:28:37 PM

Pause 10: Wait Step Initiated Pause

- Length of pause: 120 seconds
- Start of pause: Thursday, February 26, 2026 1:29:03 PM
- End of pause: Thursday, February 26, 2026 1:31:03 PM

Pause 11: Wait Step Initiated Pause

- Length of pause: 120 seconds
- Start of pause: Thursday, February 26, 2026 1:31:30 PM
- End of pause: Thursday, February 26, 2026 1:33:30 PM

Pause 12: Wait Step Initiated Pause

- Length of pause: 3540 seconds
- Start of pause: Thursday, February 26, 2026 1:34:09 PM
- End of pause: Thursday, February 26, 2026 2:33:09 PM

Pause 13: Wait Step Initiated Pause

- Length of pause: 3540 seconds
- Start of pause: Thursday, February 26, 2026 2:33:46 PM
- End of pause: Thursday, February 26, 2026 3:32:46 PM

Pause 14: Wait Step Initiated Pause

- Length of pause: 120 seconds
- Start of pause: Thursday, February 26, 2026 3:34:05 PM
- End of pause: Thursday, February 26, 2026 3:36:05 PM

Pause 15: Wait Step Initiated Pause

- Length of pause: 120 seconds
- Start of pause: Thursday, February 26, 2026 3:36:31 PM
- End of pause: Thursday, February 26, 2026 3:38:31 PM

Pause 16: Wait Step Initiated Pause

- Length of pause: 1200 seconds
- Start of pause: Thursday, February 26, 2026 3:38:58 PM
- End of pause: Thursday, February 26, 2026 3:58:58 PM

Pause 17: Wait Step Initiated Pause

- Length of pause: 1200 seconds
- Start of pause: Thursday, February 26, 2026 3:59:51 PM
- End of pause: Thursday, February 26, 2026 4:19:51 PM

Pause 18: Wait Step Initiated Pause

- Length of pause: 1800 seconds
- Start of pause: Thursday, February 26, 2026 4:21:41 PM
- End of pause: Thursday, February 26, 2026 4:51:41 PM

Pause 19: Wait Step Initiated Pause

- Length of pause: 120 seconds
- Start of pause: Thursday, February 26, 2026 4:52:07 PM
- End of pause: Thursday, February 26, 2026 4:54:07 PM

Pause 20: Wait Step Initiated Pause

- Length of pause: 120 seconds
- Start of pause: Thursday, February 26, 2026 4:54:34 PM
- End of pause: Thursday, February 26, 2026 4:56:34 PM

Pause 21: Wait Step Initiated Pause

- Length of pause: 2700 seconds
- Start of pause: Thursday, February 26, 2026 4:57:16 PM
- End of pause: Thursday, February 26, 2026 5:42:16 PM

Pause 22: Wait Step Initiated Pause

- Length of pause: 2700 seconds
- Start of pause: Thursday, February 26, 2026 5:42:46 PM
- End of pause: Thursday, February 26, 2026 6:27:46 PM

Pause 23: Wait Step Initiated Pause

- Length of pause: 300 seconds
- Start of pause: Thursday, February 26, 2026 6:28:10 PM
- End of pause: Thursday, February 26, 2026 6:33:10 PM

Pause 24: Wait Step Initiated Pause

- Length of pause: 300 seconds
- Start of pause: Thursday, February 26, 2026 6:33:34 PM
- End of pause: Thursday, February 26, 2026 6:38:34 PM

Pause 25: Wait Step Initiated Pause

- Length of pause: 300 seconds
- Start of pause: Thursday, February 26, 2026 6:39:00 PM
- End of pause: Thursday, February 26, 2026 6:44:00 PM

Pause 26: Wait Step Initiated Pause

- Length of pause: 300 seconds
- Start of pause: Thursday, February 26, 2026 6:45:22 PM
- End of pause: Thursday, February 26, 2026 6:50:23 PM

Pause 27: Wait Step Initiated Pause

- Length of pause: 1800 seconds
- Start of pause: Thursday, February 26, 2026 6:50:52 PM
- End of pause: Thursday, February 26, 2026 7:20:52 PM

Pause 28: Wait Step Initiated Pause

- Length of pause: 120 seconds
- Start of pause: Thursday, February 26, 2026 7:21:16 PM
- End of pause: Thursday, February 26, 2026 7:23:16 PM

Pause 29: Wait Step Initiated Pause

- Length of pause: 120 seconds
- Start of pause: Thursday, February 26, 2026 7:23:40 PM
- End of pause: Thursday, February 26, 2026 7:25:40 PM

Pause 30: Wait Step Initiated Pause

- Length of pause: 600 seconds
- Start of pause: Thursday, February 26, 2026 7:26:38 PM
- End of pause: Thursday, February 26, 2026 7:36:38 PM

Pause 31: Wait Step Initiated Pause

- Length of pause: 120 seconds
- Start of pause: Thursday, February 26, 2026 7:37:05 PM
- End of pause: Thursday, February 26, 2026 7:39:05 PM

Pause 32: Wait Step Initiated Pause

- Length of pause: 120 seconds
- Start of pause: Thursday, February 26, 2026 7:39:32 PM
- End of pause: Thursday, February 26, 2026 7:41:32 PM

Pause 33: Wait Step Initiated Pause

- Length of pause: 240 seconds
- Start of pause: Thursday, February 26, 2026 7:42:06 PM
- End of pause: Thursday, February 26, 2026 7:46:06 PM

Pause 34: Wait Step Initiated Pause

- Length of pause: 240 seconds
- Start of pause: Thursday, February 26, 2026 7:46:40 PM
- End of pause: Thursday, February 26, 2026 7:50:40 PM

Pause 35: Wait Step Initiated Pause

- Length of pause: 120 seconds

- Start of pause: Thursday, February 26, 2026 7:51:07 PM
- End of pause: Thursday, February 26, 2026 7:53:07 PM

Pause 36: Wait Step Initiated Pause

- Length of pause: 120 seconds
- Start of pause: Thursday, February 26, 2026 7:54:29 PM
- End of pause: Thursday, February 26, 2026 7:56:29 PM

Pause 37: Step Initiated Pause

- Length of pause: 46879 seconds
- Start of pause: Thursday, February 26, 2026 8:06:40 PM
- End of pause: Friday, February 27, 2026 9:07:59 AM

### Door Events

- Door is open @ Thursday, February 26, 2026 10:42:53 AM
- Door is closed @ Thursday, February 26, 2026 10:43:00 AM

### Error Messages

[none]

### Deck Snapshot

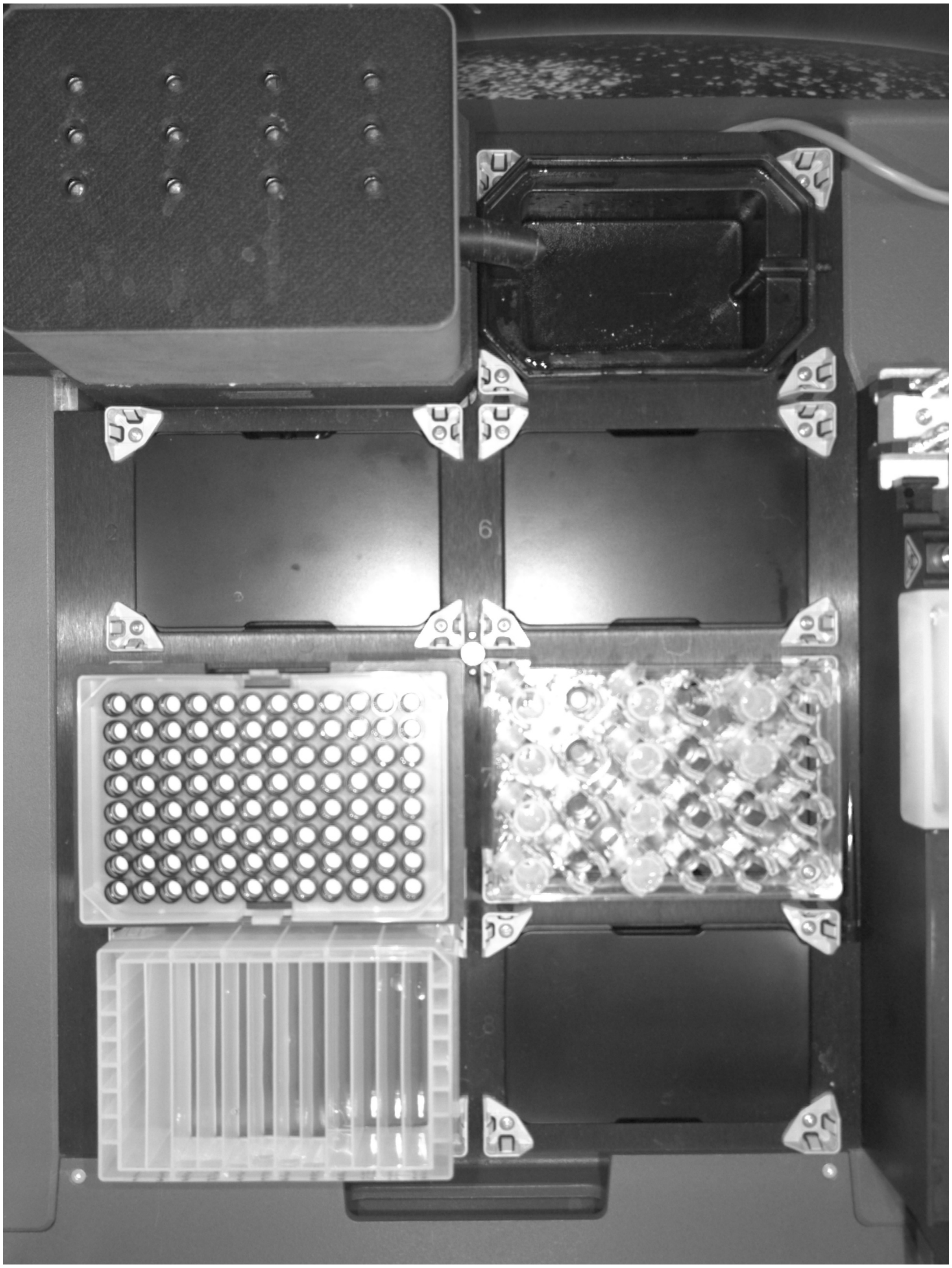

Signature: ________________________________

Operator: spu
