## Supplemental Figure 4 for "A workflow for combined detection of protein interactions and cell types for translational studies"

### Unmixing Quality Metrics Report
### EVOS™ S1000 Spatial Imaging System

**September 23, 2025 (Rev. 1.0)**

Thermo Fisher SCIENTIFIC

#### Sample information

| | |
|---|---|
| Protocol | EV-project-V3-230925 |
| Resolution | 0.324 µm/pixel |
| Objective | 20x |
| Number of fluorophores | 7 |
| Number of channels | 15 |

#### Spectra extracted from single-color control (SCC) samples

In Figure 1, the spectra extracted from the SCC samples are shown. Each extracted spectrum should have a unique signature. If unexpected signatures are seen, refer to the “Masks and extraction channel images” section. If a spectrum is missing, it indicates that data extraction has not been successful and therefore SCC samples will need to be reimaged.

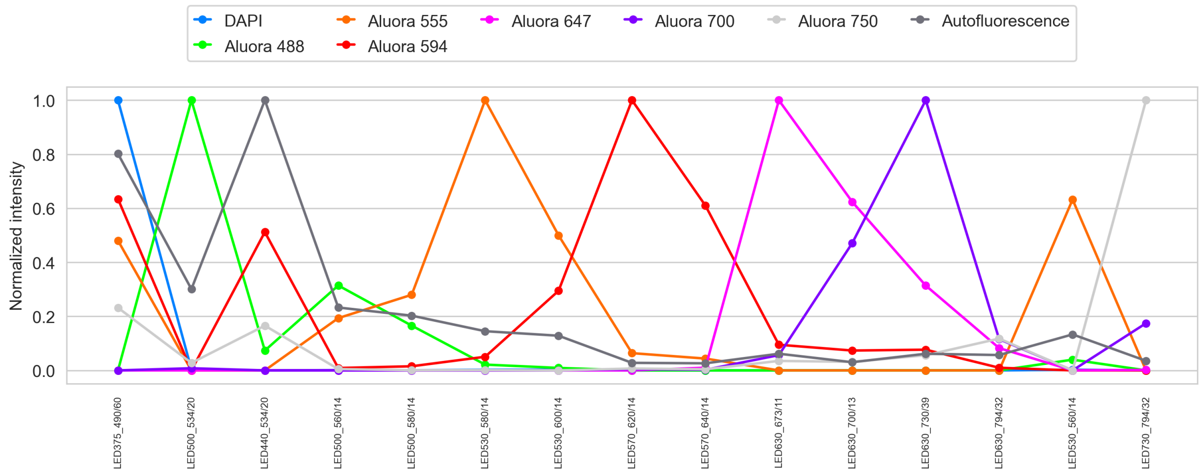


**Figure 1.** Spectra extracted from single-color controls samples

In Figure 2, the spectra extracted in every primary and support channel for every SCC sample are visualized as heatmaps, Relative intensity is shown in a grey shade-scale. The signal is represented from low (black) to high (white). Of note, if an acquired image has more than 1% overexposed pixels, it is highlighted in red in Figure 2, and it indicates that the SCC will need to be re-imaged. High-quality spectral extraction necessitates that appropriate exposure settings are used during image acquisition, in order to minimize overexposed pixels. Refer to the “Troubleshooting tips” section for more information.



**For Research Use Only. Not for use in diagnostic procedures** © 2024 Thermo Fisher Scientific Inc. All rights reserved. All Unmixing Metrics Report trademarks are the property of Thermo Fisher Scientific and its subsidiaries unless otherwise specified.

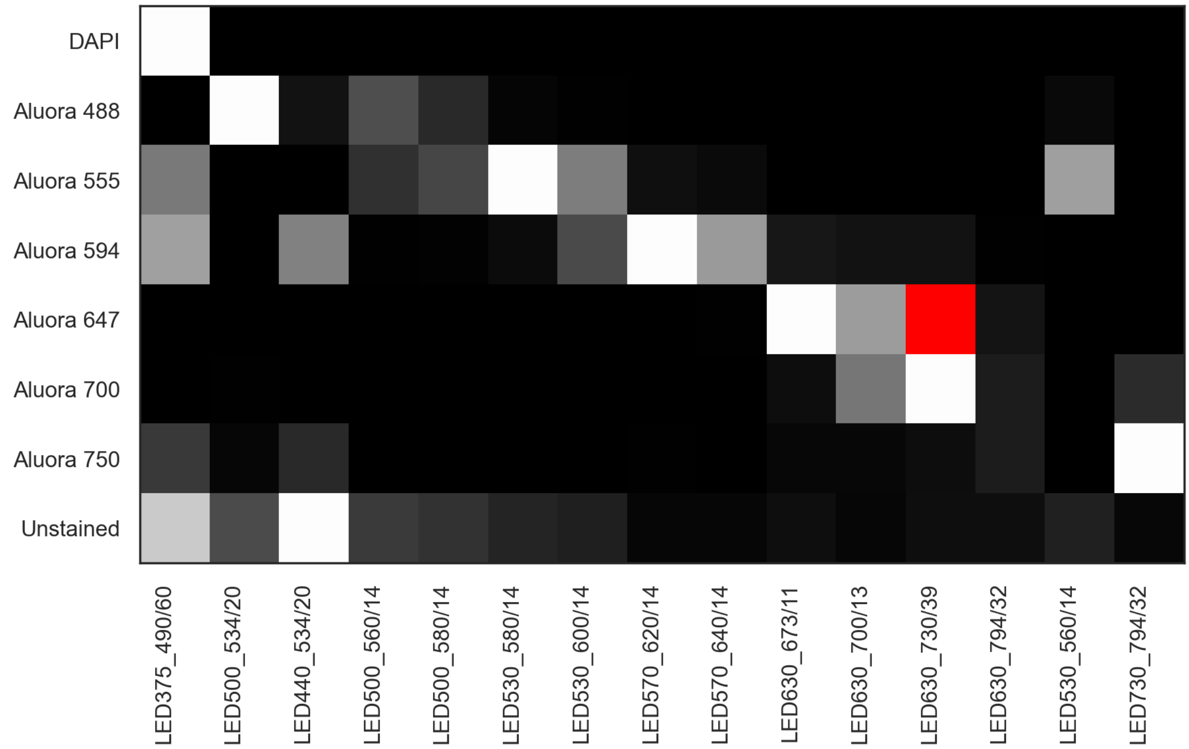


**Figure 2.** Heatmap visualization of the spectra extracted in every primary and support channel (shown in the columns) for every SCC sample (shown in the rows).

#### Unmixing matrix assessment on SCC samples

In Figure 3, the raw images (before unmixing) acquired for each primary channel are displayed. Columns represent the SCC samples whereas rows correspond to the primary channels that were in the protocol, listed in the Sample information in page 1. Images on the diagonal line correspond to the raw images acquired for the targets in their primary channels. Spectral overlap is observed in raw images when a fluorophore is detected in off-target channels, outside the diagonal line.


**For Research Use Only. Not for use in diagnostic procedures** © 2024 Thermo Fisher Scientific Inc. All rights reserved. All Unmixing Metrics Report trademarks are the property of Thermo Fisher Scientific and its subsidiaries unless otherwise specified.

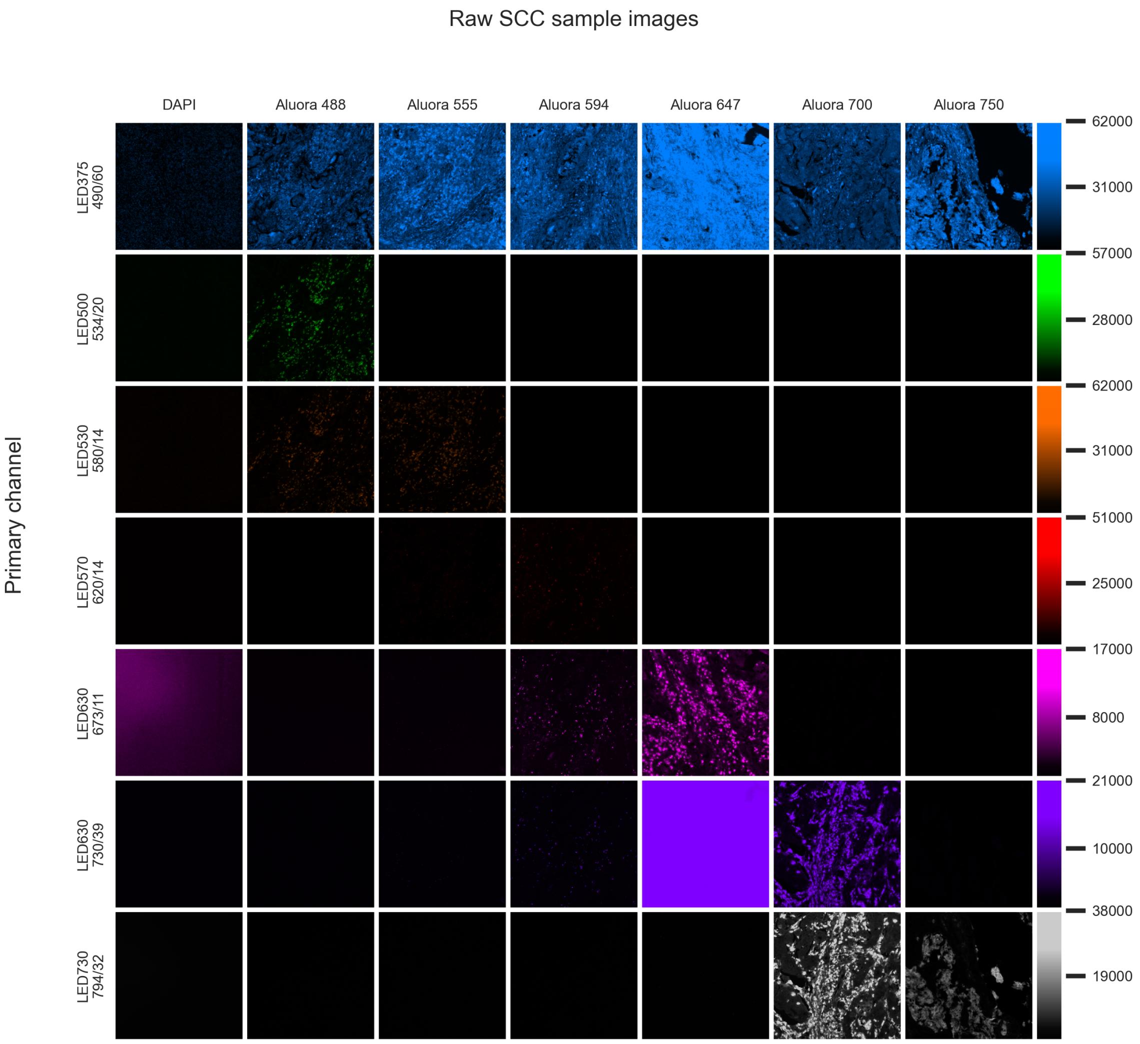


**Figure 3.** Visualization of the raw images acquired for SCC Samples (in each column) in each of the primary channels (in every row).

In Figure 4, the images obtained after applying the unmixing matrix are shown. Columns represent the SCC samples whereas rows correspond to the unmixed channels in the protocol. In this image, the unmixing matrix has been applied to each SCC, thus separating signals into their own target channels and eliminating signal bleedthrough.

A high-quality unmixing matrix is expected to result in:

- a high signal output in all primary channels along the diagonal line
- a low signal in off-target channels, outside the diagonal line


**For Research Use Only. Not for use in diagnostic procedures** © 2024 Thermo Fisher Scientific Inc. All rights reserved. All Unmixing Metrics Report trademarks are the property of Thermo Fisher Scientific and its subsidiaries unless otherwise specified.

Unmixed SCC sample images

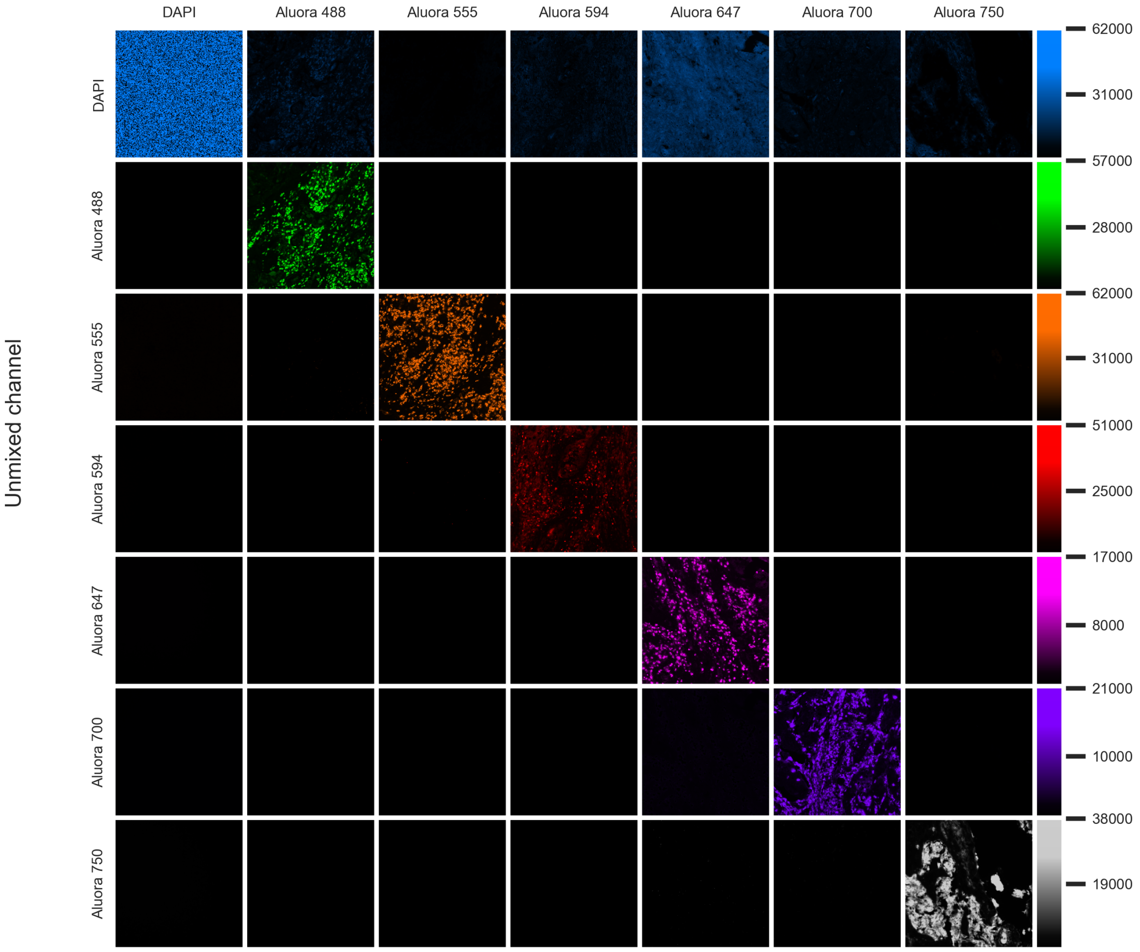


**Figure 4.** Visualization of the spectrally unmixed images acquired for SCC Samples (in each column) in each of the primary channels (in every row).

#### Quantitative unmixing metrics

In Figure 5, a quantitative assessment of the unmixing quality is presented, via the calculation of the Fraction Bleedthrough (FB) remaining in off-target channels, with an additional visualization using heatmaps. This parameter is defined as the ratio of the signal observed in off-target channels to the signal of the target fluorophore (obtained in primary channels) after unmixing has been applied. It is used for quantifying the fluorescence seen in the unmixed images shown in Figure 4.

A high-quality unmixing matrix is characterized by FB values that are equal or less than 0.25. If the values are found to be above this threshold, it is recommended that the pair of fluorophores in question is first identified. Then, their spectral intensities and the images used to extract the spectra should be further examined in the “Masks and extraction channel images” section. Also, please refer to the “Troubleshooting tips” section for more information.



**For Research Use Only. Not for use in diagnostic procedures** © 2024 Thermo Fisher Scientific Inc. All rights reserved. All Unmixing Metrics Report trademarks are the property of Thermo Fisher Scientific and its subsidiaries unless otherwise specified.

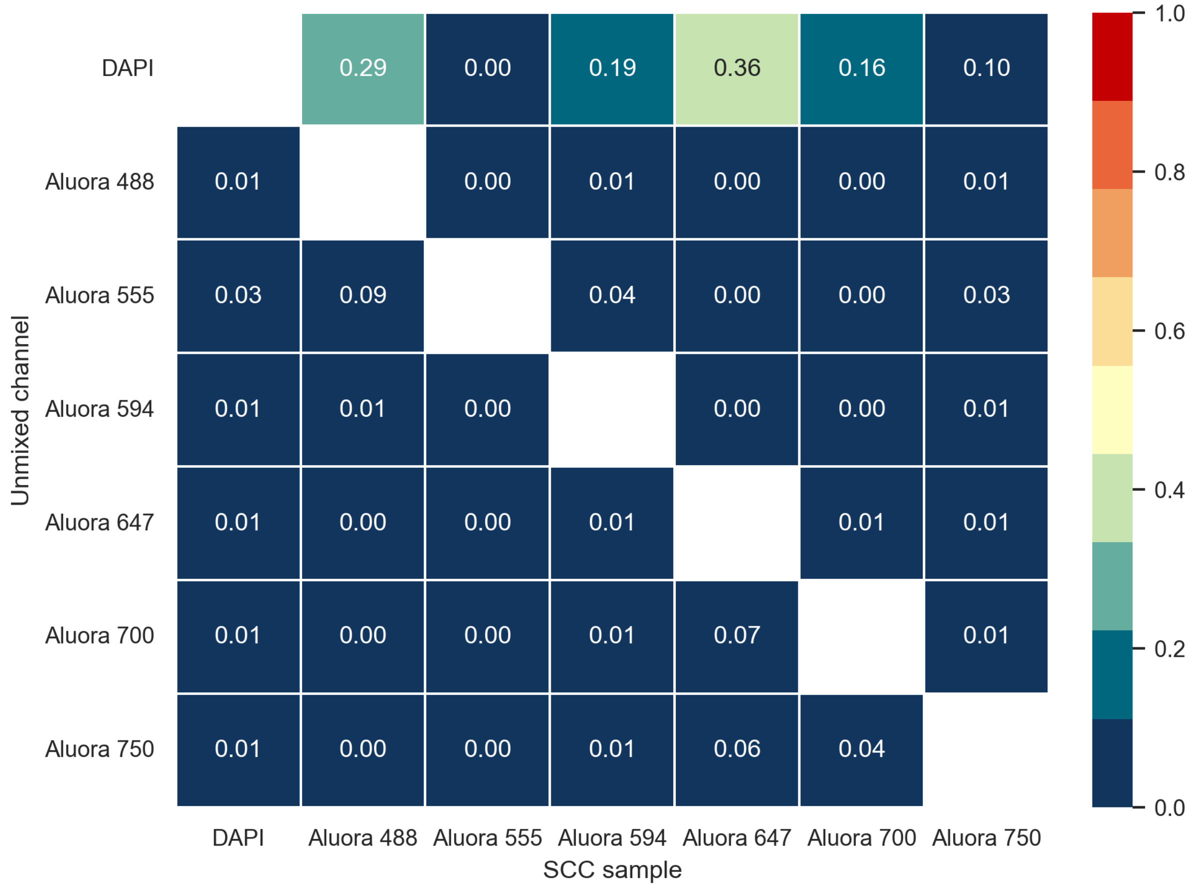


**Figure 5.** Quantitative assessment of the unmixing quality via calculation of the fraction bleedthrough values.

In Figure 6, a second metric for quantitative assessment of the unmixing quality is shown, in this case via the calculation of the Effective Dynamic Range (EDR). This parameter is defined as the fluorescent intensity from every target in its dedicated primary channel after subtracting the bleedthrough from neighboring fluorophores.

A high EDR ensures that a target can be unmixed from the multiplex sample with high confidence. Specifically, a value of EDR of at least 1000 can be regarded as an indication of a high-quality unmixing matrix. If the EDR is lower than 1000, then please refer to the suggestions indicated in the “Troubleshooting tips” section.

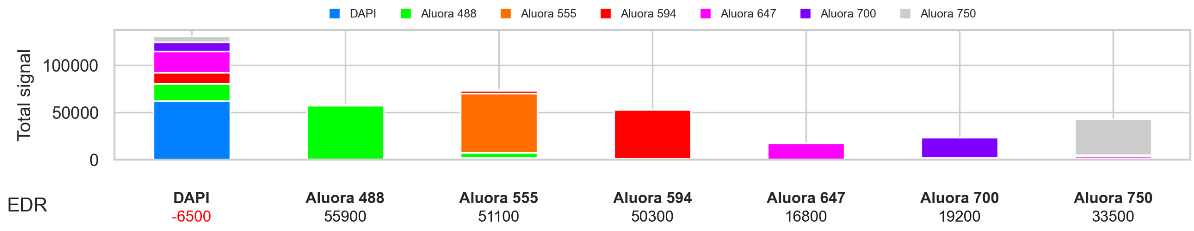


**Figure 6.** Quantitative assessment of the predicted unmixing quality via calculation of Effective Dynamic Range (EDR) values presented as Relative Fluorescent Units (RFU).



**For Research Use Only. Not for use in diagnostic procedures** © 2024 Thermo Fisher Scientific Inc. All rights reserved. All Unmixing Metrics Report trademarks are the property of Thermo Fisher Scientific and its subsidiaries unless otherwise specified.

#### Masks and extraction channels images

In Figure 7, the masks of positive pixels created from foreground and background channels are displayed. These created masks for each target (shown in the left most images) are those used for spectral extraction. Foreground and background channel are automatically set in the spectral extraction algorithm for each fluorophore, to generate the unmixing matrix.

Masks should select highly stained areas of the tissue with no artifacts. Inspect masks, foreground (FG) and background (BG) channels to ensure spectra were extracted from the correct region. The pattern in the mask image should closely resemble the staining pattern of the fluorophore as shown in the FG channel, whereas the BG channel should have low signal and be absent of staining patterns. Each extracted spectrum should have a unique signature. If the mask shows undesired pixels, refer to the Troubleshooting tips section to extract new spectra.

Note: Extracted spectra may differ from reference spectra. That could be attributed to differences on the tissues as well as targets.

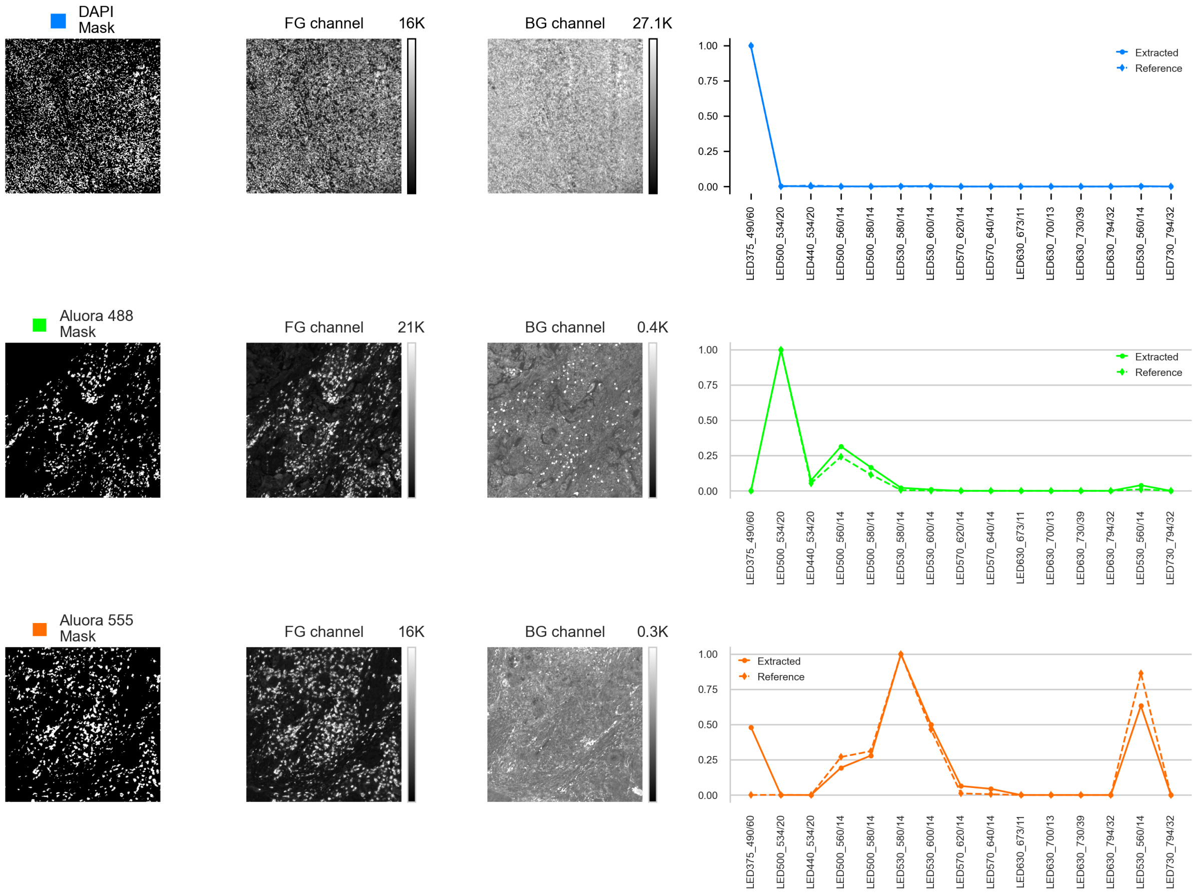




**For Research Use Only. Not for use in diagnostic procedures** © 2024 Thermo Fisher Scientific Inc. All rights reserved. All Unmixing Metrics Report trademarks are the property of Thermo Fisher Scientific and its subsidiaries unless otherwise specified.

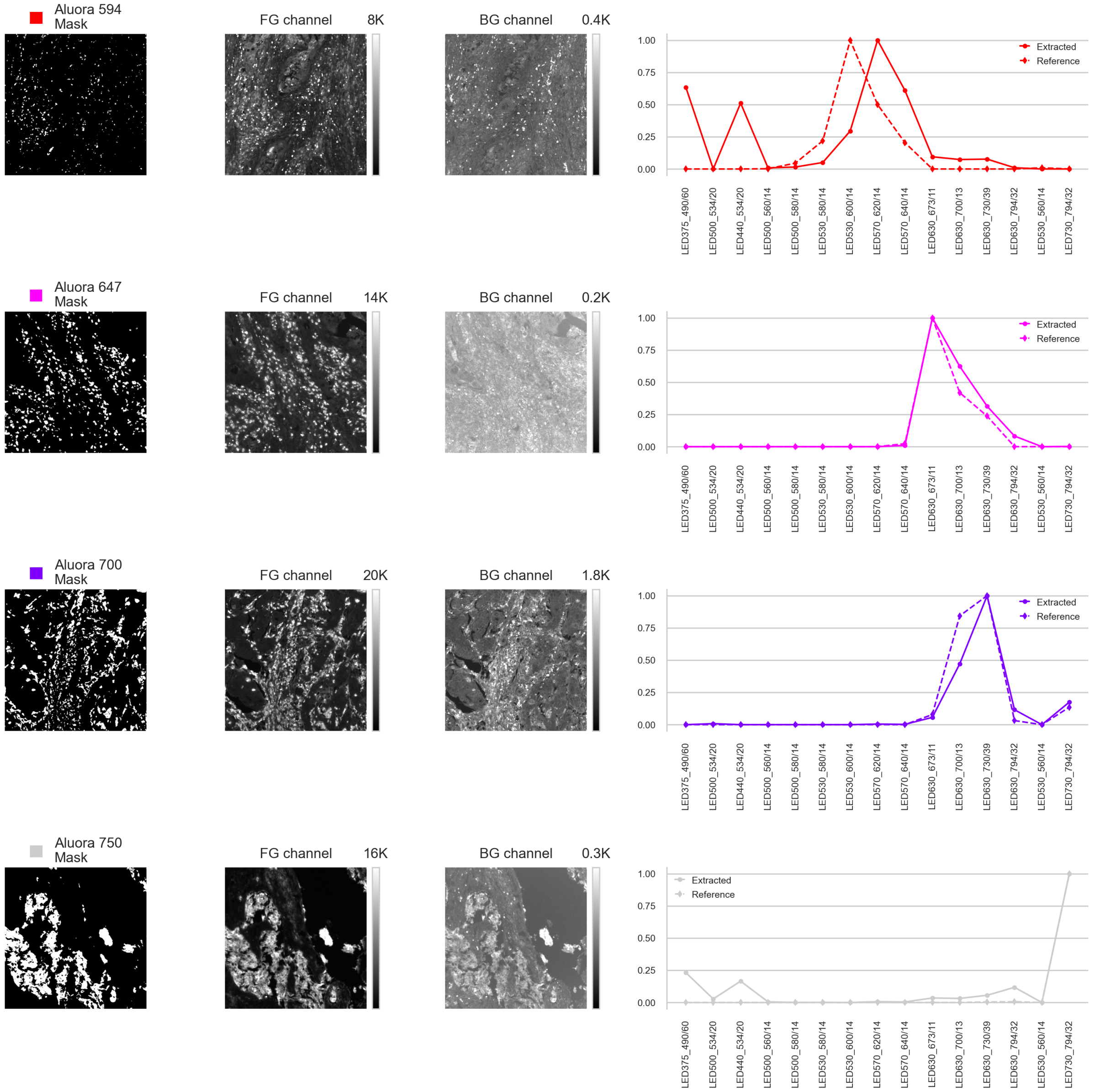


**Figure 7.** Mask of positive pixels created from the foreground and background channels to extract spectra.

#### The power of spectral unmixing

The presence of spectral overlap between fluorophores limits the number of individual biomarkers which can be identified by classic fluorescence microscopy. Spectral microscopy, on the other hand, allows for simultaneous detection of multiple biomarkers even in the presence of a high degree of spectral overlap. This is possible by applying spectral unmixing to the image data to resolve and separate signals from individual fluorophores. This is possible through the application of an unmixing algorithm, which calculates an unmixing matrix by measuring individual spectra from single-color control samples and applies the



**For Research Use Only. Not for use in diagnostic procedures** © 2024 Thermo Fisher Scientific Inc. All rights reserved. All Unmixing Metrics Report trademarks are the property of Thermo Fisher Scientific and its subsidiaries unless otherwise specified.

matrix to multiplexed samples labeled with the same fluorophores. The resulting output are unmixed images consisting of abundances of individual fluorophores at each pixel position across an image.

The images and metrics provided in this report enable the evaluation of the predicted unmixing performance based on single-color control input images. If the unmixing report indicates a high probability of error, consider the "Troubleshooting tips" in the following section.

#### Troubleshooting tips

Unmixing performance is heavily reliant on the quality of single-color controls (SCC) used to create the unmixing matrix. In turn, the quality of SCC is heavily reliant on antibody and labeling performance. If the metrics calculated above indicate the created unmixing matrix may not lead to high confidence unmixing, consider the following suggestions:

Regarding spectral extraction, the field of view captured for each single-color control is where the spectral signature is extracted. Extracted spectra are influenced by the specific marker and tissue.

- If the mask does not look like the fluorescent signal, ensure that the correct fluorophore was selected from the dropdown menu in the SCC menu to match with the sample.
- If the mask contains non-specific pixels from debris or other artifacts, re-image in a different region to ensure the field of view has sufficient desired staining to be detected while eliminating debris and artifacts.

Regarding quantitative metrics, FB and EDR are dependent on the signal intensity of all individual fluorophores.

- Be sure that images are not over or under exposed considering both primary and support channels. SCC sample images are automatically saved in the Target Files Location (defined in Settings). SCC sample images can be opened in Review tab for evaluation.
- Avoid using markers with an intensity difference of five times or more between any two adjacent channels that have spectral overlap. This can be observed in the EDR graph (Figure 6). If the target signal (primary channel) is less than the bleedthrough seen in that channel, the target fluorescence intensity is too weak and a different pair of markers should be assigned to those fluorophores.
- During labeling of your sample,
    - for the lowest expression targets, choose the brightest fluorophores (for example Aluora 647 or Alexa Fluor 647), and conversely,
    - for the highest expression targets use the less bright or dimmer fluorophores (for example Aluora 750 or Alexa Fluor 750).



**For Research Use Only. Not for use in diagnostic procedures** © 2024 Thermo Fisher Scientific Inc. All rights reserved. All Unmixing Metrics Report trademarks are the property of Thermo Fisher Scientific and its subsidiaries unless otherwise specified.
